# The combination of feedforward and feedback processing accounts for contextual effects in visual cortex

**DOI:** 10.1101/2022.05.27.493753

**Authors:** Serena Di Santo, Mario Dipoppa, Andreas Keller, Morgane Roth, Massimo Scanziani, Kenneth D. Miller

## Abstract

Sensory systems must combine local features with context to infer meaning. Accordingly, context profoundly influences neural responses. We developed a unified circuit model demonstrating how feedforward and feedback inputs are integrated to produce three forms of contextual effects in mouse primary visual cortex (V1). First, reanalyzing existing data, we discovered that increasing stimulus size only weakly increases the area of V1 neural response, conflicting with previous models of surround suppression (SS). Second, through modeling, we found that, in Layer 2/3, (1) SS and its contrast dependence are largely inherited from Layer 4; (2) Inverse responses (IR) – size-tuned responses to a gray “hole” in a full-field grating – are driven by feedback connections provided they are sufficiently wide; (3) Cross-orientation surround facilitation is induced by the summation of feedback input driving IR with the feedforward-driven classical center response. The model accounts for many previous findings and makes multiple testable predictions.

**Highlights:**

- One model explains three different types of contextual modulation
- The widths of spatial response patterns grow much more slowly than stimulus size.
- Inverse responses depend on the geometry of feedback response fields and projections
- Summation of classical and inverse response accounts for surround facilitation.

## 1 Introduction

When an edge or other feature appears in a visual scene, its meaning – is it an object boundary? an element of texture? a shadow? – must be inferred from the larger scene in which it is embedded. More generally, sensory systems must combine local features with context to infer meaning. Accordingly, context profoundly influences our perception, as is made strikingly clear by visual illusions. For example, the perceived contrast or luminance of a patch of the visual scene can be radically altered by whether the context makes the patch appear a small part of a larger object or an independent object, or in light or in shadow (Fig. 1a-c).

**Figure 1:**
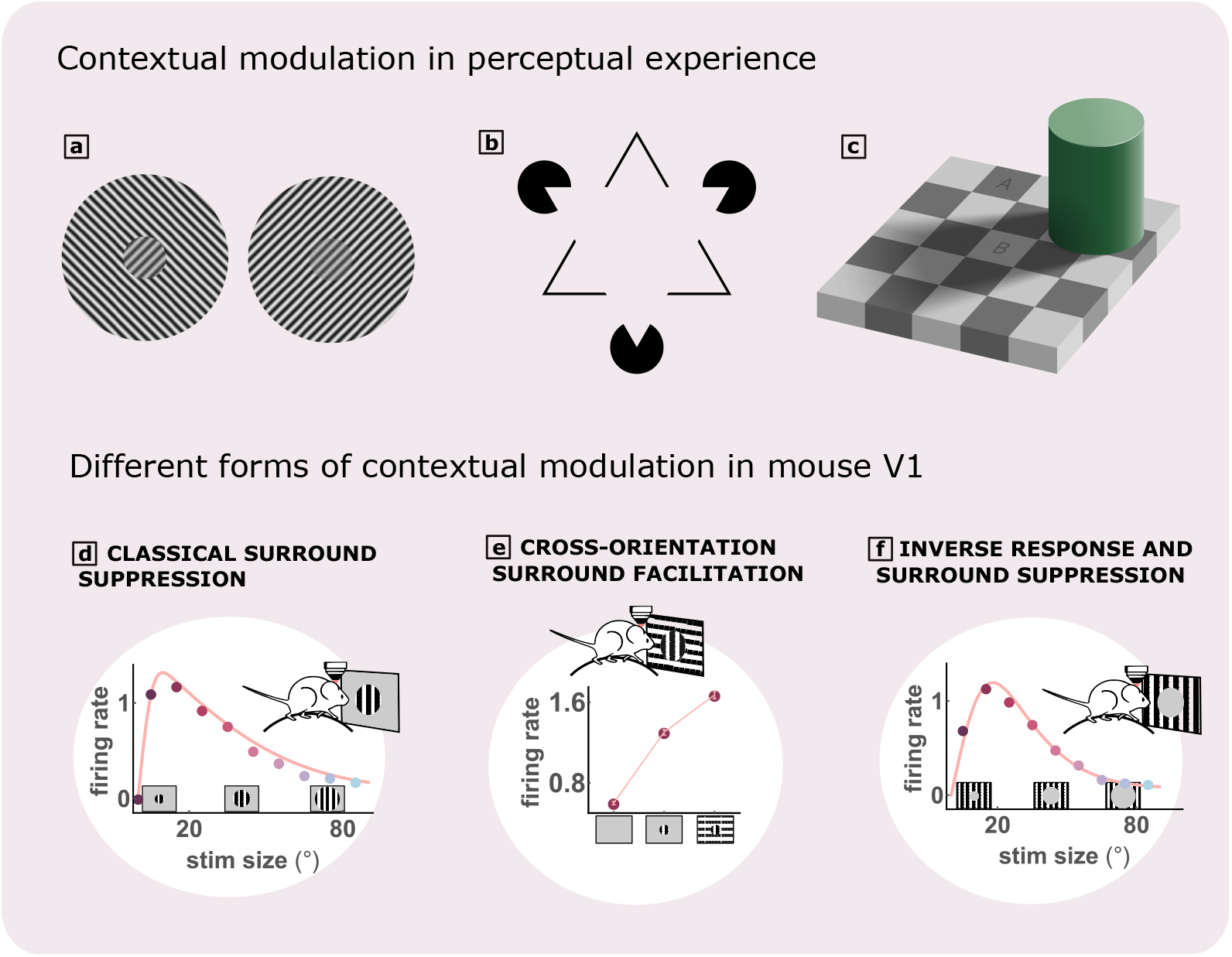
Different forms of contextual modulation. **a**. The grating patches in the centers are identical but are perceived as more or less salient due to the distinct surround. **b**. Kanizsa triangle illusion: a nonexistent white triangle in the center appears to occlude the shapes around it **c**. Checker shadow illusion: the areas labeled A and B have identical brightness but A appears darker than B due to brain’s inference that B is in shadow. **d** Classical size tuning and surround suppression: responses decrease when the stimulus size is increased beyond the CRF size; data (circles) from pyramidal cells studied by Keller et al. (2020b), line is a fit to the data. Here and in what follows, firing rates are in arbitrary units. **e**. Cross-orientation surround facilitation: responses to a patch of drifting grating are enhanced by addition of an orthogonally-oriented surrounding grating. Data replotted with permission from (Keller et al., 2020a). **f**. Inverse response: response to a ‘hole’ or inverse stimulus in the CRF, *i*.*e*. a patch with zero contrast on a full field drifting grating. Note similarity of inverse size tuning to classical size tuning. Data (from Keller et al., 2020b) and fit as in (a).

Context strongly impacts not only our perception but also neuronal responses. Some of the best-studied examples involve the effects of context on neurons in the primary visual cortex (V1) (*e*.*g*., Angelucci et al., 2017; Dipoppa et al., 2018; Keller et al., 2020a,b; Ziemba et al., 2018; Zipser et al., 1996). V1 neurons are primarily driven by stimuli only within a small region of visual space, known as the neuron’s classical receptive field (CRF). Yet the visual stimulus in the surrounding region (the neuron’s “surround”) can substantially influence the cell’s response. This contextual influence is likely to be critical to the parsing of objects and formation of a coherent perceptual representation of the visual scene (Field et al., 1993; Kirch-berger et al., 2021, 2023a; Pak et al., 2020; Roelfsema, 2006).

Here we develop a simple theoretical framework to address the question: how do the properties of feedforward, feedback and lateral input currents contribute to contextual modulation in V1? We arrive at a unified mechanistic picture of how these different inputs are integrated to produce diverse forms of contextual modulation. This gives a coherent, experimentally testable theory of V1 computation that can serve as a more general framework for studying how cortical circuits integrate context into sensory processing.

Three exemplary paradigms have been widely used to study contextual modulation in V1 (Fig. 1d-f).

- The response of a cell whose CRF is centered on a stimulus decreases when a similar stimulus is presented in the surround. This *surround suppression* (Allman et al., 1985; Angelucci et al., 2017; Dipoppa et al., 2018; Hubel and Wiesel, 1965; Keller et al., 2020a; Sceniak et al., 2001; Shushruth et al., 2009; Spillmann et al., 2015) may represent a discounting of visual input that can be well predicted from other parts of the visual scene, thus allowing an efficient neural representation (Barlow, 2001, 1972; Coen-Cagli et al., 2015).
- In contrast, addition of a stimulus surround that is very distinct from the CRF stimulus can enhance a neuron’s response (Cavanaugh et al., 2002; Field et al., 2010; Keller et al., 2020a; Paffen et al., 2005; Polat et al., 1998; Self et al., 2014; Xing and Heeger, 2001). This *surround facilitation* can enhance the perceptual salience of discontinuities in the visual world, which likely separate different objects (Craft et al., 2007; Lamme, 1995; Rubin, 2001). Given an oriented grating as CRF stimulus, this facilitation can be evoked by an orthogonally oriented stimulus in the surround (*cross-orientation surround facilitation*).
- Surrounding context that indicates occluded structure in the center (Fig. 1b) can evoke illusory perception of that structure (Kanizsa, 1976; Pak et al., 2020). While illusory contour responses in V1 are rare (Peterhans and von der Heydt, 1989; Shin et al., 2023; von der Heydt, 1987; but see Grosof et al., 1993), a robust example of such context-induced neural responses in V1 is the “inverse response”: When a large drifting grating is presented, and a uniform patch of mean luminance (a ‘hole’ in the grating) of varying sizes is centered on the CRF, neurons show responses and size tuning to the hole very much like those to drifting gratings of various sizes (Jones et al., 2001; Keller et al., 2020b; Kirchberger et al., 2023b; Rossi et al., 2001). We will refer to the ‘hole’ stimulus as an “inverse stimulus”, and response suppression for large inverse stimuli as *inverse surround suppression* (although this is something of a misnomer, in that the suppression is induced by expanding the hole, leaving less visual contrast in the surround). In contrast, we will refer to a drifting grating stimulus as a “classical stimulus”. Inverse responses have substantially longer latencies than classical responses (Keller et al., 2020b; Kirchberger et al., 2023a; Rossi et al., 2001), and are substantially reduced if higher visual areas (HVAs) are optogenetically suppressed (Keller et al., 2020b), suggesting that feedback connections play a key role in inverse responses.

Here we develop a unified model of these three contextual effects in layers 2/3 (L2/3) of mouse V1. We consider the four cortical cell types that have been prominently studied in recent years: excitatory or pyramidal (Pyr) cells, and parvalbumin-expressing (PV), somatostatin-expressing (SOM) and vaso-active-intestinal-peptide-expressing (VIP) inhibitory interneurons. We model the cellular response patterns across 2D continuous space, which allows us to determine how the geometry of spatial connections contributes to contextual modulation. We incorporate experimental constraints on the spatial extents of connections (Billeh et al., 2020; Li and Young, 2021; Marques et al., 2018; Rossi et al., 2020) and data from Keller et al. (2020b) on the response profiles across space of each cell type and of inputs from L4 and from an HVA, area LM, to each stimulus type. We focus on L2/3 both because of the availability of this data and because inverse responses, which depend on feedback from HVAs, are found in L2/3 but not in L4 (Keller et al., 2020b), suggesting that L2/3 is a key site of integration of top-down with bottom-up input.

A number of theoretical works, at different levels of abstraction, have been developed previously to account for various aspects of contextual modulation. Phenomenological models place it within a broad range of nonlinear response properties described as “divisive normalization” (*e*.*g*., Carandini and Heeger, 2012; Carandini et al., 2005; Coen-Cagli et al., 2015). Circuit models (*e*.*g*., Dipoppa et al., 2018; Keller et al., 2020a; Li and Young, 2021; Mossing et al., 2021; Obeid and Miller, 2021b; Rubin et al., 2015; Schwabe et al., 2006, 2010; Shushruth et al., 2012b) have not previously addressed inverse tuning or the roles of top-down feedback or, in many cases, multiple types of inhibitory cell. In addition, our approach differs in being based on analysis of experimental data on the 2D spatial structure of activity (as also was Dipoppa et al. (2018)) and connectivity, and developing an analytical framework that gives insight into precisely how interactions across space yield contextual modulation. Our analysis of experimental data produces the surprising finding that, with increasing stimulus size, activity patterns show weak or no increase in spatial width, which forces revision of most previous models of contextual modulation.

To isolate basic principles, we focus on an abstract, “minimal” nonlinear model of a single recurrently-connected cell type representing the joint activity of Pyr and PV cells in L2/3, along with external input from SOM cells, that can be fully solved analytically. We then show that a full model of the four cell types reproduces all contextual modulation phenomena via the same mechanisms uncovered in this minimal model (Supplementary SS15–SS17). We focus on qualitatively explaining response properties and understanding their underlying mechanisms. This enables multiple insights (summarized in Table 1), including the spatial structure of top-down projections required for inverse responses to arise, how the combination of classical and inverse responses can produce cross-orientation facilitation, and that SOM should make opposite contributions to classical vs. inverse surround suppression.

**Table 1:** All observed phenomena that are included in or generated by the model. Light brown rows indicate known effects reproduced by the model, orange ones indicate phenomena or mechanisms that are highlighted here for the first time but are compatible with work in the literature and light red ones indicates testable predictions. The last four predictions follow from the model prediction that cross-orientation surround facilitation is, at least in part, simply the addition of the inverse response to the center response.

| Observations and predictions | Fig. | Ref |
| --- | --- | --- |
| classical and inverse size-dependent spatial responses, cross-orientation surround facilitation | 3c-d, 5b-c, 7e-f, SF31 | (Keller et al., 2020a,b),... |
| the rate field width grows much more slowly than the stimulus size. | 2e-h, 4e-h | Compatible with (Dipoppa et al., 2018) |
| L2/3 responses are stabilized by PV inhibition | SF15, SF35 | (Denève and Machens, 2016; Sanzeni et al., 2020).. |
| by activating PV inhibition, classical surround suppression decreases. | SF15 | (Nienborg et al., 2013a) |
| in mouse L2/3 of V1, suppression is largely inherited from the input, rather than due to recruiting more lateral PV inhibitory activity. | 3e-k | Compatible with (Mossing et al., 2021) |
| classical preferred-size-contrast-dependence is largely inherited from feedforward and feedback layers. | SF23b-c | Compatible with (Mossing et al., 2021) |
| classical preferred-size-contrast-dependence is supported by large VIP/small SOM response for low contrasts. | SF23h | (Mossing et al., 2021) |
| SOM cells can generate ring profiles in the rate field of Pyr and PV cells. | 3m | Compatible with (Dipoppa et al., 2018) |
| hyperpolarizing SOM reduces classical surround suppression. | 3n, SF36 | (Adesnik et al., 2012b) |
| inverse response (but not classical response) is substantially reduced when input from HVAs is reduced. | 5e, SF26, SF37 | (Keller et al., 2020b; Vangeneugden et al., 2019) |
| hyperpolarizing SOM increases inverse surround suppression. | 5d, SF36 | (prediction) |
| inverse responses increase with contrast. | 5f, SF27, SF38 | (Keller et al., 2020b) |
| inverse response requires wide enough feedback projections. | 6a-d, SF39a | Compatible with (Marques et al., 2018) |
| inverse surround suppression is given by widening of HVAs pattern of activity | 6e-m, SF39b-c | Compatible with (Keller et al., 2020b) |
| when optogenetically silencing a fraction of PV neurons, surround facilitation increases. | 7h | (prediction) |
| cross-orientation surround facilitation should be stronger for cells that have orthogonal classical and inverse preferred orientation, compared to cells with the same preferred orientation for classical and inverse stimuli. | 7d-e | (prediction) |
| Pyr and PV cells in L2/3 centered on the stimulus and with preferred orientation orthogonal to the center stimulus show significant response to the cross stimulus. | 7e | (prediction) |
| optogenetically silencing HVAs during cross stimulus presentation reduces surround facilitation. | 7g | (prediction) |
| hyperpolarizing SOM cells during cross stimulus presentation reduces surround facilitation. | 7g | Compatible with (Keller et al., 2020a) |

## 2 Results

### 2.1 A spatial schematic description preserving anatomical length-scales

To take into account input from neurons with CRFs distributed across visual space (Dipoppa et al., 2018; Keller et al., 2020b), we require knowledge of two factors: i) how the mean effective strength of synaptic connections depends on the distance in visual space between the CRF centers of the pre- and post-synaptic cells and ii) how the responses of cells to a stimulus depend on the distance between the cell’s CRF center and the stimulus center. For dependence (i), we assume that synaptic strengths decrease with distance between two neurons as a Gaussian function, with length scales taken from experimental measurements (Billeh et al., 2020; Keller et al., 2020b; Li and Young, 2021; Marques et al., 2018; Rossi et al., 2020) (see Supplementary SS1). We take this approach, rather than trying to infer connections that lead to responses that fit the data (*e*.*g*., Dipoppa et al., 2018), to more tightly constrain our models. We show that our results are robust when considering very large relative errors in the experimental estimates in a biologically realistic model (Supplementary SS15, SS16). For dependence (ii), we use the dataset of Keller et al. (2020b) and a smoothing method introduced in Dipoppa et al. (2018) to reconstruct the *firing-rate field* (firing rate vs. CRF spatial position) of Pyr, PV, SOM and VIP cells in L2/3 of V1, as well as Pyr cells in L4 of V1 and in the higher visual area LM (Fig. 2, Supplementary SS2 B). We will also describe these firing-rate fields as Gaussian (or linear combinations of Gaussian) functions centered on the stimulus center.

**Figure 2:**
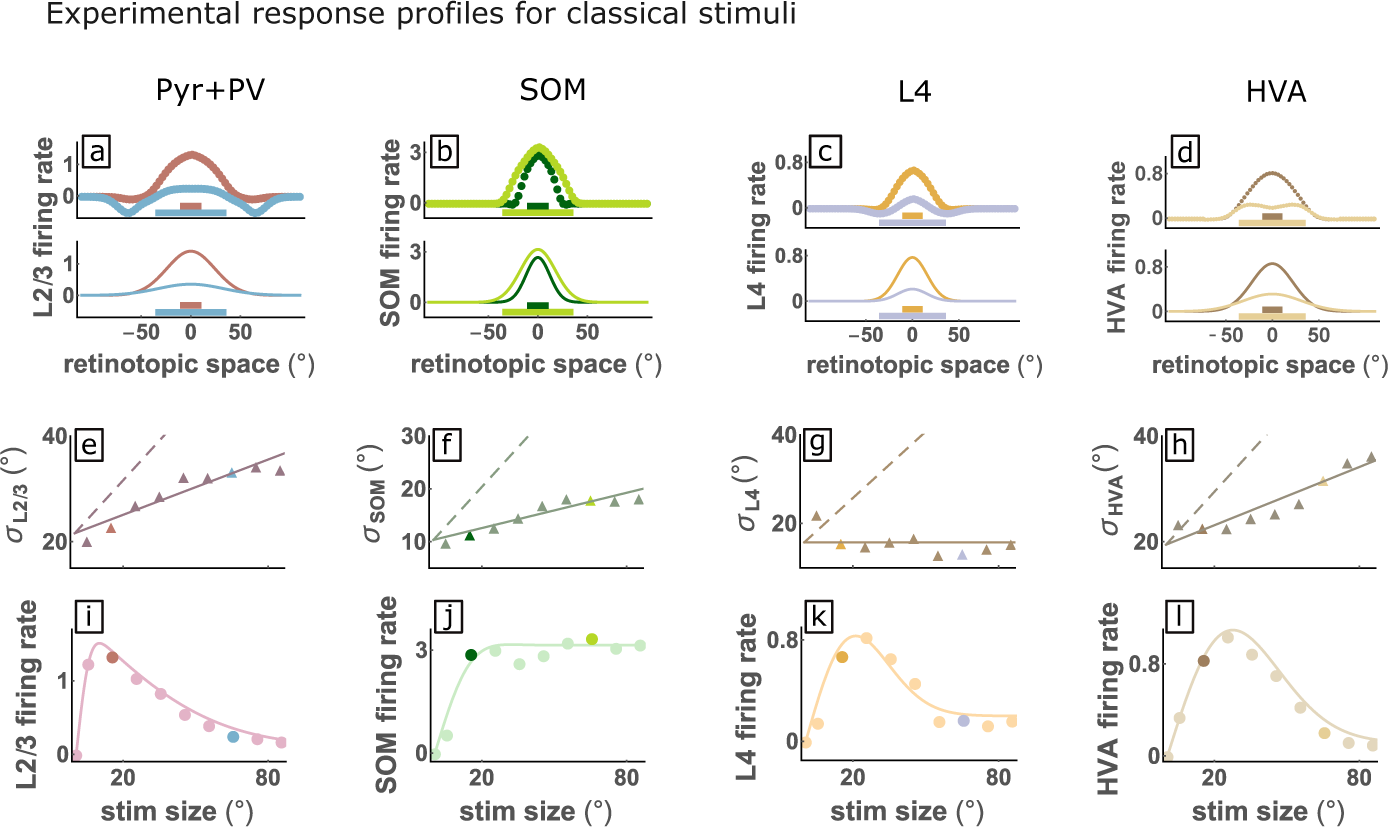
The widths of spatial response patterns grow much more slowly than stimulus size. Pyr+Pv, SOM, L4 and HVAs spatial responses, calculated from experimental data (Supplementary SS2 B). **a-d**. experimental rate field (top) and best Gaussian fit (bottom), for stimulus sizes *s* = *{*15^*°*^, 65^*°*^*}*; x axis represents distance of RF from stimulus center. Colored horizontal bars represent size of the corresponding stimuli. **e-h**. Full triangles are the widths of the best Gaussian fits for each stimulus size, the full line is a linear fit and the dashed line (slope 1/2) represents the case in which the rate field width grows at the same rate as the stimulus radius (i.e. 1/2 of the stimulus size). **i-l**. Response of cells at stimulus center vs. stimulus size (size tuning curve). Full circles, data points; full line, fits; dots in different colors highlight responses for the two stimuli in **a-d**.)

#### 2.1.1 The widths of spatial response patterns grow much more slowly than the stimulus size

We estimated rate fields elicited by classical stimuli of varying sizes from the recordings in (Keller et al., 2020b) (Fig. 2; see also Supplementary SS2 B). We combine Pyr and PV cells into a single joint population, because of the similarity of their responses and for better comparison with the model to be developed in Section 2.2. The rate fields for Pyr+PV and SOM cells and for the inputs, L4 and area LM (HVAs), have similar widths as functions of retinotopic space for small and large stimulus size (Fig. 2a-d top) (rate fields for all cell types and all stimulus sizes, as computed by two different methods, are shown in Supplementary Fig. SF2). This is quantified by fitting a Gaussian function to each spatial profile (Fig. 2a-d bottom), and comparing (Fig. 2e-h) the width (standard deviation; points and solid lines) to the width expected if the rate field expanded in size equally to the stimulus (dashed lines) (shown for all cell types and both methods of calculating spatial rate fields in Supplementary Fig. SF6). The rate fields expand far less than the expected rate, and not at all in the case of L4. Finally, the amplitude of the Gaussian fits vs. stimulus size gives the size tuning curve of the neurons that are centered on the stimulus (Fig. 2i-l) (shown for all cell types in Supplementary Fig. SF7).

The assumption that rate field width grows to match the stimulus radius is at the core of existing theoretical models of classical surround suppression (Angelucci et al., 2017; Dipoppa et al., 2018; Li and Young, 2021; Obeid and Miller, 2021a; Rubin et al., 2015), in which larger stimuli recruit more wide-ranging lateral activation, and these activated lateral cells have a suppressive influence on the center. The failure of this assumption is, to the best of our knowledge, systematically discussed here for the first time, although it can also be seen in the plots of Dipoppa et al. (2018) (their Supplementary Figure 11).

In what follows, we first develop a minimal model for ordinary surround suppression (Allman et al., 1985; Angelucci et al., 2017; Hubel and Wiesel, 1965; Sceniak et al., 2001; Shushruth et al., 2009; Spillmann et al., 2015). We then apply this model to understand inverse response (Keller et al., 2020b). Finally, we extend the model to understand cross-orientation surround facilitation (Keller et al., 2020a).

### 2.2 A minimal model for contextual modulation

We develop a minimal model to address the question: how do feedforward, feedback and lateral input currents contribute to contextual modulation in L2/3? In this model, L2/3 consists of only a single recurrently connected cell type, representing the combination of Pyr and PV cells. This is based on observations that PV responses track those of Pyr cells in multiple contexts and over a wide dynamic range of inputs (Haider et al., 2006; Keller et al., 2020b; Mossing et al., 2021; Palmigiano et al., 2020), thereby dynamically balancing (preventing an excess of) excitation and functioning as a stabilizer for the network (Adesnik, 2017; Ahmadian and Miller, 2021; Bos et al., 2020; Den`eve and Machens, 2016; Dipoppa et al., 2018; Isaacson and Scanziani, 2011; Kato et al., 2017; Okun and Lampl, 2008; Ozeki et al., 2009; Palmigiano et al., 2020; Renart et al., 2010; Sanzeni et al., 2020). In contrast, VIP and particularly SOM cell responses generally differ substantially from those of Pyr and PV cells (Dipoppa et al., 2018; Keller et al., 2020a,b; Mossing et al., 2021; Palmigiano et al., 2020). We treat SOM cells as a static external input, responding across space as measured experimentally, and ignore VIP cells, which act primarily on SOM cells (Billeh et al., 2020; Campagnola et al., 2021; Pfeffer et al., 2013). We also take as external inputs the measured responses across space of layer 4 (L4) and Lateromedial area (area LM) excitatory neurons. We treat LM as a static input, rather than as part of the recurrent circuit, due to the limited information available about non-L2/3 inputs to LM (see Discussion).

We assume neurons have a supralinear input/output function (taken to be rectified quadratic, see STAR Methods), describing the steady-state firing rate induced by a given net input to the neuron (Priebe and Ferster, 2008). Such an expansive input/output function is expected when firing activity is driven by input fluctuations rather than by the mean input (Hansel and van Vreeswijk, 2002; Miller and Troyer, 2002), and has been shown (Ahmadian and Miller, 2021; Rubin et al., 2015) to yield network behavior that reproduces a variety of nonlinear visual cortical behaviors, including contrast-dependent surround suppression, that have often been summarized phenomenologically as “divisive normalization” (Carandini and Heeger, 2012). The use of these power-law input/output functions, combined with our assumptions that connectivity and firing-rate fields are described by Gaussian functions, allows us, building on previous theoretical work (Persi et al., 2011), to develop an approximate analytic solution to the one-population model, which yields deeper insight into the mechanisms driving model behavior.

A crucial and novel aspect of the model is to include inputs from HVAs. A difficulty in doing so is that cells in L2/3 are targeted by all of the HVAs (Billeh et al., 2020; Keller et al., 2020b; Siegle et al., 2021), but we have recordings of the V1 boutons from only one HVA, area LM (Keller et al., 2020b). Extended Data Fig.9 in Keller et al. (2020b) shows that optogenetic silencing of different HVAs generates quantitatively different reductions of classical versus inverse responses. We assume that the HVAs considered altogether have a rate field profile proportional to that recorded in LM, but with proportionality constants that can differ between the classical and inverse stimulus conditions. We infer these constants based on the amplitudes of inverse vs. classical response in L2/3 (see Supplementary SS5). Keller et al. (2020b) suggested that LM cells projecting to a given point in L2/3 from the same retinotopic location or from peripheral locations (aligned vs offset cells) had qualitatively different RF properties, but our analysis finds no such difference (Supplementary SS3), so we treat HVA cells as being homogeneous across space.

### 2.3 Classical surround suppression in the minimal model

In our minimal model of classical size tuning (Fig. 3), inputs to L2/3 are given by the convolution of the rate fields of L4 and HVAs (and SOM when specified), which were derived from the data (Fig. 2), with the corresponding Gaussian connectivity (Fig. 3b). The recurrent rate fields resulting from the approximate analytical solution and the simulation of the minimal model agree well (Fig. 3c-d and Supplementary SS6), so in what follows we rely on the analytical formulas to efficiently compute model results. The model’s free parameters are the amplitudes and widths of the feedforward, feedback and recurrent connection strengths. Since we consider the recurrent population to be a joint population of Pyr and PV cells, the recurrent connection strength *W*_0_ can be either positive or negative, depending on whether Pyr or PV dominates. We find that L2/3 responses are unstable if *W*_0_ is too strongly positive, while making *W*_0_ too strongly negative decreases surround modulation (Supplementary SS7 B), consistent with effects of optogenetic activation of PV cells (Nienborg et al., 2013b). We set *W*_0_ to a moderate negative value. The feedforward and feedback connection strengths were then set to reproduce the amplitude of the response in L2/3 (Supplementary SS7 C).

**Figure 3:**
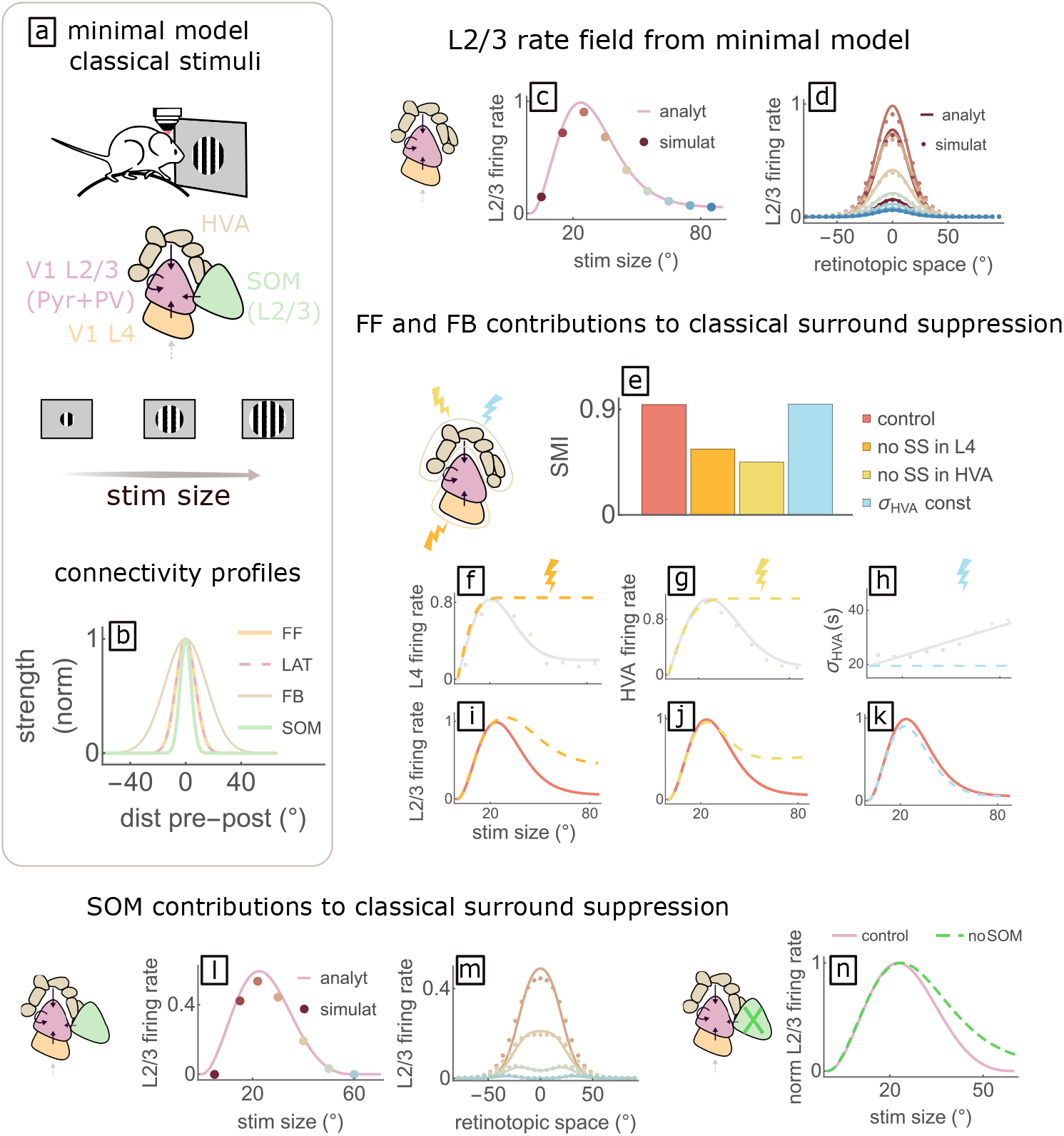
Feedforward, feedback and lateral SOM inputs shape classical surround suppression. **a**. Top to bottom, sketches of: experimental setup with classical stimuli; minimal model with one recurrent cell type in V1 L2/3 receiving feedforward (FF) input from L4, feedback (FB) input from HVAs, lateral input from its own type (LAT) and from SOM cells; classical size tuning stimuli. **b**. Strength of connections vs. CRF distance between pre- and post-synaptic neurons, for the four types of connections. **c**. Size-tuning curve and **d**. rate field for the recurrent layer of the model. Here and in **l**,**m**, color code of the full circles indicates stimulus size; full circles represent simulations, full lines represent analytical results. **e-k**. Counterfactual modifications of the inputs. **e**. Bar plot representing the SMI (see main text; = 0, complete suppression; = 1, no suppression) in the control condition, and when we eliminate either: surround suppression (SS) in L4, surround suppression in HVAs, or growth of the width of the HVAs rate field with stimulus size (*σ*_*HV A*_ const). **f-h**. Illustration of the counterfactual modifications (dashed, same color code as bar plot) compared to the empirical values (full lines and symbols as in Fig. 2). **i-k**. Effects of the counterfactual modifications on L2/3 size tuning curve (same color code as above). **l-n**. Effects of adding SOM input. **l**,**m** Same as **c**,**d** but with SOM input. For large stimuli, the rate field convexity in the center changes. Compare with experimental profiles in Supplementary SS6 A. **n**. Size tuning curve of recurrent layer with (pink; same as **l**) and without (dashed green; same as **c**) SOM input.

#### 2.3.1 Feedforward, feedback and lateral inhibitory input shape classical surround suppression

Many experiments show that feedforward, horizontal, and feedback connections all contribute to V1 classical surround suppression (*e*.*g*., Angelucci et al., 2017 and references therein): the minimal model allows us to analyze their interplay. We first analyze the contributions of feedforward (L4) and feedback (HVAs) inputs. In the next section, we consider addition of SOM input.

In Supplementary SS8, using our analytical solution, we show that surround suppression in the recurrent layer can be produced by either or both of: i) surround suppression in the input population and ii) broadening of the input rate field as stimulus size is increased. In case (i), L2/3 surround suppression is inherited from the input (although it may be enhanced by L2/3 network mechanisms that increase response differences relative to input differences). In case (ii), L2/3 surround suppression arises from increased lateral activity driving greater center response suppression. Since the scaling of the widths is weak for LM and absent for L4 (Fig. 2) we expect the second mechanism to contribute only marginally to L2/3 surround suppression.

To test this hypothesis we consider the effects of (counterfactually) eliminating surround suppression in L4 and HVAs (Figure 3e-g,i-j) and of eliminating the growth of the width of the HVAs rate field with increasing stimulus size (Figure 3h,k; note the width of the L4 rate field does not increase with stimulus size). In each condition, we calculate a *Surround Modulation Index (SMI), SMI* = 1*−r*_Large_*/r*_Pref_, where *r*_Large_ and *r*_Pref_ are the firing rate of the centered cells at the largest stimulus size (*s* = 85^*°*^) and the preferred size, respectively (if *r*_*large*_ = *r*_*pref*_, then *SMI* = 0). Consistent with our hypothesis, eliminating surround suppression in the inputs strongly reduces surround suppression in L2/3, while eliminating growth of the HVAs rate field width does not alter surround suppression in L2/3.

In Supplementary SS9 we compare the recurrent rate field obtained from a supralinear vs. a linear transfer function and show that the former supports stronger surround suppression, presumably because the nonlinearity increases the gap between responses to stronger vs. weaker stimuli.

Surround suppression is weak for low-contrast stimuli and increases in strength, while optimal size shrinks, with increasing stimulus contrast (*e*.*g*., Mossing et al., 2021). In Supplementary SS8 D we analyze the contrast dependence of size tuning in the model. We find that a supralinear one-population model with Gaussian input whose rate field width increases with stimulus size can generate the observed contrast-dependent size tuning, i.e. higher contrasts produce larger surround modulation and smaller preferred sizes, in line with the results in the two-population E/I model of Rubin et al. (2015). However, the contrast-dependence in our one-population model is parameter dependent, and in particular does not occur with the connectivity length scales we are using, which are matched with those observed experimentally in mouse L2/3. Instead, and in agreement with (Mossing et al., 2021), our model suggests that in mouse L2/3 contrast-dependence is largely inherited from its feedforward layers. In Supplementary SS10, we examine this contrast dependence further, and find that additional contributing factors may be i) scaling of rate field width in the feedback input and ii) large VIP/small SOM response for low contrasts (in agreement with Mossing et al. (2021)).

In summary, our model suggests that both surround suppression and its contrast dependence in L2/3 are largely inherited from its inputs, although the supralinear input/output function also strengthens surround suppression.

#### 2.3.2 SOM cells enhance classical surround suppression and generate more complex spatial profiles

The rate field of SOM neurons in L2/3 lacks surround suppression (Fig. 2j), in agreement with previous results (Adesnik et al., 2012a) (see Supplementary SS4 A). We compare the size tuning curve of the model recurrent population in the presence or absence of SOM input, as in Adesnik et al. (2012b). Since SOM cells prefer larger stimulus sizes, they inhibit L2/3 the most for larger stimuli, thus enhancing L2/3 classical surround suppression (Fig. 3l,n). This effect is robust against changes in the free parameters, i.e. the connection strengths. However, when the effective strength of SOM projections onto the Pyr+PV population is large enough, the model Pyr+PV rate field has a “ring” shape for large stimulus sizes (Fig. 3m), which is well captured by the analytic solution of the minimal model. This ring shape for large stimulus sizes has been reported in Dipoppa et al. (2018); their data is replotted for the joint population of Pyr and PV in Supplementary SS6 A).

The absence of this feature in the dataset of (Keller et al., 2020b) (Fig. 2i) could conceivably be explained if those animals had a smaller effective strength of SOM projections than in (Dipoppa et al., 2018). From a geometric point of view, the SOM input field is a (relatively narrow) Gaussian-shaped bump (*e*.*g*. see SOM rate field in Fig. 2), so it most strongly inhibits central Pyr+PV cells. For large stimuli its amplitude becomes larger than that of other inputs, generating a concave Pyr+PV rate field. The excitatory input from HVAs can also contribute to this effect, since for large stimuli it also exhibits a ring profile (Fig. 2l, Supplementary SS4 A).

In conclusion, our model predicts that classical surround suppression in L2/3 is largely inherited from surround suppression of its feedforward (L4) and feedback (HVAs) inputs (consistent with the experimental literature, Angelucci et al. 2017; Vangeneugden et al. 2019), marginally enhanced by lateral (PV) inhibition progressively recruited as the stimulus size increases (Obeid and Miller, 2021a; Rubin et al., 2015), enhanced by SOM lateral input (Adesnik et al., 2012a) and partially supported by the supralinear input/output function (Hansel and van Vreeswijk, 2002; Miller and Troyer, 2002).

### 2.4 Inverse response and size tuning properties of L2/3 are shaped by the feedback input

In the previous Sections we developed the minimal model, with most parameters set from experimental data. Here we use the same model, with no adjustment of parameters except the increased amplitude of HVAs input for inverse vs. classical stimuli (described in Section 2.2, further detailed in Supplementary SS5), to understand responses to inverse stimuli of varying sizes (*i*.*e*., hole diameters).

In experiments, letting position 0° be the stimulus center, L2/3 cells with CRF center near zero (“aligned cells”) respond well to inverse stimuli, but the aligned inputs from L4 and HVAs have very low firing rates (Fig. 4, Supplementary SS4 and SS6). This led the authors of Keller et al. (2020b) to conclude that the inverse tuning properties of L2/3 are not directly inherited from its retinotopically aligned inputs. Aligned cells in L2/3 show size tuning to inverse stimuli very similar to classical size tuning (Fig. 4i, full circles, compare with Fig. 2i). However, the spatial representation of inverse stimuli is qualitatively different from the spatial representation of classical ones: for moderate-sized (35°) to large stimuli, the L2/3 rate field for inverse stimuli has a local minimum at 0°, so that the peak response is at a characteristic distance from the center (Fig. 4a, Supplementary Fig. SF4). This is not true for classical stimuli (Supplementary Fig. SF2), although, as we noted previously, it can occur for large enough stimuli both in the model (Fig. 3m) and in experiments (Dipoppa et al., 2018). Similarly to the classical stimulus condition, the widths of the inverse rate fields scale weakly with the stimulus size (Fig. 4e-h, Supplementary SS4 A). To the best of our knowledge, these latter two features of the inverse rate fields are identified here for the first time.

**Figure 4:**
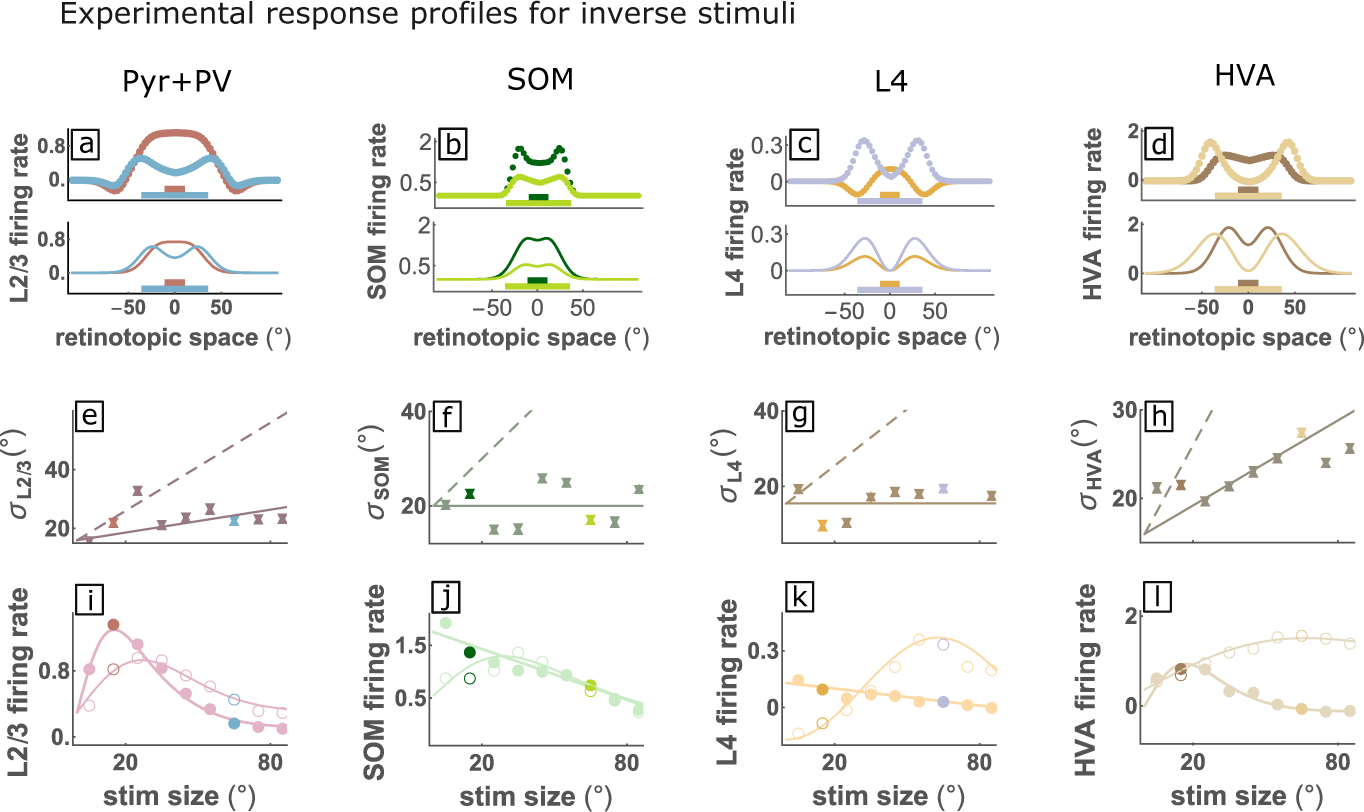
Inverse response and size tuning properties of V1 and HVAs. Same as Fig. 2 but for inverse stimuli. **a-d**. Experimental rate field (top) and best difference of Gaussians fit (bottom) for two stimulus sizes. The bottom bars represent the size of the corresponding inverse stimuli. **e-h**. Up and Down full triangles are the widths of the positive and negative Gaussian function in the best difference-of-Gaussian fits to the rate fields. Differences in widths of the two Gaussians are small, *i*.*e*. up and down triangles are almost overlapping. Full line: linear fit, dashed line: fit if rate field width (of the positive Gaussian) grew as the radius of the stimulus. **i-l**. Inverse size tuning curves. Full (empty) circles are experimental responses of cells centered on (with an offset of 30^*°*^ with respect to) the stimulus center, vs. inverse stimulus size. Full lines are fits.

To leverage our analytical framework, we parameterize the input rate fields with difference-of-Gaussian functions (Fig. 4a-d bottom) whose parameters vary continuously with stimulus size (see STAR Methods and Supplementary SS4, SS4 A). In doing so, we trade off some details of the spatial profiles of the responses to gain interpretability. In particular, to take advantage of the minimal model, we adopt a compact analytical expression for the L4 rate field, that limits the accuracy of the fit for small stimuli (Fig. 4c). However this should be irrelevant for recovering inverse response, because inverse response is mainly dependent on input from HVAs (Keller et al., 2020b). In Supplementary SS11 we show that the results are qualitatively equivalent with far more accurate fits.

The minimal model reproduces inverse size tuning and the spatial profile of inverse response as a function of size (Fig. 5b-c): centered cells show high response to small inverse stimuli, while for large inverse stimuli offset cells show a large response and centered cells respond less. Adding SOM-cell input to the model, using experimentally measured SOM responses, slightly *reduces* the degree of inverse surround suppression, the opposite of the effect seen in the classical case (Fig. 5d, and see Fig. 3n for comparison). This is because SOM cells respond more strongly to small than to large inverse stimuli (Fig. 4b and Supplementary SS4 A), whereas they respond more strongly to large than to small classical stimuli (Fig. 2).

**Figure 5:**
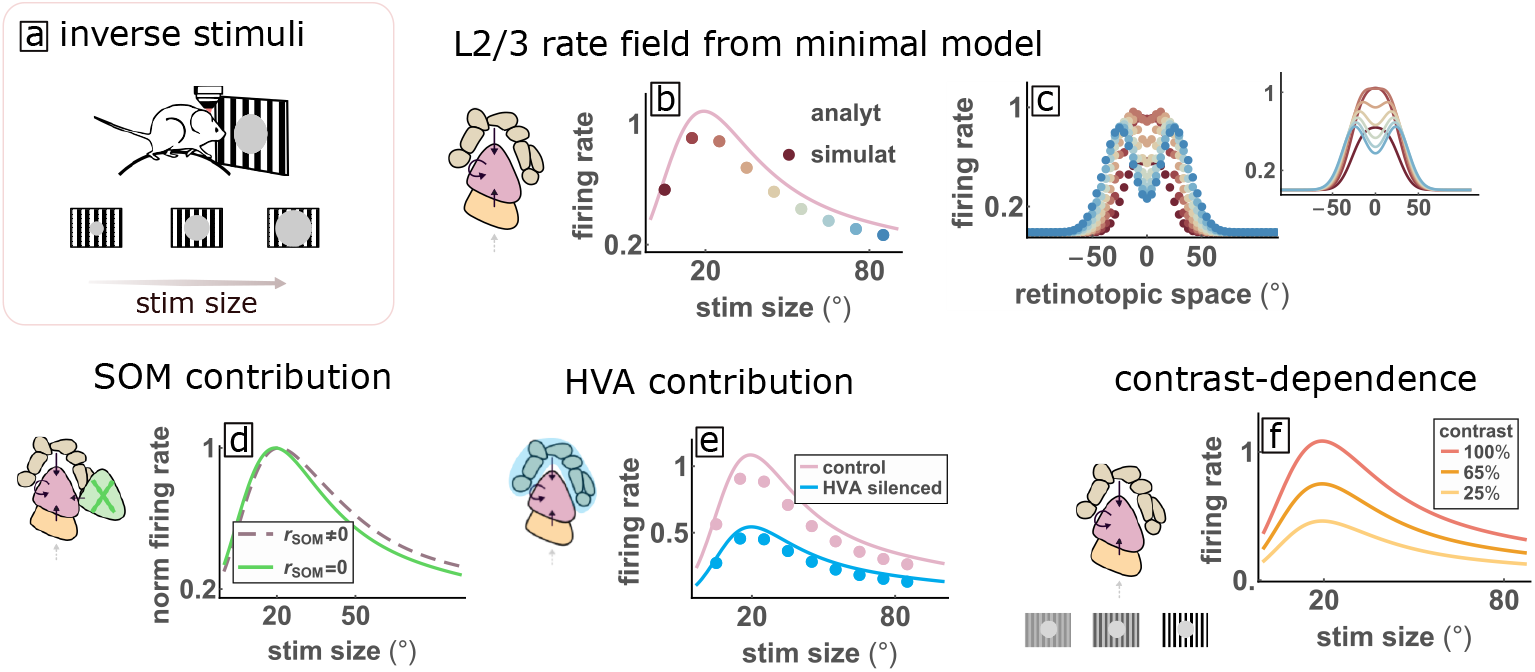
Inverse response and size tuning properties of L2/3 are shaped by its feedback input. **a** Inverse size-tuning stimuli, **b** sizetuning curve, and **c** rate fields for L2/3 of the model. In **c**, results for the simulations (dots) and the analytics (curves) are represented separately for better readability. The model recovers inverse size tuning as well as the change in convexity of the rate field (compare Fig. 4a,i). **d**. SOM neurons reduce inverse surround suppression (compare Fig. 3n for classical stimuli). **e**. Size tuning curve in the control case (pink) and for silencing of HVAs (rate field of HVAs reduced by a factor 0.65) (compare Fig. SS13 A for classical stimuli). **f**. Size tuning curves for varying contrast (both L4 and HVAs rate fields are reduced by a factor *c* = log_2_ 2*C*_0_, *C*_0_ values shown in legend; compare Fig. SF22 for classical stimuli).

Inverse response is substantially reduced by reduction of HVAs input by 35% (Fig. 5e). For classical stimuli, the same modification affects L2/3 response only marginally (Supplementary SS13 A). This is consistent with experimental results on optogenetic silencing of HVAs (see Fig.5c in (Keller et al., 2020b)). This differential effect of HVA suppression on inverse vs. classical response is consistent across varying levels of HVA suppression (Supplementary SS13 A). Finally, the model shows an increase of inverse responses with contrast (Fig. 5f, see also Supplementary SS13 B) similar to that found experimentally (Extended Data Fig. 3h of Keller et al. (2020b)).

#### 2.4.1 Inverse response requires wide enough feedback projections and feedback activity profiles scaling with stimulus size

We use our analytical framework to determine the conditions needed for inverse responses to arise. We first investigate the role of the width of the feedback projections *σ*_*L*23*−HV A*_. We vary its value around the experimental estimate (Fig. 6a,b, see also Supplementary SS1) while the inverse stimulus size is fixed at 15° (Fig. 6a). For this stimulus size, the L2/3 rate field is peaked at its center. For this to occur in the model, *σ*_*L*23*−HV A*_ must be sufficiently large, but not too large (Fig. 6c), as we now explain. HVAs activity peaks in a ring of cells with RFs corresponding approximately to the hole edge, although the ring size grows somewhat more slowly than the hole size (Fig. 4, SS12). We posit that each HVA cell on the ring projects a Gaussian-shaped bump of projections back to V1 L2/3, centered on its position on the ring. If these feedback projections are wide enough, the bumps will all overlap in the middle, giving the strongest input, and thus the strongest response, at the stimulus center (shown schematically in Fig. 6a and discussed more precisely in Supplementary SS14).

**Figure 6:**
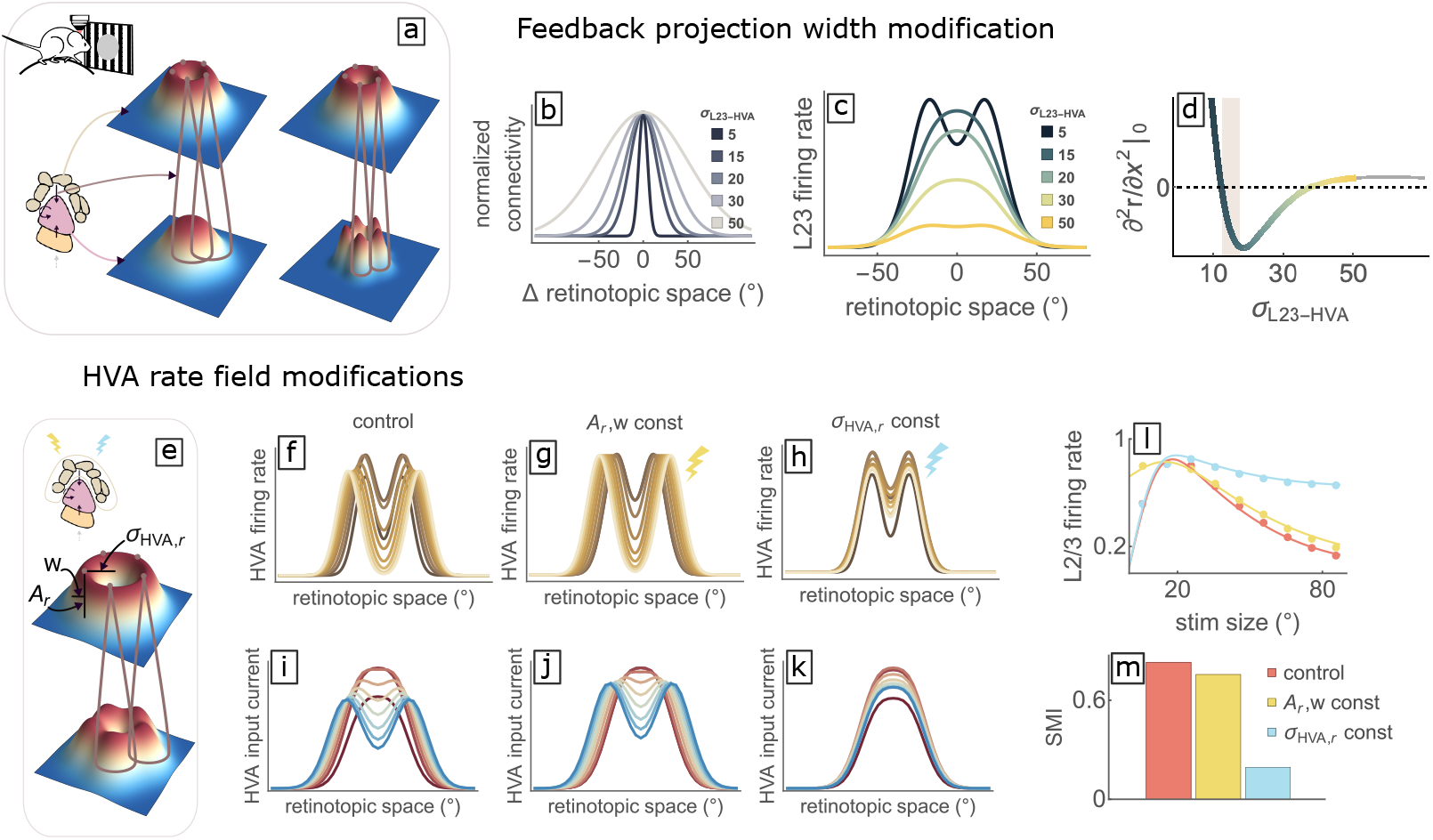
Inverse response requires wide enough feedback projections and feedback activity profiles that scale with stimulus size. **a**. Rate fields in HVAs for a small stimulus (top 3D-plots, ring profile). The feedback projections are represented by the empty cones (narrower projection width on the right). The bottom 3D-plots represent the L2/3 field of input current arising from the feedback projections from 5 example points along the ring of peak firing rate in HVAs; the L2/3 input current would be the sum of similar projections from all points in the HVAs rate field. Narrower feedback projections yield a trough in the center (right), while broader projections yield a peak (left). **b**. Counterfactual modifications of the width of projections from HVAs to L2/3, *σ*_*L*23*−HV A*_ (see STAR Methods), widths in legend. Amplitudes are normalized to 1. **c**. Rate field of L2/3 for the different values of *σ*_*L*23*−HV A*_ for a fixed stimulus width of 15°. Inverse response depends on the feedback projection’s width. **d**. Second derivative of L2/3 rate field evaluated at the origin, vs. *σ*_*L*23*−HV A*_. The color code indicates the value of *σ*_*L*23*−HV A*_ as in **c**, while the shaded bar corresponds to the experimental estimate in (Marques et al., 2018). **e**. Top: parametrization of HVAs rate field used for **f-m**: we fit the rate field as a convolution of a circle of radius *σ*_*HV A,r*_ with a Gaussian of standard deviation *w* and amplitude *A*_*r*_. Bottom: for fixed feedback projection width which would yield a center peak in **c**, a wide enough radius of HVAs activity, *σ*_*HV A,r*_, results in a trough in the center. **f-k**. HVAs rate field (**f-h**) and input current (**i-k**) for the control condition (**f**,**i**; parameters as in previous figures), and for two counterfactual modifications: either the amplitude *A*_*r*_ and the width *w* (**g**,**j**, dark yellow symbol) or the size of the ring *σ*_*HV A,r*_ (**h**,**k**, light blue symbol) are kept constant (fixed at their value for size *s* = 15^*°*^) with varying stimulus size. **l**. Size tuning curve and **m**. Surround Modulation Index for the control condition (red) and the modified conditions (colors as in **g**,**h**)

To quantify the range of values of *σ*_*L*23*−HV A*_ compatible with the inverse response, we compute the second derivative of the recurrent rate field at its center as a function of *σ*_*L*23*−HV A*_ for the 15° stimulus (Fig. 6c-d). A negative second derivative denotes a maximum in the center, as observed experimentally. For this to occur, *σ*_*L*23*−HV A*_ needs to fall within the range (12^*°*^, 35^*°*^) (Fig. 6d). The experimental estimate in (Marques et al., 2018) is indicated by the shaded area in Fig. 6d, which falls within this range. The same results hold in a model of L2/3 with 4 recurrent cell types (see Supplementary Fig. SF39).

We can similarly understand why inverse response is size tuned. As the stimulus size increases and the ring-shaped HVAs rate field becomes wider, the input to L2/3 cells centered on the stimulus center decreases (the overlap of the bumps in the middle decreases, see Fig. 6e). At some point, the input peak moves away from the center, and an ever-deepening trough forms at the stimulus center (Fig. 6i), representing the decrease in response with increasing stimulus size (Fig. 6f).

While this explanation attributes the decrease in response with increasing hole size to the increasing width of the ring of HVAs activity (which we also show analytically, Supplementary SS17), another contributing factor could be the decrease in amplitude and thickness of the ring. To test this, we fit the HVAs rate field with an alternative parameterization, a circle of radius *σ*_*HV A,r*_ (ring radius) convolved with a Gaussian of amplitude *A*_*r*_ and width *w* (ring thickness; see Fig. 6e). We then apply two counterfactual modifications to the HVAs rate field: i) eliminate the size-dependence of *A*_*r*_ and *w* while preserving the scaling of *σ*_*HV A,r*_ with stimulus size (dark yellow symbol, Fig. 6g), ii) eliminate the scaling of *σ*_*HV A,r*_ while preserving *A*_*r*_ and *w* (light blue symbol, Fig. 6h). If our explanation is right, (i) should only have a minor effect while (ii) should significantly reduce inverse surround suppression.

Indeed, in case (i), the field of HVA input to L2/3 develops a trough for large stimuli (Fig. 6j), and the inverse size tuning curve is very similar to that in the control case, with a minor decrease in SMI (Fig. 6l-m). The only notable change is that the response to the smallest size hole, 5°, becomes almost as large as the preferred size response. In contrast, in case (ii), the HVA input field does not develop a trough as the stimulus size increases (Fig. 6k) and, correspondingly, inverse surround suppression is drastically reduced (Fig. 6l-m). Note that, for a classical stimulus, surround suppression is only slightly decreased by an analogous modification (compare Fig. 3e and Fig. 6m, light blue bar).

This difference between the classical and inverse cases in the effect of a lack of widening of the input rate field suggests that, despite the similarity of their size tuning curves, their origin is substantially different. Our model shows that classical surround suppression in L2/3 is mostly inherited from the aligned cells of its inputs (and thus is not impacted by widening input) and enhanced by both PV inhibitory input and the SOM-VIP subcircuit. On the other hand, inverse size tuning is mediated by offset cells in HVAs. As the stimulus becomes larger, peak responses in HVAs move to cells at larger offsets from the center, providing less and less excitatory input to the center as the stimulus radius grows beyond the span of the feedback projections.

In Supplementary Fig. SF39 we show that the explanations derived using the minimal model also apply to a more biologically realistic model with 4 cell-types (see in particular SS14).

The arguments presented here provide a purely geometrical explanation of inverse response and inverse size tuning, illustrating the importance of considering a spatial framework with realistic anatomical and physiological length scales to understand contextual effects.

### 2.5 Surround facilitation can be explained by the inverse response to the orthogonal surround

When a stimulus is presented together with an orthogonal surround (“cross” stimulus), the response of L2/3 – but not L4 – Pyr cells centered on the stimulus is facilitated (Keller et al., 2020a; Shushruth et al., 2012a; Sillito et al., 1995). We propose that this effect may be, at least in part, the counterpart of the inverse response. More specifically, the response of aligned Pyr cells may increase due to the orthogonal surround evoking the same inputs that drive the inverse response. To test this hypothesis, we extend the minimal model to include orientation preference (see Fig. 7a and Supplementary SS18) and connectivity that decreases in strength with increasing difference in preferred orientation according to experimental estimates (Ko et al., 2011, 2013; Rossi et al., 2020) (see Supplementary SS1).

**Figure 7:**
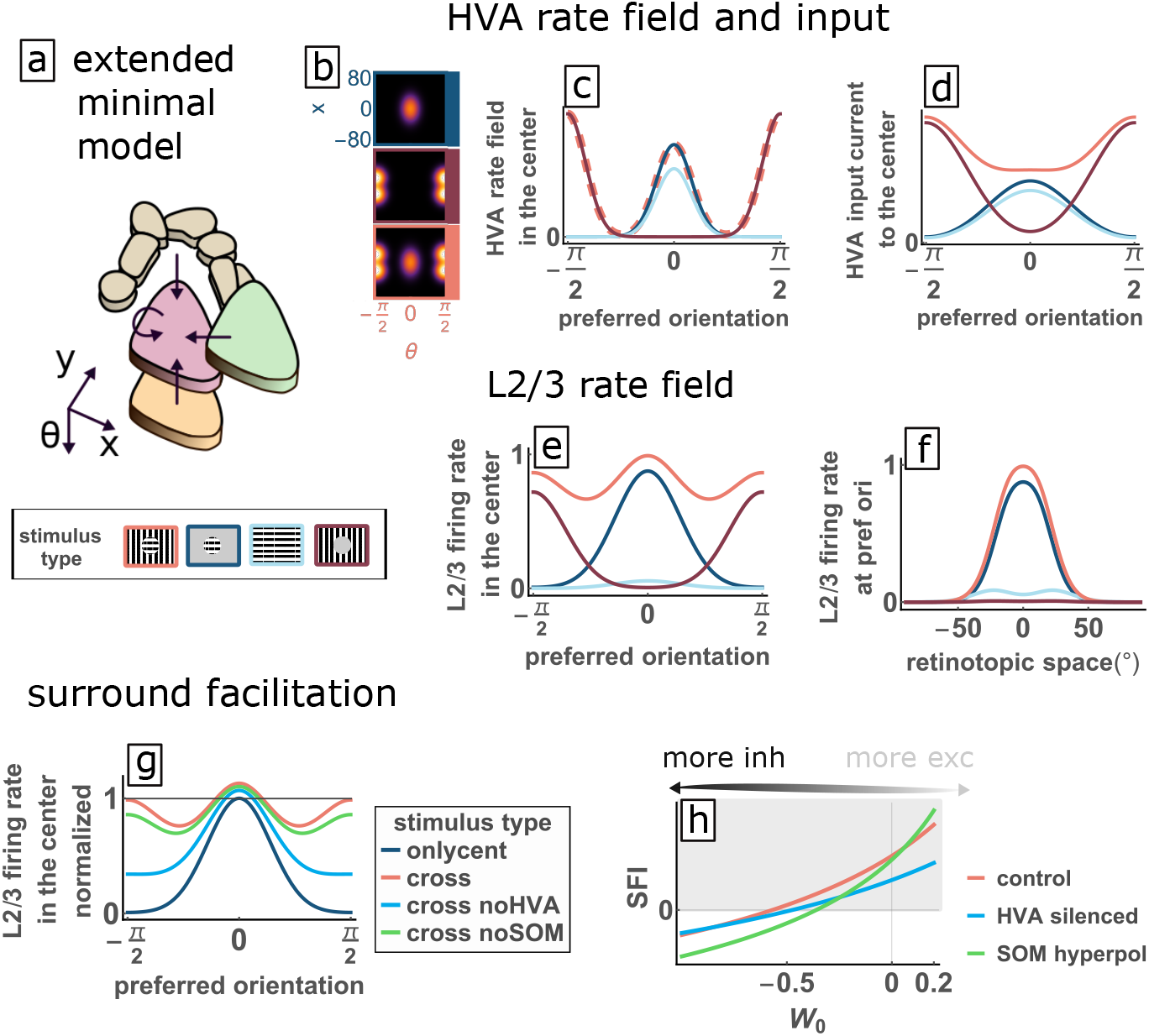
Surround facilitation is generated by inverse response to the orthogonal surround. **a**. Sketch of the extended minimal model, accounting for orientation preference as an additional dimension for the rate fields. Bottom: Legend of the stimuli analysed: cross (center at orientation 0, surround at *π/*2; light red), classical/only center (dark blue), iso-oriented surround (light blue), inverse (dark red). **b**. Density plot of HVAs rate fields in one spatial dimension (vertical axis) and orientation preference (horizontal axis) for classical, inverse and cross stimuli (color code as in the legend). The rate field of the inputs for the cross stimulus (bottom) is the sum of a classical stimulus at orientation *θ* = 0 (top) plus an inverse stimulus at *θ* = *π/*2 (middle). **c**. HVAs rate fields of cells whose CRF is centred on the stimulus center and with all possible orientation preferences (horizontal axis). Note, light red line for cross stimulus is shown dashed and thick, so overlapping lines (dark blue, central region; dark red, larger preferred orientations) can be seen; light blue and dark blue (not visible) overlap for larger preferred orientations. **d**. Input current from HVAs to the centered cells in L2/3 as a function of their preferred orientation. Note that the input from L4 is much stronger in the center than in the surround (not shown). **e-f**. The response of centered L2/3 cells preferring orientation *θ* = 0 is larger for the cross stimulus than for the classical stimulus alone (surround facilitation). **e**. Rate field of L2/3 neurons centered on the stimulus center, as a function of their preferred orientation. **f**. Rate field of L2/3 cells that prefer orientation *θ* = 0, as a function of their retinotopic location. **g**. We study the cross condition (control) and two manipulations of it (no HVA, no SOM). For each condition, rate fields of L2/3 are normalized to the amplitude of the response to a center-only stimulus. Surround facilitation (responses larger than 1) decreases when HVAs are silenced or SOM cells are hyperpolarized. **h**. Dependence of surround facilitation index (SFI) on the amplitude of the recurrent connections *W*_0_. SFI decreases both when HVAs or SOM cells are silenced. Larger recurrent excitation (less negative or more positive *W*_0_) contributes to larger SFI.

We assume that the rate fields of the input populations for the cross stimulus can be approximated as the sum of the rate fields for a direct stimulus of one orientation in the center plus an inverse stimulus of the orthogonal orientation (Fig. 7b,c). In general the response fields are not additive. Nevertheless, for cross stimuli the classical and inverse stimuli are largely encoded by complementary sets of cells, i.e. cells with orthogonal orientation preferences, hence we make the assumption of additivity.

We consider a classical stimulus with orientation 0, an inverse stimulus with orientation *π/*2, and a cross stimulus that combines the two. The extended minimal model shows that L2/3 cells are surround facilitated (Fig. 7e-f). This effect can be understood as follows: HVAs cells at the stimulus center with preferred orientation 0 have zero response to the inverse stimulus (dark red line in Fig. 7c). However, L2/3 cells with the same position and preferred orientation receive small but positive HVA input (Fig. 7d), which comes from more offset HVA cells via their broad feedback projections to L2/3. Thus the inverse stimulus at orientation *θ* = *π/*2 provides a small HVA input to the L2/3 centered cells that prefer orientation *θ* = 0. This input generates cross-oriented facilitation (light red curves in Fig. 7e-f).

Thus, our model suggests that HVAs support L2/3 cross-orientation surround facilitation (Fig. 7g, light red curve). If HVAs are blocked (cyan curve), responses to both center-only and cross stimuli decrease; cross is still facilitated relative to center-only, but less so than with HVAs intact (the cyan curve in Fig. 7g lays below the light red curve). The same is true also when SOM input is silenced (green curve), implying that both HVAs and SOM cells support cross-orientation surround facilitation.

To better quantify surround facilitation, we define the Surround Facilitation Index: *SFI* = (*r*_x_ *− r*_c_)*/*(*r*_x_+*r*_c_), where *r*_x_ (*r*_c_) is the firing rate of the L2/3 centered cells in the cross (classical) stimulus condition. We analyse the dependence of SFI on the relative weight of excitation and inhibition in the joint Pyr+PV population, controlled by the amplitude of the recurrent connections *W*_0_ (Fig. 7h). Larger *W*_0_ (i.e. more recurrent excitation) yields larger SFI. For excitation dominated systems (*W*_0_ *>* 0), SOM input suppresses surround facilitation (green curve above the light red one). On the other hand, for inhibition-dominated systems (*W*_0_ *<* 0), as in Fig.7g, SOM input increases SFI (green curve below the light red). More generally, in Supplementary SS19), we show that SOM input increases (decreases) Pyr+PV SFI if the ratio of SOM input in the inverse vs. center conditions is less (greater) than the ratio of total input in the two conditions.

To summarize, assuming that inputs evoked by center-only and orthogonal surround-only or “inverse” stimuli add linearly, both HVAs and lateral SOM inputs contribute to cross-orientation surround facilitation, while lateral PV input tends to reduce it. Note that the suppression of SOM was the primary mechanism of enhancement of response to the cross-oriented stimulus proposed in (Keller et al., 2020a) (in that case due to activation of VIP by the cross stimulus, which inhibited SOM), here we are proposing that both HVAs and the suppression of SOM contribute to the facilitation.

## 3 Discussion

We propose a unifying model accounting for three types of contextual modulation and connecting two previously unrelated phenomena: inverse response and cross-orientation surround facilitation. To our knowledge no previous model has addressed the emergence of inverse responses. Our model suggests that classical surround suppression in L2/3 is largely inherited from surround suppression of its feedforward and feedback inputs (consistent with feedforward and feedback contributions noted in Angelucci et al., 2017; Vangeneugden et al., 2019). Size-tuned inverse response arises from the geometry of feedback projections from HVAs to V1 L2/3, along with a ring-like HVAs spatial pattern of activity to an inverse stimulus that widens with increasing stimulus size. Cross-orientation facilitation can be understood as a combination of classical center response and inverse response.

All previous models of surround suppression have assumed it is mediated by lateral inhibition progressively recruited as the stimulus size increases (Angelucci and Bressloff, 2006; Angelucci et al., 2017; Dipoppa et al., 2018; Li and Young, 2021; Obeid and Miller, 2021a; Rubin et al., 2015). However, we find that, in mouse L2/3 of V1, this mechanism only marginally contributes to surround suppression, because the rate field size increases much more slowly than the stimulus size. Furthermore, the rate field size shows little or no growth with stimulus size in L4, so this mechanism also should not contribute to L4 surround suppression. This holds true independently of the preprocessing procedure used to define the rate fields (Supplementary SS2 B), and is consistent with the recordings in Dipoppa et al. (2018) (their Supp. Fig. 11D) and in Michaiel et al. (2019) (their Fig.2A), although those authors did not note the effect. Pinpointing this counterintuitive feature of neural responses in mouse V1 is one of our major findings. We should, however, caution that all of the recordings in this and those two papers were done with Ca^++^ imaging, and it is conceivable that there is widening of the spiking signal but with intensity below threshold for Ca^++^ recording.

Our finding suggests the interesting psychophysical correlate that the perceived area of stimuli should grow more slowly than the actual area. This is in agreement with the results of Yousif and Keil (2019, 2021), who had participants compare total area of multiple circular patches of color, distributed in random locations within a square region, and with different diameter distributions. For samples with equal total area, participants consistently perceived less cumulative area in samples with fewer larger patches than in samples with more smaller patches. This correlate could be further tested with another experiment: alternately present a small circular stimulus centered at position **x**, and a larger circular stimulus whose edge intersects **x**. We predict that the larger stimulus should be perceived as not extending to the center of the smaller one.

To gain robust mechanistic insight, we include in the minimal model as few elements as possible, to easily disentangle the effects of each. Moreover we keep the parameter space as low-dimensional as possible (as little as 3-dimensional). Instead of quantitatively fine tuning the model to a particular data set, we aim to describe results at a qualitative level, at which they are more robust and consistent across different data sets. While the minimal model neglects the full recurrent microcircuitry of L2/3, it yields mechanistic insights into cortical computation, which we show also apply to a fully recurrent 4-cell-type circuit (Supplementary SS17).

We find that classical surround suppression in L2/3 is enhanced by increasing SOM lateral input (Adesnik et al., 2012a; Nienborg et al., 2013b), reduced by an increase in the firing of PV cells (Nienborg et al., 2013b), and partially supported by the supralinear neuronal input/output function (Hansel and van Vreeswijk, 2002; Miller and Troyer, 2002; Rubin et al., 2015). In addition, we find that the contrast dependence of classical surround suppression is also largely inherited from L4 inputs(Mossing et al., 2021), and show that the spatial response profile of Pyr and PV cells can change from a peak in the center to a trough in the center for increasingly large classical stimuli, as observed in Dipoppa et al. (2018), if SOM input is sufficiently strong.

Keller et al. (Keller et al., 2020b) highlight the importance of input from retinotopically offset neurons in HVAs for the inverse response. Here, we adopt a continuous description of retinotopic space, as in (Dipoppa et al., 2018; Li and Young, 2021), to preserve the information about length scales of the spatial patterns of activity. In addition, we constrain the range of the projections between different populations based on anatomical results (Billeh et al., 2020; Campagnola et al., 2021; Karnani et al., 2016; Li and Young, 2021; Pfeffer et al., 2013; Rossi et al., 2020). This allows us to characterize the inverse response in terms of the geometry of the feedback input.

Our schematic description of space opens the way to an analytic treatment of the problem, following and generalizing (Persi et al., 2011). This allows us to determine robust determinants of qualitative response properties, such as: the input properties that are necessary to give rise to surround suppression and that determine the preferred size of the response; possible causes for the decrease of the preferred size when the stimulus contrast is increased; the conditions needed for inverse response rate fields to peak at the center; and the conditions required for inverse surround suppression.

We treat HVAs as a static input, even though they are part of a coupled system, receiving input from V1 and projecting back to V1. We could not model HVAs dynamically because, in the data set considered, L2/3 responses to small-size classical and inverse stimuli are very similar, while LM responds very differently to the two stimuli. This implies that LM must receive a significant part of its input from other sources, which we are not modelling (Blot et al., 2021; Harris et al., 2019; Tohmi et al., 2014).

We modeled HVAs input based on recordings from Area LM (Keller et al., 2020b). However, in these recordings, LM responses are much weaker in the inverse condition than in the classical condition. Since inverse responses have similar strength to classical responses, we assumed that input from other HVAs must equalize overall HVA responses in the two conditions, and scaled observed LM inverse responses accordingly. This assumption needs to be tested experimentally.

Despite the very limited number of parameters, our minimal approach is highly informative. In addition to the three overall phenomena of classical and inverse size tuning and surround facilitation, it reproduces stabilization of the circuit by PV cells, the modulations of the classical and inverse response when silencing HVAs, and the effects of changing stimulus contrast and hyperpolarizing SOM cells. At the same time, it generates insights that translate into several experimentally testable predictions, as summarized in Table 1 and as we now discuss.

### 3.1 Testable predictions

SOM neurons reduce the degree of inverse surround suppression (Fig. 5d). Thus, we predict that optogenetically suppressing SOM cells should boost the L2/3 response to smaller inverse stimuli more than to larger inverse stimuli.

We reason that the spatial activity patterns evoked by a center stimulus of one orientation and an inverse stimulus of the orthogonal orientation should add up roughly linearly, to give the spatial pattern evoked by a cross-oriented center/surround stimulus. This can be experimentally tested by recording the spatial pattern of responses to center-only and orthogonally-oriented inverse stimuli to see if their sum indeed predicts the spatial response to cross stimuli. The model also predicts that under presentation of the cross stimulus, L2/3 Pyr and PV cells centered on the stimulus and with preferred orientation orthogonal to the center stimulus should show significant response (Fig. 7c).

More generally, our model raises many interesting questions in understanding how the orientation tuning of classical and inverse responses are related to one another and to cross responses. In our model, we assume that all neurons have the same OSI and that the orientation preference for classical and inverse stimuli is the same. On the contrary, Extended Data Figure 3 in Keller et al. (2020b) shows that for neurons with high OSI for both classical and inverse stimuli, a significant fraction of neurons have a preferred orientation for classical stimuli that is orthogonal to their preferred orientation for inverse stimuli. Our model predicts that these cells should have the strongest cross-orientation surround facilitation, since they would have the strongest response to the cross surround. They might also have the strongest surround suppression, since they would have the weakest top-down input evoked by the iso surround. More generally, it will be interesting to see if responses to orthogonal center and inverse stimuli add linearly, regardless of a neuron’s relative orientation preference for center vs. inverse stimuli, and how these relative preferences impact surround suppression and cross-orientation facilitation.

Our model predicts that optogenetically silencing HVAs during cross stimulus presentation should reduce surround facilitation. Likewise, the model predicts that hyperpolarizing or optogenetically silencing SOM cells during cross stimulus presentation should also reduce surround facilitation. These points are illustrated in Fig. 7g,h.

The fact that the SFI increases as a function of the relative weight of excitation and inhibition (Fig. 7h) predicts that, when optogenetically silencing a fraction of PV neurons, surround facilitation should increase. Fig. 7h also shows that SFI drops when HVAs or SOM inputs are reduced.

Context shapes the readout not only at a neural level, but also at a perceptual level. Animals are active interpreters of the visual scene, as opposed to passive decoders (Agrillo et al., 2015; Agrochao et al., 2020; Bååth et al., 2014; Bach and Poloschek, 2006; Carbon, 2014; Eagleman, 2001; Endler et al., 2010; Gori et al., 2014; Keemink et al., 2018; Luo et al., 2019; Mély et al., 2018; Pak et al., 2019, 2020; Santac`a et al., 2019). The study of the mechanisms underlying contextual modulation of responses, including information that higher processing stages project back onto the canvas of visual cortex, opens a window to understand how our brains create our perceptual experiences.

We believe that the simplicity, analytical tractability, and spatial nature of the model presented here, together with the fact that it can be easily extended to account for other feature spaces, will enable further mechanistic insight into visual cortical circuitry and responses in future studies.

## Acknowledgments

We acknowledge funding sources NIH U01NS108683, NIH R01EY029999, NIH U19NS107613, NSF 1707398, Gatsby Charitable Foundation GAT3708.

## Author Contributions

Conceptualization, K.D.M., S.D.S and M.D.; Methodology, S.D.S.; Software, S.D.S.; Formal Analysis, S.D.S; Experimental Data, A.K., M.R.; Writing-Original Draft, S.D.S.; Writing-Review and Editing, K.D.M., S.D.S. and M.D.; Visualization, S.D.S.; Supervision, K.D.M.; Funding Acquisition,K.D.M., M.S.

## STAR Methods

### RESOURCE AVAILABILITY

#### Lead contact

Further information and requests for resources and code should be directed to and will be fulfilled by the lead contact, Serena Di Santo

#### Materials availability

This study did not generate new unique reagents.

#### Data and code availability

All original code will been deposited and is publicly available as of the date of publication. DOIs are listed in the key resources table.

## METHOD DETAILS

### The minimal model

We consider a system with one recurrent population defined on a two-dimensional retinotopic space of coordinates *{***x** *}*, whose rate field is *r*(**x**) and which receives a visually-driven input described by a 2D isotropic Gaussian function *I*(**x**) = *I*_0_ *G*(**x**, *v*) (where 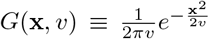). Let the recurrent connections be *W* (**x** *−* **y**) = *W*_0_*G*(**x** *−* **y**, *v*_*r*_). We consider the SSN dynamical equations for the system, 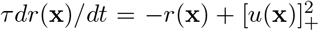 where *u*(**x**), the total input current, i.e. the sum of the visually driven and recurrent input. We will hereafter focus on the steady state of the system, where 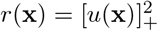, or:

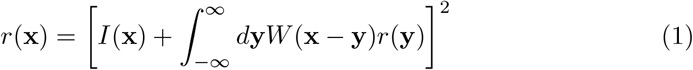

In Eq.1 we omit the rectification sign, based on the assumption that *u*(**x**) is always greater than or equal to 0. We will show that the results we then derive obey this assumption.

Analogously we can write the SSN equations for the total input current at steady state as:

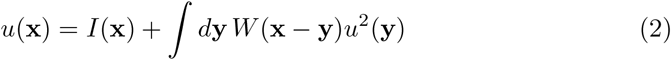

To find *u*(**x**) we make the ansatz that *u*(**x**) has the following form:

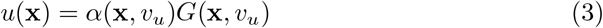

where *α*(**x**, *v*_*u*_) is required to vary slowly as a function of **x**. We will henceforth write it simply as *α*(**x**), leaving the dependence on *v*_*u*_ implicit. Note that *v*_*u*_ is a free parameter that is constrained only by the fact that when we solve our equation the resulting *α*(**x**) must be slowly varying. Then the recurrent term is:

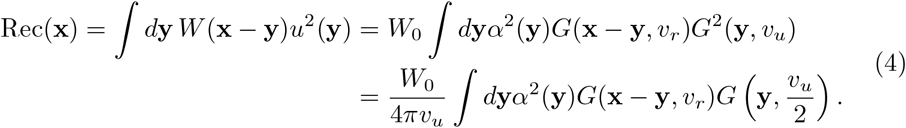

The largest contribution to the integral is given by the maximum of its argument (Laplace approximation), but since *α*^2^(**y**) varies slowly (*α*^*’*^(**y**) *« α*(**y**)), the maximum of the full argument can be approximated with the maximum of 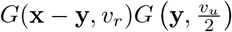, which occurs at 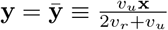

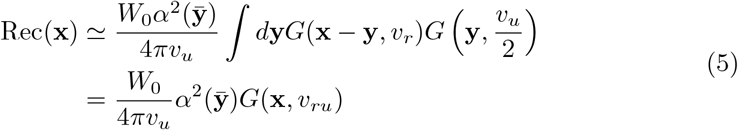

where we defined 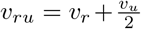. Since *α*(**y**) varies slowly with its argument, we take the further approximation that 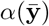 is well approximated by *α*(**x**). From Eq. 2 we get:

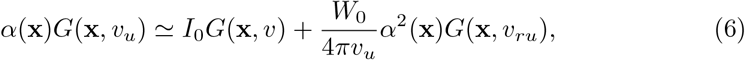

We now solve the equation above for *α*(**x**) and get:

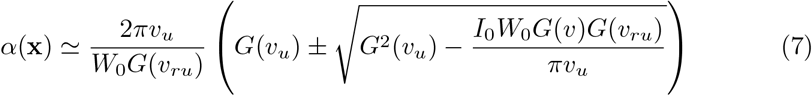

where for better readability we are omitting the spatial dependence of the function *G*(*·*). In Supplementary SS7 A we show that *α*(**x**) *« α*^*’*^ (**x**).

We now choose *v*_*u*_ to satisfy:

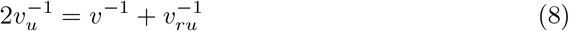

This simplifies the expression under the square root, making it independent on **x**:

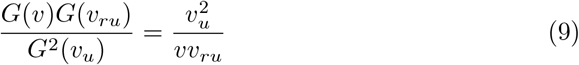

and thus we can write:

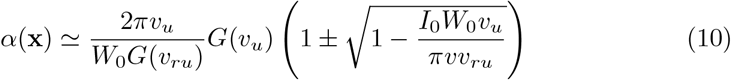

Finally, using Eq. 9 again:

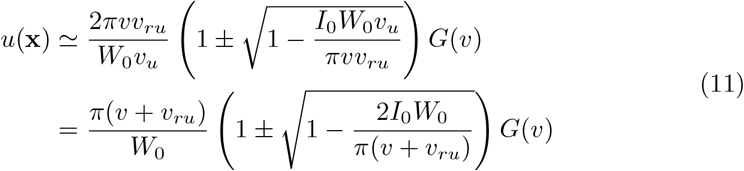

Note that only one of the two solutions is actually acceptable, depending on whether the recurrent population described is excitatory or inhibitory. The only acceptable solution is the one with *−* for negative *W*_0_ (because the one with + gives *u*(**x**) *<* 0). For positive *W*_0_, instead, the solution requires that 2*I*_0_*W*_0_ *< π*(*v* + *v*_*ru*_). Violation of this condition means that recurrent excitatory input is very large (*W*_0_ *> π*(*v* + *v*_*ru*_)*/*2*I*_0_), and in this case we find numerically that the system diverges (see Supplementary SS7 B).

Analogously, if the external input can be described by a sum of Gaussians:

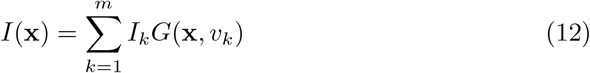

making the same ansatz as above, we obtain

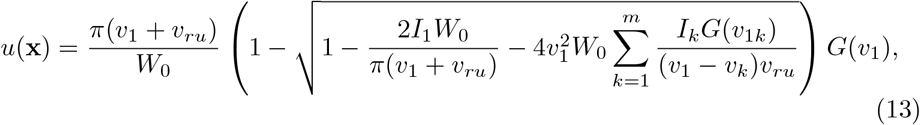

where we defined

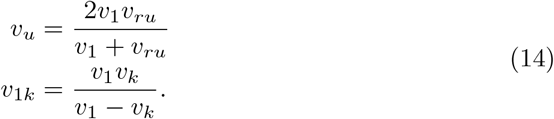

Note that, despite the multiple approximations, the analytic solution provides a really good description of the rate field (see *e*.*g*. Fig. 3c,e and Fig. 5a,c).

## QUANTIFICATION AND STATISTICAL ANALYSIS

### Data preprocessing

The distance between the center of the stimulus and the center of the CRF of cells is estimated in (Keller et al., 2020b) through CRF mapping experiments, where a stimulus is presented at many positions spanning the area of the visual field accessible for recording and the response of thousands of neurons are recorded through 2-photon calcium imaging (Keller et al., 2020b). Since the distance between contiguous presentations of the stimulus is 5^*°*^, we take this value to be the error on the measure of the CRF location. For each cell we then take a Gaussian in its spatial location with *σ*_*err*_ = 5^*°*^ and with amplitude equal to the recorded (trial averaged) response of that cell, i.e. 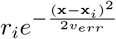 with 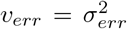. We then sum the Gaussian functions for all recorded cells and divide by the sum of unitary Gaussians, to account for the non-uniform distribution of data-points. The rate field of population *A* is thus given by:

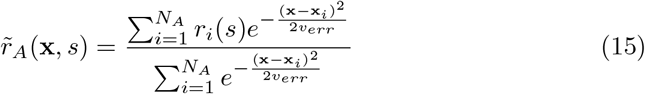

where *N*_*A*_ is the number of recorded cells of type *A* and *s* indicates the stimulus size. In Supplementary SS2 B we show an example of the smoothening that this procedure (first presented in (Dipoppa et al., 2018)) generates. Rate fields are plotted in 1 spatial dimension, but they are in fact 2-dimensional and isotropic in retinotopic space. The retinotopic space is considered continuous, representing the limit of very large number of neurons.

### Functional form of the rate fields

In order to be able to understand the system analytically we take the approximation:

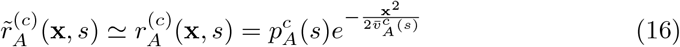

where *A* = *E, P, S, V, L, M* indicate respectively populations of Pyramidal, PV, SOM, VIP, Layer 4, LM cells and the superscript ^(*c*)^ stands for classical. The prefactor 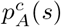 represents the size tuning curve, whereas the dependence on retinotopic space is purely Gaussian. Operatively, we fit the experimental rate fields for each stimulus condition with a Gaussian function. We report the goodness of fits in Supplementary SS4.

For inverse rate fields the response profiles are more complex, therefore we schematize them through a difference of Gaussian functions:

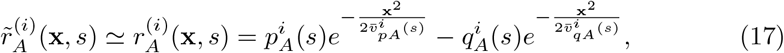

with 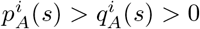.

Note that in the inverse stimulus condition we define scaling as the growth of the width of the outer Gaussian of the fit when stimulus size is increased. We do not distinguish between outer and inner scaling because they appear to be very similar. The difference between outer and inner scaling is mostly very small (less that 1 degree) and:

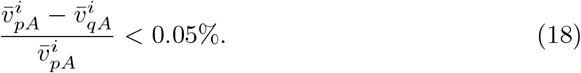

Moreover its distribution is very narrow.

In order to have a full functional description of the rate fields as a function of the classical stimulus size we need to define the size tuning curve 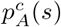 for the populations that we are going to use as inputs for the minimal model, i.e. *A* = *S, L, M*. This will allow to give an analytical account of surround suppression of the inputs by analyzing the interval where 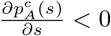 and preferred size (or CRF size) as 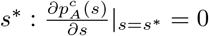 (see Supplementary SS8). We use the parametrization:

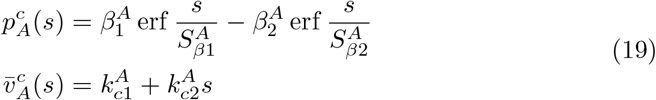

where the first equation encodes the size tuning curve and 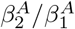 modulates the amount of surround suppression, 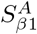 modulates the steepness of the increase of size tuning for small sizes and 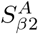 modulates the steepness of the decrease of size tuning for large sizes. Moreover 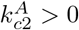 indicates the growth of the rate field width with stimulus size.

The size tuning curves described by the functional form in 19, together with the data and the fitted samples of the rate fields are shown in Supplementary SS4 A.

For the inverse stimuli we need to specify the functions 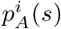 and 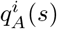. We devise such functional forms and show a comparison with the experimental rate fields.

For SOM population we use the parametrization:

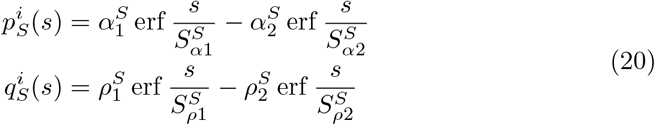

The evidence presented in (Keller et al., 2020b) suggest that HVAs highly influence the inverse size-dependent response profiles, and on the contrary, the input from L4 is very weak in the inverse stimulus condition, due to classical surround suppression, thus while we focus on reproducing accurately the spatial nuances of LM, we allow less accuracy for L4, in order to keep the model simple and the number of fitted parameters as small as possible (for a more nuanced parametrization of the inputs see Supplementary SS11). Thus for L4 cells we use:

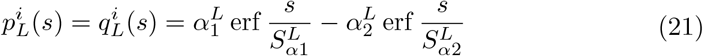

that constrains the center to have zero response, consistently with the very low response of aligned cells in L4 shown in (Keller et al., 2020b). Finally for LM we use:

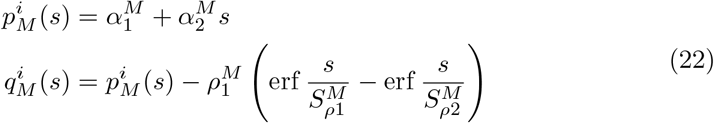

Note that the size tuning curve of the aligned cells can then be described as 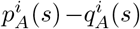. The size tuning curve of the offset cells is here defined as: 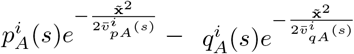 with 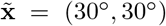. Aligned and offset size tuning curves pre and post parametrization are shown in Supplementary SS4 A. All the parameters *α, β, S, k, ρ* are obtained by fitting the recorded rate fields.

Finally the growth of the inner and outer scaling is given by:

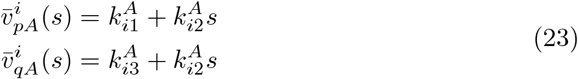

The recorded rate field of LM shows a much lower amplitude in the inverse stimulus condition versus the classical one (as evaluated for example computing the integral over space of the rate field). If the input from HVAs consisted solely of LM contribution, we would not be able to recover an inverse response in L2/3 with firing rates as high as or even higher than the classical case, reported in (Keller et al., 2020b). But we know that V1 receives input from all HVAs. Moreover Extended Data Fig. 9 in (Keller et al., 2020b) shows that optogenetic silencing of an individual HVA generates a quantitatively different reduction of classical versus inverse responses for each HVA considered. Thus, to quantitatively recover the relative amplitude of the rate field in L2/3, we take the minimal assumption that HVAs considered all together have the same rate field profiles of LM, but with an amplitude that is inferred according to the amplitude of the inverse response of L2/3 cells. In formulas:

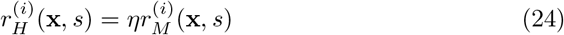

and from here on we will consider *A* = *{E, P, S, V, L, H}*, where *H* is a population representing the conjunction of all HVAs (see Supplementary SS5).

### Input currents

For the classical stimulus condition, the input to retinotopic position **x** of the recurrent layer is given by a convolution with the inter-layer connectivity profile 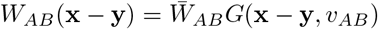, where 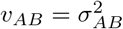:

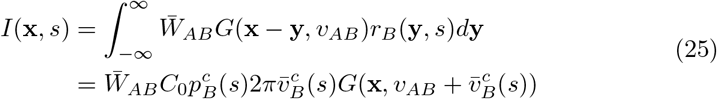

Thus we can read:

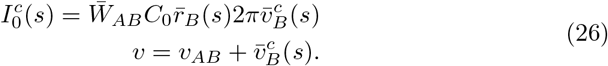

Finally, putting together the functional forms that we derived in the paragraph above, with Eq. 26, we can readout the full dependence of the total input current field of the recurrent population on the stimulus size.

For the inverse stimulus condition, the input to the recurrent layer can be computed through a convolution with the inter-layer connectivity

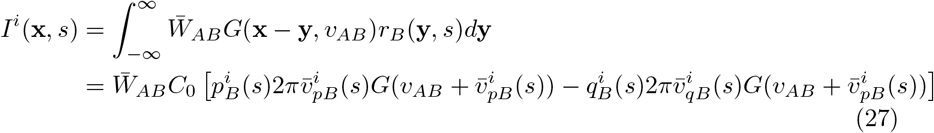

thus more compactly

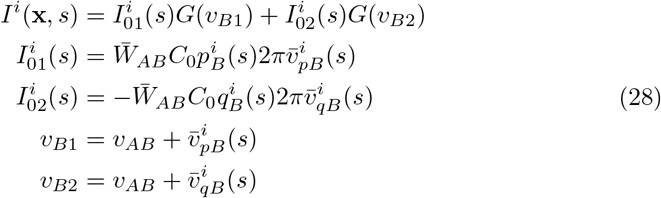

