## supplemental text and figures for "The combination of feedforward and feedback processing accounts for contextual effects in visual cortex"

(Dated: January 23, 2024)

#### CONTENTS

SS1. Anatomical length scales 3

SS2. Experimental rate fields 4
- A. Data preprocessing 4
- B. Rate fields calculated with two different preprocessing procedures 4
- C. Weak increase in activated V1 area with increasing stimulus size 4

SS3. Are aligned and offset LM neurons intrinsically different? 8

SS4. Gaussian approximation for the retinotopic spatial profile of the rate fields 10
- A. Parametrization of classical and inverse size tuning curves 12

SS5. From LM to HVAs 14

SS6. Rate fields for the joint population of Pyr and PV 15
- A. Comparison with a different dataset 16

SS7. The minimal model with inputs from L4 and HVAs 17
- A. Validation of the minimal model 17
- B. The joint population of Pyr+PV is stable if inhibition dominates 17
- C. Dependence of the firing rate of L2/3 on the FF and FB connections strengths 18

SS8. Analytical insights on classical size tuning curves with non-normalized input currents and comparison to previous models of SS 19
- A. Dissecting Classical surround suppression 19
- B. Classical surround suppression with non-surround suppressed input rate fields 21
- C. Preferred size depends on the preferred size of the inputs 21
- D. Preferred size decreases with contrast 21

SS9. Noisy neural dynamics enhance the surround modulation of the recurrent layer 23

SS10. Modulations induced by stimulus contrast 23
- A. Contrast dependent classical size tuning curves 23
- B. Recovering preferred size dependence on contrast 25

SS11. More nuanced parametrization of the inputs in the inverse stimulus condition 26

---

*

SS12. HVAs cells close to the edge of the inverse stimulus respond maximally 27

SS13. The minimal model for inverse response and inverse size tuning 28
A. Optogenetic silencing of HVAs 28
B. Contrast dependent inverse size tuning curves 28

SS14. The origin of inverse response and inverse size tuning 29
A. Analytical results for the minimal model in the inverse stimulus condition 29

SS15. A computational model trained to fit cell-type-specific recordings for classical stimuli predicts inverse responses 31
A. Non-Negative-Least-Squares approach to fit the full model 33
B. Inputs to SOM cells in the full model 35
C. Model selection for the full model 35

SS16. Robustness of the full model against errors on the measure of the projection widths 37

SS17. The full model exploits the same mechanisms as the minimal model 38
A. PV cells stabilize the system 38
B. Modulations induced by changing stimulus contrast or optogenetic manipulations in the full model 38

SS18. Extending the minimal model to include orientation tuning 41

SS19. Lateral input from SOM cells increases surround facilitation and the connection with SOM inverse response 42

References 44

#### SS1. ANATOMICAL LENGTH SCALES

We define a two-dimensional Gaussian function $G_{AB}(\mathbf{x}_1, \mathbf{x}_2, v_{AB}) = \frac{1}{2\pi v_{AB}} \exp[-\frac{(\mathbf{x}_1-\mathbf{x}_2)^2}{2v_{AB}}]$ to describe the connection strength to a cell of type $A$ at location $\mathbf{x}_1 = (x_1, y_1)$ from a cell of type $B$ at location $\mathbf{x}_2 = (x_2, y_2)$ Dipoppa et al. (2018); Rossi et al. (2020); Marques et al. (2018); Persi et al. (2011). We then estimate the length scales of the connectivity $v_{AB} = \sigma^2_{AB}$ based on recent experimental work. We assume isotropy in the azimuth and elevation directions ($x$ and $y$).

In a recent paper, Rossi et al. Rossi et al. (2020) could trace the excitatory (L4 and L2/3) and inhibitory (L2/3) presynaptic inputs to a L2/3 pyramidal neuron as a function of the horizontal distance. The average resulting curve shows an excellent agreement with a Gaussian function and the measured widths are $\sigma_{EE} \simeq 8^\circ$ and $\sigma_{EI} \simeq 5^\circ$, where $I$ indicates a generic inhibitory cell. Here and in what follows the magnification factor is taken to be $0.05^\circ/\mu m$ Rossi et al. (2020).

Distance dependent connections from Pyr to SOM cells in a slice of mouse L2/3 visual cortex were investigated in Adesnik et al. (2012). Adesnik et al. recorded the spiking activity of a L2/3 SOM in response to blue light spots of increasing diameters to activate progressively wider areas of L2/3. By cutting the slice to transect L2/3 horizontal axons, they show that SOM cells firing is significantly reduced when inputs from distances greater than $8^\circ$ were cut off. Although this does not exclude the possibility that in the control condition (uncut), nearby ($d < 8^\circ$) Pyr cells are more activated by offset Pyr cells, thus providing more excitation to SOM, the effect is still compelling and we draw from their measure a conservative estimate of $\sigma_{ES} \simeq 8^\circ$.

Marques et al Marques et al. (2018) measure retinotopic specificity in inputs from the lateromedial (LM) visual area in mouse V1 and quantify the retinotopic span of feedback projections from LM to V1. They report that LM inputs target, on average, retinotopically matched locations in V1, but many of them relay distal visual information. They estimate that about half of the visual coverage relayed by LM varicosities was more than $24^\circ$ away from their retinotopic position in V1. Assuming Gaussian connectivity, this yields a $\sigma_{EM} \simeq 15^\circ$. We assume that this value holds for HVAs in general, i.e. $\sigma_{EH} = \sigma_{EM}$.

In another recent paper, Billeh et al Billeh et al. (2020) also report the distance-dependent connection probability profiles for different classes of connections. The cortico-cortical connection probabilities for different cell-class pairs were estimated based on a survey of the existing literature, considering valid sources of information (in order): mouse visual cortex, mouse non-visual cortex, rat visual cortex, rat auditory cortex and rat somatosensory cortex. The measures were assumed to be integrated values of (distance-dependent) connections up to a certain average distance ($75\mu m$) between pre- and post-synaptic cells somas and assuming Gaussian probability distribution. This allowed the authors of Billeh et al. (2020) to estimate the widths of the Gaussian connectivity profiles, yielding:

$\sigma_{EE} \simeq 6^\circ, \sigma_{PE} = 5^\circ, \sigma_{SE} \simeq \sigma_{VE} \simeq 5^\circ$, $\sigma_{EP} \simeq 5^\circ, \sigma_{ES} \simeq 4^\circ$. Moreover all inhibitory to inhibitory widths were estimated to be the same as PV to PV, which was measured $\sigma_{PP} \simeq 6^\circ$.

These values are also largely consistent with more recent measures of simultaneous whole-cell patch-clamp in mouse V1 Campagnola et al. (2021). The authors fit a Gaussian to the connection probability as a function of lateral intersomatic distance and report: $\sigma_{EE} = 6^\circ, \sigma_{IE} = 5^\circ, \sigma_{EI} = 5^\circ, \sigma_{II} = 6^\circ$, where $I$ is unspecified inhibition.

Fig. SF31 reports the values of the cortico-cortical widths of the connection probabilities/effective strengths for different cell-class pairs that we draw from this review of the literature: $\sigma_{EE} = \sigma_{EL} = 7^\circ$, $\sigma_{SE} = 8^\circ$, $\sigma_{PE} = \sigma_{VE} = \sigma_{P4} = 5^\circ$, $\sigma_{EP} = 5^\circ$, $\sigma_{ES} = 4^\circ$, $\sigma_{II} = 6^\circ$ where $I$ indicates all inhibitory cell types $I = \{P, S, V\}$. Finally, the projection width in orientation space is taken from a fit of the measures in Rossi et al. (2020) and its value is $\lambda = \frac{\pi}{6}$ (same for all cell types).

#### SS2. EXPERIMENTAL RATE FIELDS

##### A. Data preprocessing

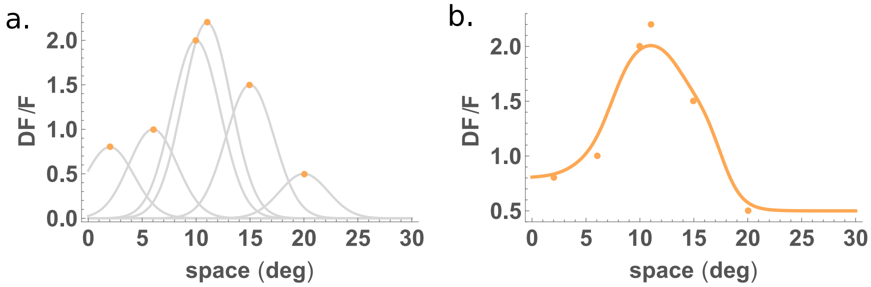


FIG. SF1. Sketch of the method to derive the rate field from the data points. **a**. Fictitious data points (orange dots) and Gaussian function (gray line) describing the probability that the recorded cell had a particular RF center (horizontal coordinate), considering the error estimate of $\sigma_{err} = 5°$. **b**. fictitious data points (orange dots) and relative rate field calculated as explained in the STAR methods.

##### B. Rate fields calculated with two different preprocessing procedures

Here we present the rate field as obtained from the dataset in Keller et al. (2020a). We test the robustness of the method described in the STAR methods by comparing it with a different, more rough procedure: we divide the retinotopic space in bins, then we take the median of the response of all the cells that fall within each bin (according to their RF measure). This alternative procedure allows to consider the error on the estimate of the response (standard error of the mean calculated on the set of cells falling in each bin), but not the error on the estimate of the RF position. Also we remark that there exists a non negligible dependence of the spatial profiles calculated in this latter way (and consequently their fits) with the number of bins chosen (data not shown). This dependence can be imputed to the error on the estimate of the RF position. We define a linear transformation between the fluorescence response and the firing rates, so that the rate fields are expressed in Hz, as in Keller et al. (2020b).

Note that the baselines of the inverse rate fields are chosen in accordance with those of the classical rate fields, based on continuity between the responses to the largest direct stimulus and the smallest inverse stimulus, for the cells that are furthest away from the stimulus center. Note that the responses to the largest classical stimuli are far from being uniform in space (as they would be if the stimulus was effectively covering the whole visual field of the animal). Overall the corresponding profiles obtained through the two preprocessing procedures are satisfactorily similar for both direct and inverse stimulus condition (respectively represented in Fig.SF2 and Fig.SF4).

##### C. Weak increase in activated V1 area with increasing stimulus size

a.

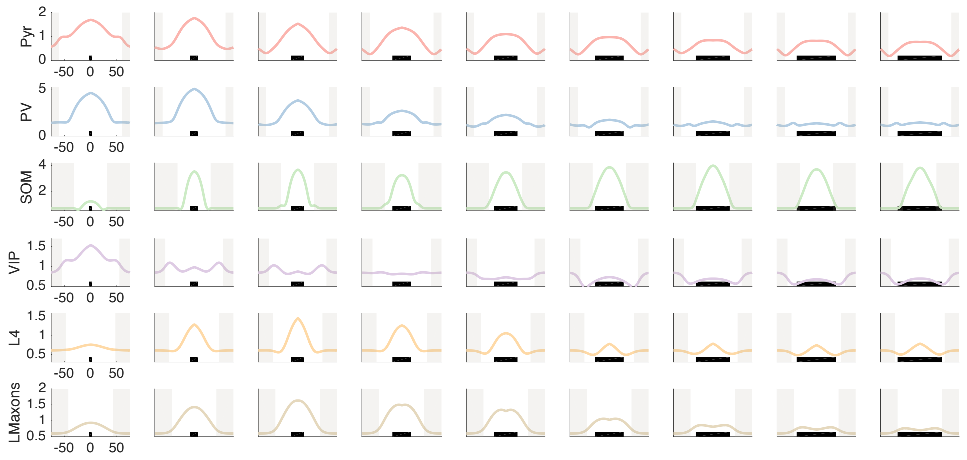


b.

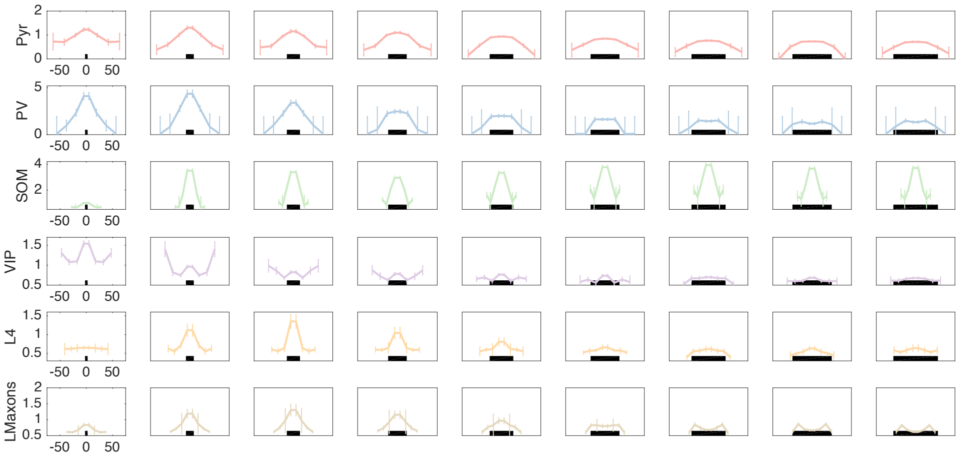


FIG. SF2. **a**. Rate fields estimated with the method presented in STAR Methods. The shaded areas correspond to regions of low confidence, where data points are rare or absent and rate fields are estimated by imposing baseline activity (under the assumption that cells at large distances from the stimulus do not respond to the stimulus). The black bars at the bottom of each panel represent the stimulus size. **b**. Rate fields estimated through the median of the response of the cells in each spatial bin, for all cell types and all stimulus sizes, in the classical stimulus condition.

a.

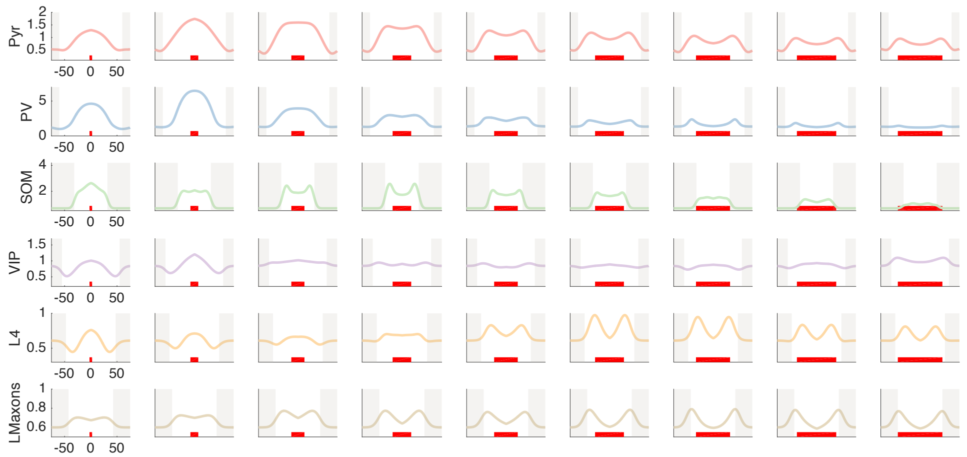


b.

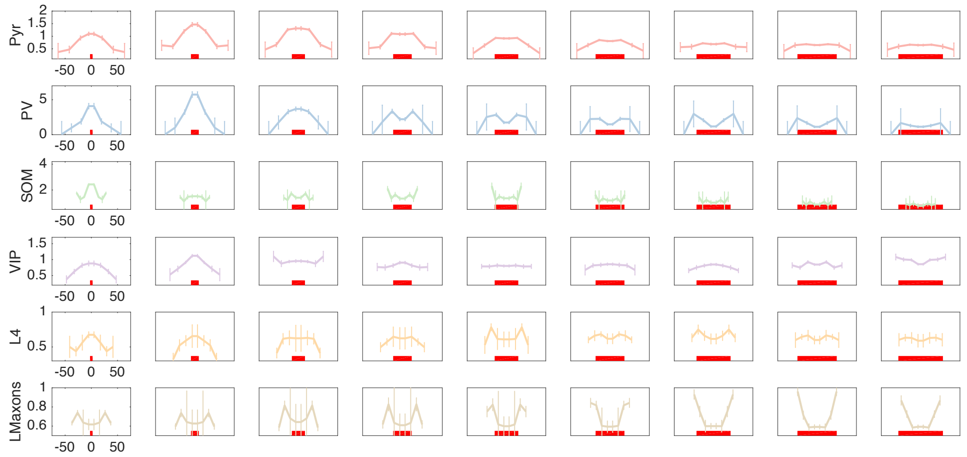


FIG. SF3. Same as FigSF2 for inverse response. The red bars at the bottom of each panel represent the inverse stimulus size (diameter of the 'hole').

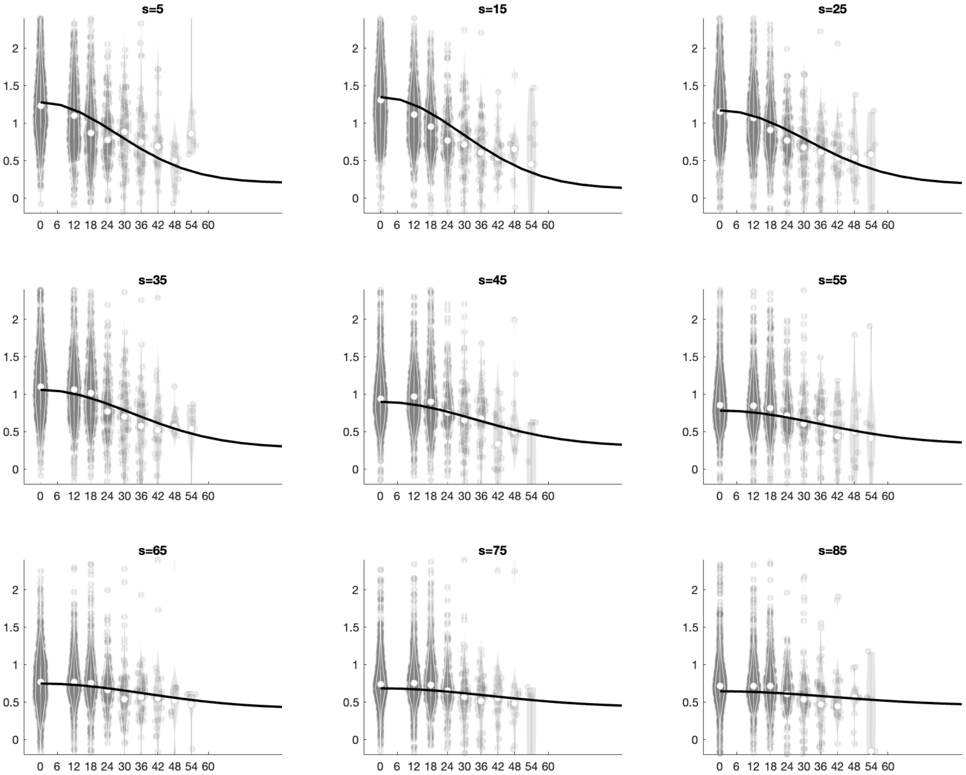


FIG. SF4. Spatial profiles of response data of Pyr cells for different stimulus sizes. The data is divided in bins according to the retinotopic location (x axis), the violin plots represent the distribution of cells falling in each bin, the white dot is the median of the distribution and the black full line is half Gaussian for comparison, whose width $\sigma$ grows with stimulus size with slope 0.13 (fitted value for width scaling of Pyr, black line in Fig.SF6b). The expected slope if the activity patch grew as fast as the stimulus size would be 0.5.

#### SS3. ARE ALIGNED AND OFFSET LM NEURONS INTRINSICALLY DIFFERENT?

The question is whether offset LM neurons and aligned (or centered) ones are intrinsically different classes of neurons (as suggested in Keller et al. (2020a)) or they are the same class of neurons. If they were intrinsically different classes of neurons, their properties (e.g. connectivity profiles) could be different and/or the response to a retinotopically matched stimulus could be different. On the contrary, if they were the same population, their response properties to (precisely) retinotopically matched stimuli would be the same, and a difference in response would be imputable to a difference in retinotopic position relative to the stimulus.

In support of the existence of 2 different populations in LM one could argue that the response to the largest stimulus is different (Figures 5g and 5i in Keller et al. (2020a)). Considering that the largest stimulus is virtually a full field grating, then a difference in the response indicates that they have a different response to the same stimulus.

In other words, if the stimulus was homogeneous in space (e.g. grating covering the full visual field of the animal), the mismatch of responses of aligned and offset neurons would imply that they have intrinsically different response properties, thus they are 2 different populations, potentially with 2 different connectivity profiles.

Nevertheless, the largest stimulus size is a disk of diameter 85° and the offset boutons with the furthest away receptive field considered are at a distance of 42° from the center of the stimulus. This means that at least some of the offset boutons' RF is close to the border of the largest stimulus. Therefore the premise that the largest stimulus size is a full field grating is not a satisfactory approximation.

In order to address the question more quantitatively, we apply a clustering algorithm (Uniform Manifold Approximation and Projection, UMAP) to the responses. An embedding in 2D is found by searching for a low dimensional projection of the data that has the closest possible equivalent fuzzy topological structure (all the details on the procedure can be found in uma). Once the clustering is done we plot the scatter plot of all data points in its 2D embedding. If there were $n$ different population (with $n$ different response patterns), the plot should show $n$ well-separated sets of points (as easily recovered when using the MNIST dataset, data not show). After the clustering algorithm is applied to the data, only for interpretation purposes, we color each point according to its label (Fig. SF5).

When we apply the algorithm to the response data of all boutons, the number of clusters found is $n = 1$, i.e. there is only one cloud of points. When we color the points according to their label (aligned, offset or unknown), we notice that the x axis is correlated with the identity of the boutons (aligned boutons –in blue– tend to be on the right). This is consistent with a situation in which the responses change with continuity, the aligned boutons being in one end of the spectrum. This is consistent with the hypothesis that the neurons differ only in their retinotopic location relative to the stimulus, but not in their intrinsic properties. We then add another color to show the boutons that are excluded in Fig.5i in Keller et al. (2020a), i.e. the boutons that do not have significant response to the inverse stimulus.

What we observe is that the neurons excluded from Fig 5i in Keller et al. (2020a) are the offsets and unknowns that are closest or overlapping with the aligned ones (colored pink, labeled 0 in Fig.SF5). This suggests that the selection rule in Fig. 5i is effectively creating two "artificial" clusters, by removing the offset boutons whose response is inbetween the two ends of the spectrum. In fact, if we repeat the clustering analysis with the same selection criteria as Fig. 5i, two (non completely separated) clusters arguably appear, corresponding to aligned (blue) and offset+unknown (green+yellow) boutons.

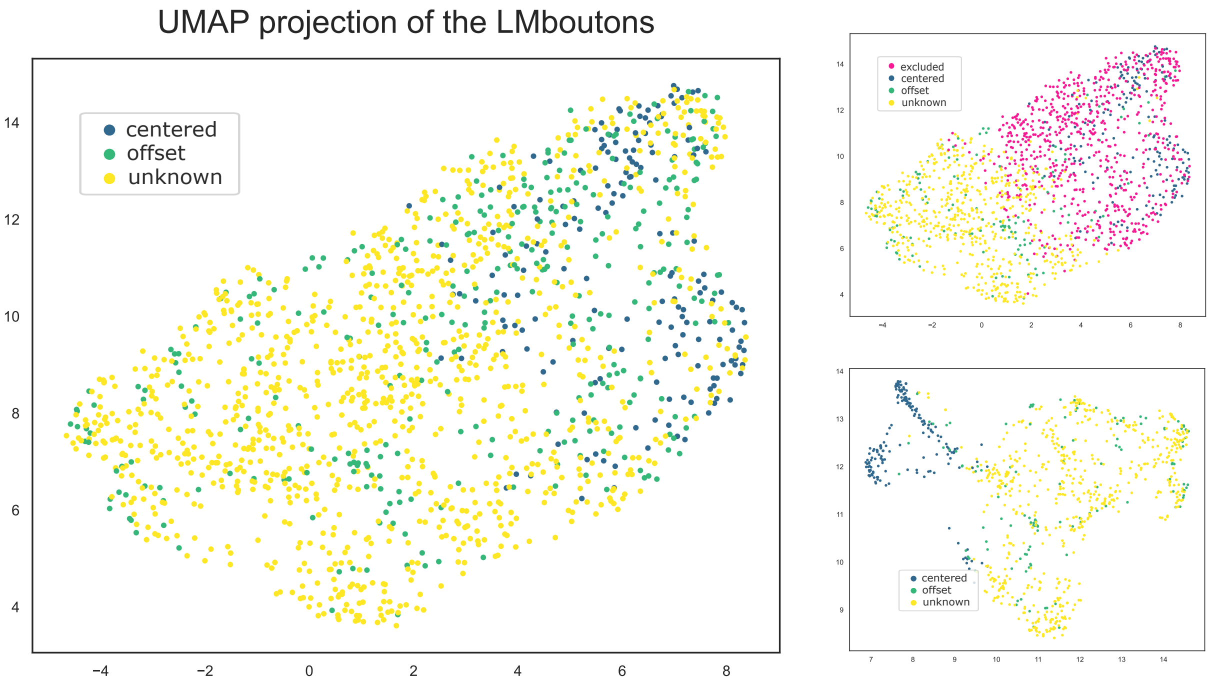


FIG. SF5. UMAP clustering procedure identifies no clustering in the data, as the data points are disposed in one unique cloud. Unknown cells are those for which a receptive field could not be identified, excluded cells are those showing non significant inverse response, excluded based on selection criteria specified in Keller et al. (2020a). In the bottom right panel a certain degree of clustering appears as an effect of the exclusion of the pink (excluded) data points.

#### SS4. GAUSSIAN APPROXIMATION FOR THE RETINOTOPIC SPATIAL PROFILE OF THE RATE FIELDS

We fit the experimental rate fields for each stimulus size in the classical stimulus condition with a Gaussian function (see STAR methods). We report the goodness of fits applied to both preprocessing procedures in Fig.SF6a (dots and diamonds).

The values of $R^2$ found are close to 1 consistently across cell types and stimulus sizes, except for VIP cells (compare with Fig.SF2a and b, fourth row) and for LM cells for large stimuli, where values of $R^2$ close to 1 can be recovered if the profiles are fit with a difference of Gaussian functions. Nevertheless, for simplicity, we consider this change in geometry a second order effect, to be tackled elsewhere. We also notice that the fits are consistently more accurate for the method discussed in the STAR Methods with respect to the one introduced above (based on a binning of the retinotopic space).

Finally, in Fig.SF6b we report the width of the Gaussian fits as a function of stimulus size, together with a linear fit of their trend as a function of stimulus size and the expected trend, assumed in previous theoretical works Dipoppa et al. (2018); Rubin et al. (2015); Obeid and Miller (2021).

In Fig.SF8a we report the goodness of fits to the rate fields obtained through both preprocessing procedures in the inverse stimulus condition. Overall the profiles are fitted quite well by a difference of Gaussians, especially when the method discussed in the STAR Methods is used. In Fig.SF8b we report the outer scaling, i.e. the value of the fit of the $\sigma = \sqrt{v}$ parameter of the Gaussian with positive amplitude, i.e. $\sigma_{iA}$.

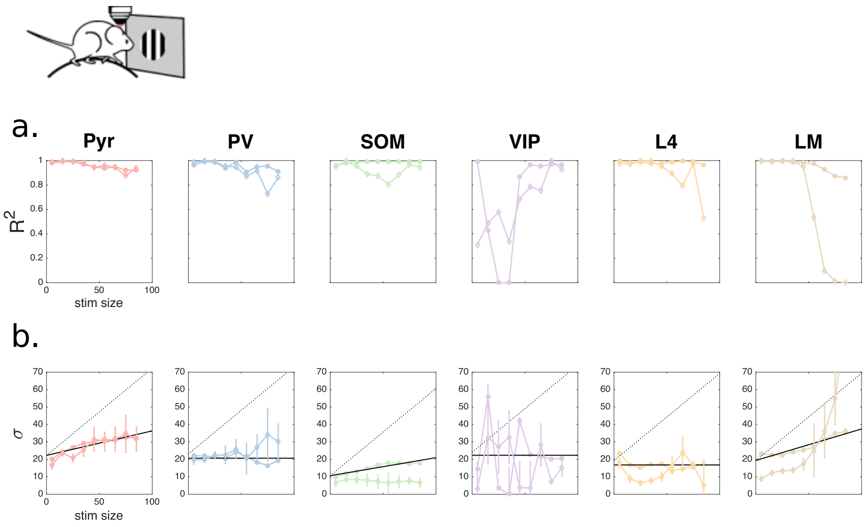


FIG. SF6. Goodness of Gaussian fit evaluated through $R^2$ (panels **a**) and $\sigma$ (panels **b**) parameter of the fitted Gaussian functions for all cell types and stimulus sizes, in the classical stimulus condition. Full dots represent fits of the rate fields obtained from the method presented in the STAR Methods, full diamonds represents fits of the rate fields estimated through the binning procedure described in Section SS2 B. The error bars are 95% confidence intervals on the fit. Solid black lines are linear fits of the dependence of $\sigma$ on the stimulus size (for the occupancy method) and dotted lines correspond to $\sigma(s) = s/2 + \sigma(0)$. We refer to the slow growth of the solid versus the dotted line as weak scaling of the rate fields width. In particular we note that $\sigma'_L(s) = 0 \pm 0.05$ and $\sigma'_M(s) = 0.18 \pm 0.05$

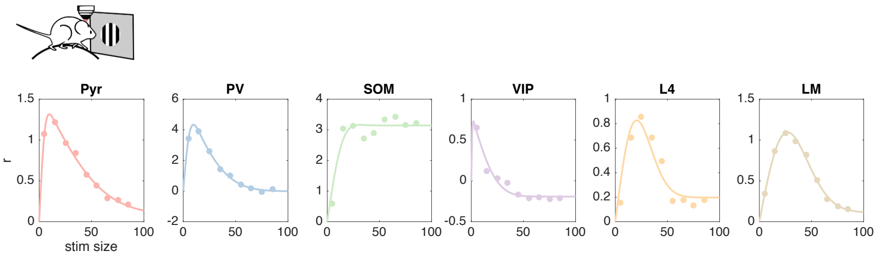


FIG. SF7. Amplitudes of the of the fitted Gaussian functions for all cell types and stimulus sizes, in the classical stimulus condition, preprocessing as in the STAR Methods. Solid lines are fits with a difference of error functions (see STAR Methods).

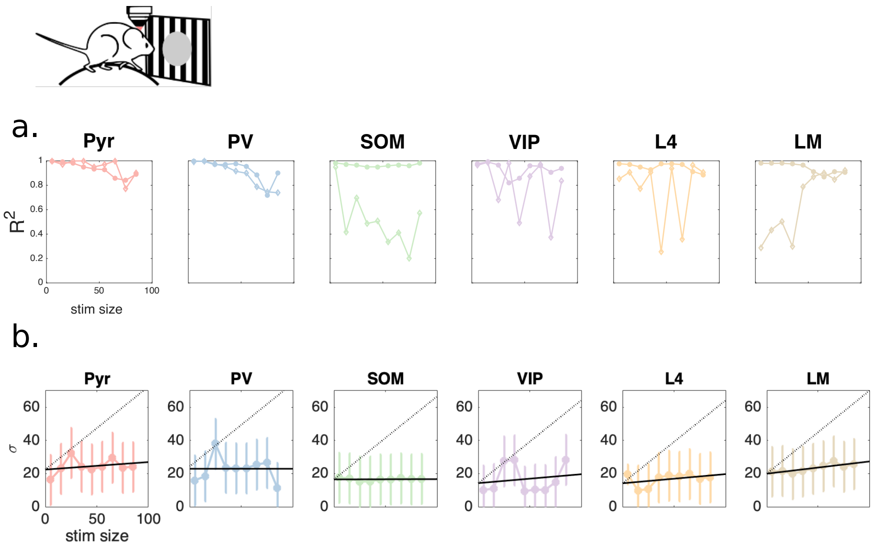


FIG. SF8. Same plot as Fig.SF6 for inverse stimuli and double Gaussian fits. Panel **a**. represents the goodness of Gaussian fits and panel **b**. the $\sigma$ parameter of the positive-amplitude Gaussian function of the fit, for all cell types and inverse stimulus sizes. The error bars are larger than in Fig.SF6b because of the increased number of degrees of freedom. Full dots represent fits of the rate fields obtained from the method presented in the STAR Methods.

##### A. Parametrization of classical and inverse size tuning curves

Fig.SF9 (Fig.SF10) shows the classical (inverse) size tuning curves described by the functional form reported in the STAR methods, as well as the data and the fitted rate fields. Classical size tuning curves are largely consistent with previous reports Dipoppa et al. (2018); Angelucci et al. (2017); Li and Young (2021). The rate field of SOM neurons in L2/3 shows a lack of surround suppression, in agreement with previous results Adesnik et al. (2012). However, a minor amount of surround suppression has been reported in Dipoppa et al. (2018) for SOM cells whose CRF distance from the stimulus center is very small. It is possible that the precision in the measure of SOM CRF mapping in the dataset of Keller et al. (2020a) may not suffice to resolve this small amount of surround suppression.

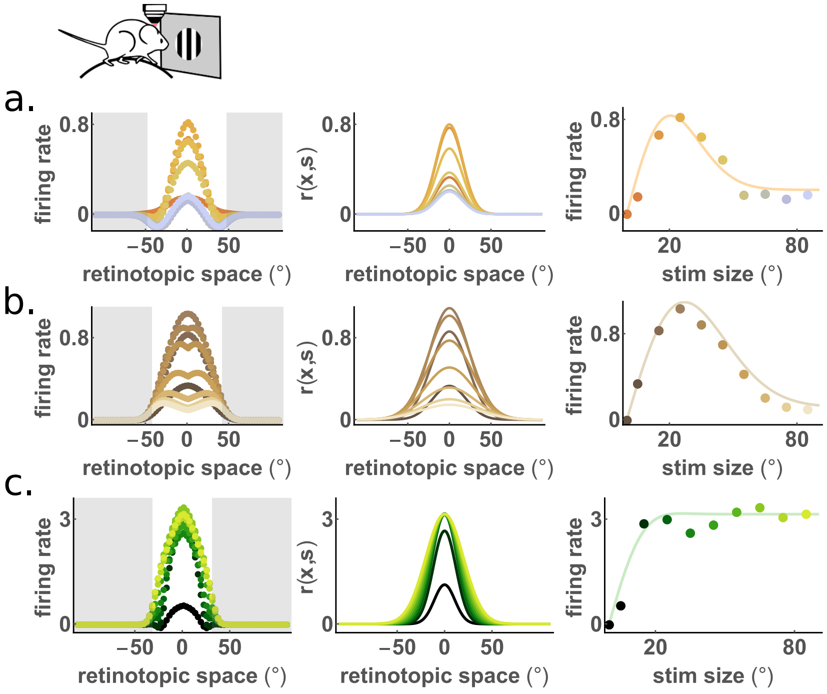


FIG. SF9. Classical stimulus condition. Experimental (left panels, same as SF2 and SF4) and parametrized (center panels) rate fields $r_A^{(i)}(\mathbf{x}, s)$, for direct comparison with figures in the main text. Right panels show the rate field for aligned cells as a function of size (full circles) and the parametrization of the size tuning curves, i.e. a (difference of error functions) fit to the parametrized rate fields. **a**. L4, **b**. LM, **c**. SOM.

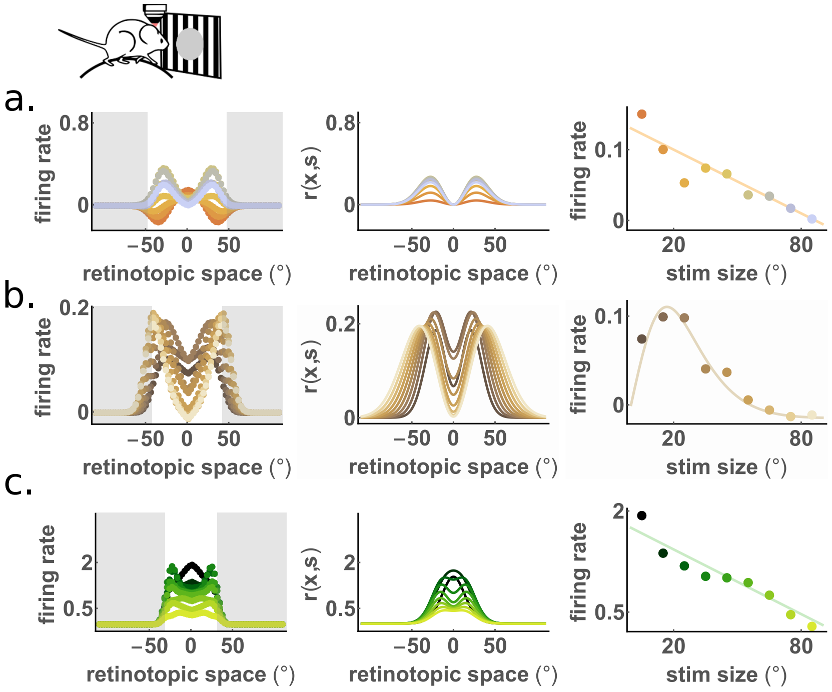


FIG. SF10. Same as Fig.SF9 for inverse stimulus condition. Experimental (left panels) and parametrized (center panels) rate fields $r_A^{(i)}(\mathbf{x}, s)$, for direct comparison with figures in the main text. Right panels show the rate field for aligned cells as a function of size (full circles) and a (linear or difference of error function) fit to these points. **a**. L4, **b**. LM, **c**. SOM.

#### SS5. FROM LM TO HVAS

The firing rates recorded from LM boutons by Keller et al, Keller et al. (2020a) were on average (across space) several times higher in the classical stimulus condition compared to the inverse one. At the same time the firing rates recorded in L4 Pyr cells were also very low for the inverse stimuli, while the firing rates for L2/3 Pyr+PV population were as large as the ones recorded during classical stimuli presentations. Finally VIP firing rates are comparable for the classical vs inverse stimuli and SOM is somewhat smaller for inverse stimuli, yet its response decreases with inverse size (the opposite of what would be needed to generate inverse surround suppression). If the net incoming current to a L2/3 unit of the Pyr+PV population is taken to be the sum of the lateral, feedforward (from L4) and feedback (from LM) currents, this would be much more excitatory in the classical case compared to the inverse, resulting in a much larger L2/3 firing rate for classical stimuli, in contrast with the experimental observation. Nevertheless L2/3 cells receive feedback input not only from LM, but from all HVAs Keller et al. (2020a); Siegle et al. (2021). Moreover Keller et al. Keller et al. (2020a) show that optogenetically silencing an individual HVA, suppress classical and inverse L2/3 response by different amounts (Extended Data Figure 9 ibidem). In particular areas M, AL, LM, LI and P all seem to contribute more to inverse than to classical response. This suggests that the total feedback input to L2/3 (proceeding from all HVAs) might be of the same average magnitude for the classical and inverse stimuli. This allows us to recover the observed relative amplitude of L2/3 responses. More precisely, we describe the total (HVAs) feedback current with a rate field that has the same spatial profile as the one recorded from LM, but we introduce a parameter $\eta$ that rescales the input from LM in the inverse condition. We fit $\eta$ to recover the amplitude of the inverse response.

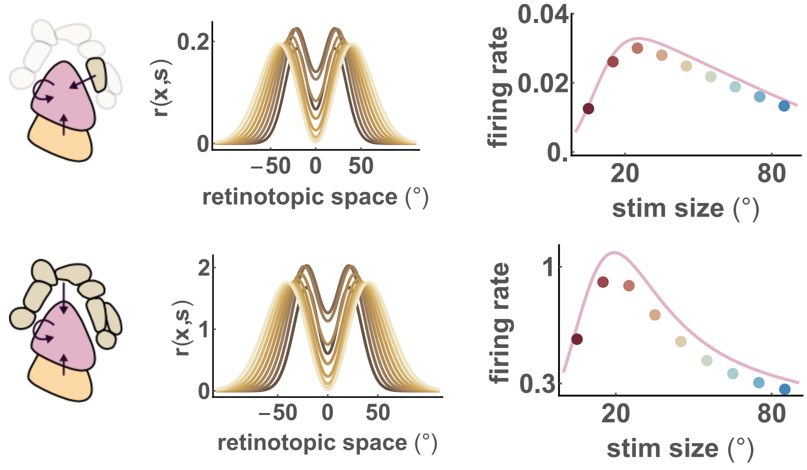


FIG. SF11. Top: minimal model receiving feedback input only from LM. Bottom: minimal model receiving feedback input from HVAs. Left panels: sketch. Center panels: the rate field of LM (top) is obtained as explained in the STAR Methods, whereas the rate field of HVAs (bottom) is assumed to have the same profile, but scaled by a constant $\eta = 9$. This scaling constant is determined in such a way to reproduce the amplitude of the inverse rate field of L2/3. Right panels: size tuning curve for the recurrent layer L2/3 for LM input (top) or HVAs input (bottom).

#### SS6. RATE FIELDS FOR THE JOINT POPULATION OF PYR AND PV

We show the rate field of the joint population of Pyramidal and PV recorded cells for comparison with the results of the minimal model in Fig.3 and 5 in the main text. The joint population is built by taking the average rate field of both populations and then weighting each of the two by the appropriate density of cells as reported in Tremblay et al. (2016).

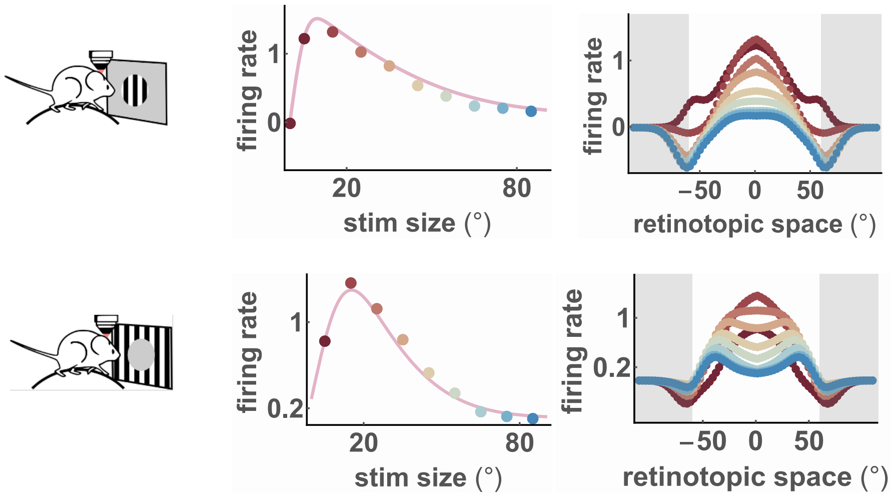


FIG. SF12. Experimental size tuning curves and rate fields for the joint population of Pyramidal and PV L2/3 cells, for classical stimuli (top panels) and inverse stimuli (bottom panels). Left: size-tuning curve, i.e. rate field in the origin as a function of stimulus size, full line is a fit with a difference of error functions; right: rate field for various stimulus sizes, color-coded as in the left panel.

This joint population is effectively either excitatory or inhibitory, depending on whether Pyramidal or PV cells dominate. Since the cortex has been found to be inhibition stabilized (Denève and Machens, 2016; Sanzeni et al., 2020; Palmigiano et al., 2020; Isaacson and Scanziani, 2011; Okun and Lampl, 2008; Bos et al., 2020), we choose it to be effectively inhibitory, thereby preventing the system to diverge when external excitatory inputs are injected. The recurrent synaptic weight that we use is then negative, although the results presented in this work still hold with a small positive weight (see Supplementary Section SS7 B).

The comparison between the recordings shown here and the results of the minimal model (Fig.3 and 5 of the main text) should be taken as an estimate of the implications of the simplifying assumptions that the minimal model is adopting. Additional inputs, cell-specific input-output functions, non-Gaussian profiles, systematic errors in the estimate of the firing rates from calcium imaging, synaptic plasticity are some of the factors plausibly responsible for the mismatch: the goal of the model is to obtain a versatile description of surround suppression, rather than reproducing in detail the experimental size tuning curve (see Discussion Section in the main text). In particular we notice that in the data, around the region of positive (larger than baseline) activity there exists a region where cells respond below baseline. This reminds us of the spatial profile of response to optogenetic perturbations (see e.g. Chettih and Harvey (2019)). Moreover similar profiles with resonant spatial frequencies have been found for an input roughly equal across the activated region ('pillbox-shaped' input) in an SSN model Rubin et al. (2015).

##### A. Comparison with a different dataset

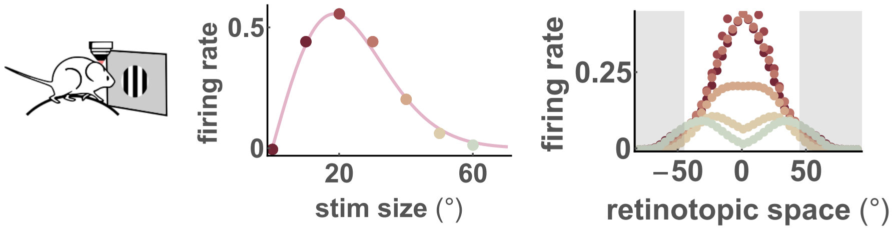


FIG. SF13. Experimental rate fields, and size tuning curves for the joint population of Pyramidal and PV L2/3 cells, for data in Dipoppa et al. (2018), classical stimulus condition. Note the change in concavity in the center of the rate field for large stimuli. To be compared with Fig. 3l-m in the main text. The data for PV and Pyr as a function of space and stimulus size were plotted in Supplementary Fig. 11 in Dipoppa et al. (2018). The change in concavity is visible as the brighter tail in the bottom right part of the heatmaps, but it was not discussed in detail.

### SS7. THE MINIMAL MODEL WITH INPUTS FROM L4 AND HVAS

#### A. Validation of the minimal model

To solve the self-consistency equations for the spatial SSN (STAR Methods, Eq. 2) we took the ansatz:

$$u(\mathbf{x}) = \alpha(\mathbf{x})G(\mathbf{x}, v_u), \tag{1}$$

with the constraint that $\alpha(\mathbf{x})$ is a slowly varying fucntion of its argument. Here we show that the solution that we find for $\alpha(\mathbf{x})$ is actually consistent with the assumption that it varies slowly with $\mathbf{x}$. Analytically, if we consider only one external input $I(\mathbf{x}) = I_0 G(\mathbf{x}, v)$ (as in STAR Methods) and we restrict to one spatial dimension for illustration purposes, we find:

$$\frac{\alpha'(x,0)}{\alpha(x,0)} = \frac{x(-3+\sqrt{\frac{4v}{v_r}+1})}{v} = \mathcal{O}\left(\frac{x}{v}\right) \tag{2}$$

which is very small for the values that are relevant in this context, i.e. $|x| < 50$, $v \sim 20^2$, as shown in Fig.SF14.

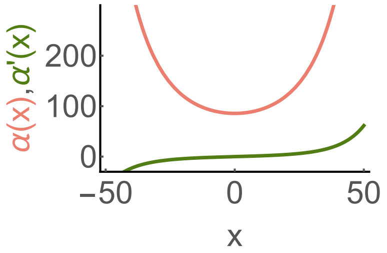


FIG. SF14. Plot of the function $\alpha(x, y = 0)$ together with its derivative $\alpha'(x) = \partial\alpha(x, y = 0)/\partial x$, showing that $\alpha'(x, 0) \ll \alpha(x, 0)$, i.e. the function is slowly varying. The parameter used are the same ones used throughout the main text; the external input chosen is the feedforward input, with $v = 15°$.

In what follows we present a set of results for the minimal model with anatomically-realistic length scales and experimental input rate fields from L4 and HVAs, obtained as decribed in the STAR Methods.

#### B. The joint population of Pyr+PV is stable if inhibition dominates

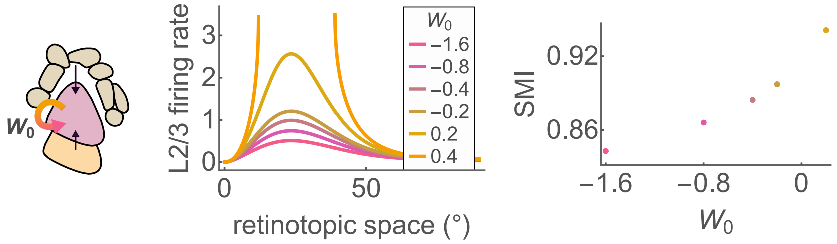


FIG. SF15. Since we consider the recurrent population to be a joint population of Pyr and PV cells, the recurrent connection strength $W_0$ can be either positive or negative, depending on whether Pyr or PV dominates. Throughout this work we choose a negative value $W_0 = -0.4$, since we want the network to be well within the inhibition stabilized regime. Here we show the change of size tuning curve as a function of $W_0$, that can be compared with experiments of activation or silencing of PV cells in L2/3 Nienborg et al. (2013). Note that i) when PV contribution is too weak, i.e. $W_0$ is too large the network is unstable (see Eq.(12) in the STAR Methods), ii) for more negative $W_0$ surround modulation decreases, which means that activation of PV cells leads to a decrease in SMI.

##### C. Dependence of the firing rate of L2/3 on the FF and FB connections strengths

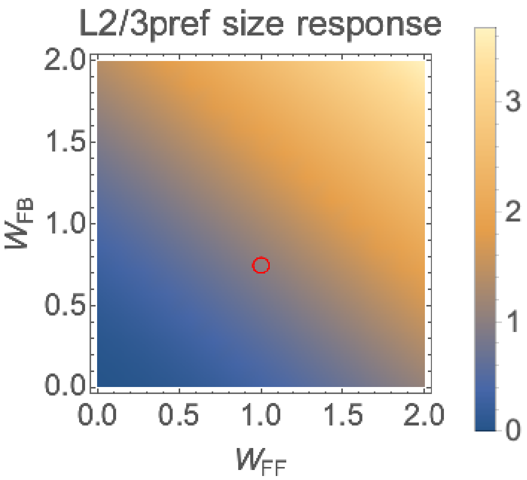


FIG. SF16. The color gradient represents the minimal model response of L2/3 as a function of the feedforward and feedback connection weights. The red circle indicates the parameters chosen in the main text and unless specified otherwise.

#### SS8. ANALYTICAL INSIGHTS ON CLASSICAL SIZE TUNING CURVES WITH NON-NORMALIZED INPUT CURRENTS AND COMPARISON TO PREVIOUS MODELS OF SS

In Eq. (26) and (28) of the STAR Methods we show that when the inputs are given by the convolution of the firing rates of the input layers with the inter-layer connectivity, the amplitude of the input is proportional to the width of the generating rate field $I^c(\mathbf{x},s) = I_0^c(s)G(\mathbf{x},v(s)) \propto 2\pi v_r(s)r(s)G(\mathbf{x},v(s))$ and $v_r(s)$ and $v(s)$ are related (see Eq. 25 of the STAR Methods). This is a difference with respect to reference models of classical surround suppression in the literature (e.g. Rubin et al. (2015); Obeid and Miller (2021)), where the input currents are not derived from rate fields.

To establish a comparison with existing models, and to simplify the analytical calculations, in this Section we first study classical surround suppression in the case in which $I(\mathbf{x},s) = I_0(s)G(\mathbf{x},v(s))$ and $\frac{dI_0(s)}{dv(s)} = 0$ and then show that the simpler case studied analytically provides insights that hold in the more involved case of our main study, where the input currents are derived from rate fields.

Note that when performing calculations for the minimal model, the dependence on stimulus size is not explicitly written (see STAR Methods section). Instead, it is encapsulated within the parameter $I_0$. Consequently, this distinction does not alter the functional form of the solution; formally it only involves a redefinition of the parameter $I_0$. Nonetheless, from a quantitative perspective, when we consider $I_0^c(s) \propto 2\pi v_r(s)$ and given that $v_r(s)$ grows –however weakly– with stimulus size, the input current to the center cells becomes larger for large stimuli because of the contributions of the offset cells of the feedforward layer.

##### A. Dissecting Classical surround suppression

In this Section we show that classical surround suppression in partially inherited from surround suppression of the input layer(s) and partially due to the width of the input rate field growing with stimulus size. Surround suppression corresponds to a size tuning curve that decreases with increasing stimulus size, i.e. $\frac{r(s)}{ds} \equiv r'(s) < 0$. The dependence on the stimulus size enters in the rate field of the recurrent layer through the amplitude of the input $I_0(s)$ and its width, described here by the input variance $v(s)$. In the simplified case considered here ($\frac{dI_0(s)}{dv(s)} = 0$) we can consider one of these contributions at a time and we will show that:

$$\begin{aligned} &\text{if } \; v'(s) = 0, \text{ then } r'(s) < 0 \Leftrightarrow I_0'(s) < 0 \\ \text{if } \; I'(s) = 0 \text{ and } &\{W_0 < 0, I_0 > 0\}, \text{ then } r'(s) < 0 \Leftrightarrow v'(s) > 0 \end{aligned} \tag{3}$$

Since the transfer function is non-negative and monotonic, the size tuning properties of the firing rate are the same as the size tuning properties of the associated total current field:

$$u(\mathbf{x}=0,s) = \frac{\pi(v(s)+v_{ru}(s))}{W_0}\left(1-\sqrt{1-\frac{2I_0(s)W_0}{\pi(v(s)+v_{ru}(s))}}\right)\frac{1}{2\pi v(s)} \tag{4}$$

If $v'(s) = 0$, we find

$$u'(\mathbf{x}=0,s) = \frac{I_0(s)'}{2\pi v\sqrt{1-\frac{2W_0I_0(s)}{vv_{ru}}}} \tag{5}$$

where one easily reads that $u'(s) < 0 \Leftrightarrow I'(s) < 0$ which means that the recurrent layer is surround suppressed when the input layer is surround suppressed.

Conversely, if $I_0'(s) = 0$ (i.e. $I_0(s) = I_0$) we have:

$$\begin{aligned} u'(\mathbf{x}=0,s) &= t_1(s)+t_2(s)+t_3(s); \\ t_1(s) &= v_{ru}(s)\frac{\pi}{W_0}\left(-1+\sqrt{1-\frac{2I_0W_0}{v(s)+v_{ru}(s)}}\right)v'(s) \\ t_2(s) &= -I_0\frac{(v'(s)+v'_{ru}(s))v(s)}{(v(s)+v_{ru}(s))\sqrt{1-\frac{2I_0W_0}{v+v_{ru}}}} \\ t_3(s) &= v(s)\frac{\pi}{W_0}\left(-1+\sqrt{1-\frac{2I_0W_0}{v(s)+v_{ru}(s)}}\right)v'_{ru}(s) \end{aligned} \tag{6}$$

Moreover taking into account that $2v_u^{-1} = v^{-1} + v_{ru}^{-1}$ (see STAR Methods) we find:

$$v'_{ru}(s) = \frac{v_r v'(s)}{\sqrt{v_r^2 + 4v_r v(s)}} > 0 \tag{7}$$

therefore it is easy to see that for $v'(s) > 0$ and $W_0 < 0, I_0 > 0$:

$$\begin{aligned} t_1(s) < 0, t_2(s) < 0 \\ t_3(s) > 0 \end{aligned} \tag{8}$$

Finally we can show that $|t_3| < |t_1|$. In fact we just have to show that $vv'_{ru} < v_{ru}v'$. Using Eq.7 we easily get to the inequality:

$$2vv_r < 4vv_r + v_r\sqrt{v_r^2 + 4vv_r} + v_r^2 \tag{9}$$

which is always true, given that the r.h.s has 3 positive terms, the first of which is already larger than the l.h.s. (Fig.SF17). Moreover it is easy to see that $\lim_{\sigma(s)\to\infty} u(s) = 0$.

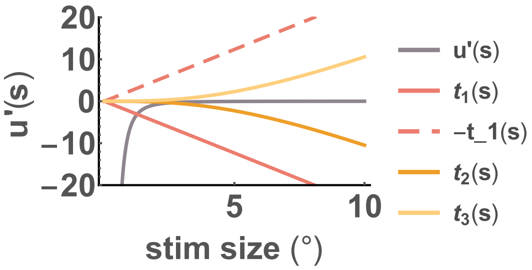


FIG. SF17. Plot of the terms $t_1(s), t_2(s), t_3(s)$ in Eq. 6, for $I_0 = 5, W_0 = -1, v_r = 7^2$ in full lines and $-t_1(s)$ in dashed line. The plot shows graphically that $|t_3| < |t_1|$ and that $u'(s) \lesssim 0$.

Thus we have shown that in this system, even if the input is not surround suppressed at all, the recurrent layer can develop surround suppression. Previous models for classical surround suppression described stimuli of increasing size with an input current of increasing width. Here we showed in our setup that our solution is consistent with this result. Nonetheless, our data analyses for feedforward and feedback inputs shows that the input currents width vary only marginally, while their amplitude vary much more significatively, thus we argue that surround suppression in L2/3 is mostly inherited from its input layers.

##### B. Classical surround suppression with non-surround suppressed input rate fields

Here we study numerically the case of one non-surround suppressed excitatory population in the feedforward layer ($\frac{dI_0(s)}{dv(s)} \neq 0$). We want to briefly study the parameter regime for which surround suppression can be imputed to the mechanism of recruiting more lateral inhibition alone. We find that a faster increase in the input width with stimulus size supports classical surround suppression. However, this only holds when the width of the connections from the input layer is small (as in Rubin et al. (2015), modelling cat V1), whereas when it is of the same order of the recurrent projections (as in mouse V1 Rossi et al. (2020)) this mechanism is not sufficient to generate significant surround suppression (see Fig. SF18).

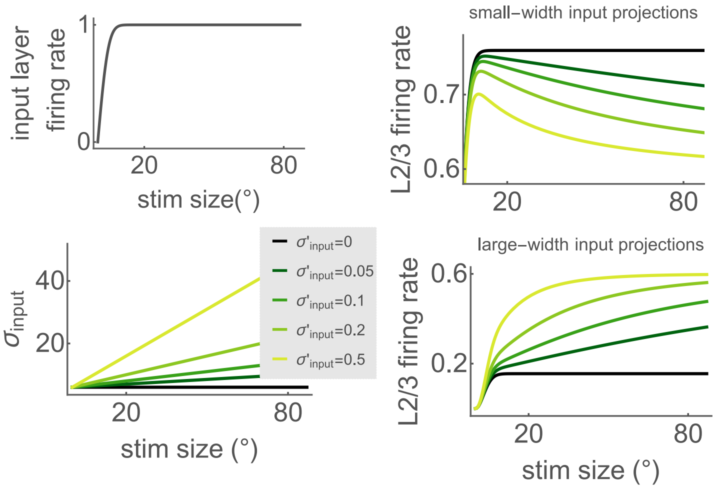


FIG. SF18. Classical surround suppression solely generated by the growth of the width of the input rate field. Top left: firing rate of the centered unit of a generic input layer. Bottom Left: width of the spatial profile of the firing rate of the input layer. Different colors represent different width growths with stimulus size. $\sigma'_{input} = 0.5$ corresponds to the assumption that the input width grows at the same pace as the radius of the stimulus –i.e. half the stimulus size– as in Rubin et al. (2015); Obeid and Miller (2021). Top right: Firing rate of the recurrent layer for different width growths. Significant surround suppression appears when the width of the input layer grows as fast as the stimulus radius. Parameters: $W_0 = -0.4, W_{FF} = 1, \sigma_{rec} = 7, \sigma_{FF} = 1$. Bottom right: Same as above but with input projection width of the same order of the recurrent projections. Same parameters as above but $\sigma_{FF} = 7$.

##### C. Preferred size depends on the preferred size of the inputs

From Eq. 5 we deduce that when $v'(s) = 0$, the preferred size of the recurrent layer is the same as the preferred size of the input layer. Therefore, deviations from this behavior might be imputed to variations in the width of the input field or to convergence of inputs with different preferred size. Moreover from the solution of the minimal model with two inputs, Eq.(13) in the STAR Methods, we can calculate that when the recurrent layer is receiving two inputs with $v'_1(s) = v'_2(s) = 0$, its preferred size $s^*$ will be in-between the preferred sizes of the two inputs. Indeed one has:

$$v_2 I'_1(s^*) = -v_1 I'_2(s^*) \tag{10}$$

where we can read that $\mathrm{sign}[I'_1(s^*)] = -\,\mathrm{sign}[I'_2(s^*)]$, which means that one of the two inputs is increasing while the other one is decreasing (see Fig.SF19).

##### D. Preferred size decreases with contrast

If instead we keep the dependence on stimulus size of one of the two input widths, for instance $v'_2(s) \neq 0$, we find the condition:

$$v_2(s^*) I'_1(s^*) - v_1 I_2(s^*) v'_2(s^*) = -v_1 v_2(s^*) I'_2(s^*), \tag{11}$$

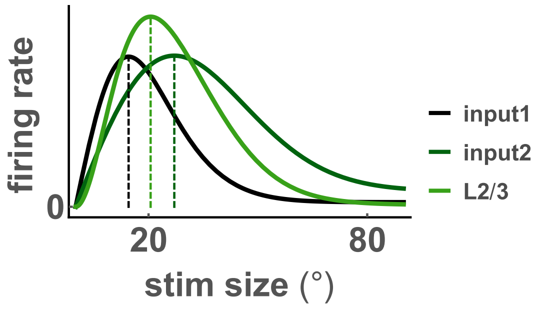


FIG. SF19. For classical stimuli, the preferred size of the recurrent layer lays in between the preferred sizes of its inputs (see Eq.10). Here the width of both inputs rate fields does does not vary with stimulus size.

where one can read that the correction introduced by the dependence $v_2(s)$ reduces the preferred size of the recurrent layer. We hypothesize that this could contribute to contrast dependent classical surround suppression.

In Figure SF20 we show that when the width of the input projections is small with respect to the width of the recurrent connections and the width of the input field grows as fast as the stimulus radius (as in Rubin et al. (2015)), then the preferred size decreases with contrasts. This is in agreement with Rubin et al. (2015). However the model that we are considering here has less ingredients than the one in Rubin et al. (2015) since i) we have only one recurrent population and ii) in (Rubin et al., 2015) the input for stimulus size $s$ was a Gaussian function of space with standard deviation $\sigma_{input}(s)$, whereas in our case because of the convolution with the input layer connectivity it has standard deviation $\sqrt{\sigma_{FF}^2 + \sigma_{input}(s)^2}$.

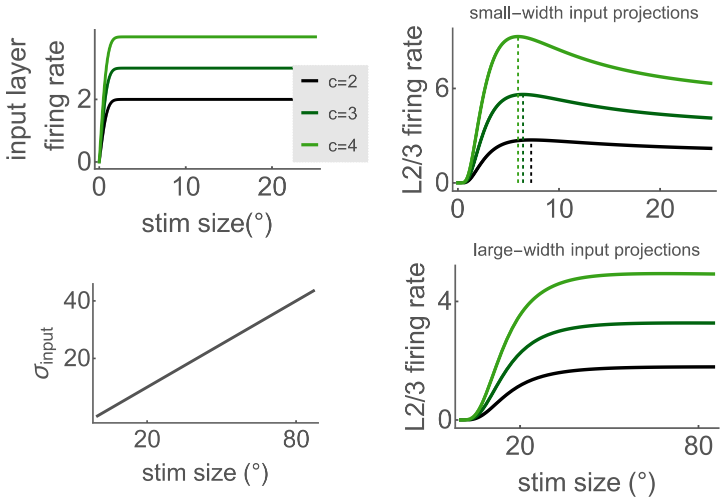


FIG. SF20. Contrast-dependent classical surround suppression solely generated by the growth of the width of the input rate field. Top left: firing rate of the centered unit of a generic input layer for different contrast levels. Bottom left: width of the spatial profile of the firing rate of the input layer ($\sigma'_{input} = 0.5$). Top right: Firing rate of the recurrent layer for different contrast showing that for lower contrast, the preferred size is larger. Parameters: $W_0 = -0.4, W_{FF} = 1, \sigma_{rec} = 7, \sigma_{FF} = 1$. Bottom right: Same as above but with input projection width of the same order of the recurrent projections, not allowing surround suppression (cfr Fig. SF18). Parameters as above but $\sigma_{FF} = 7$.

#### SS9. NOISY NEURAL DYNAMICS ENHANCE THE SURROUND MODULATION OF THE RECURRENT LAYER

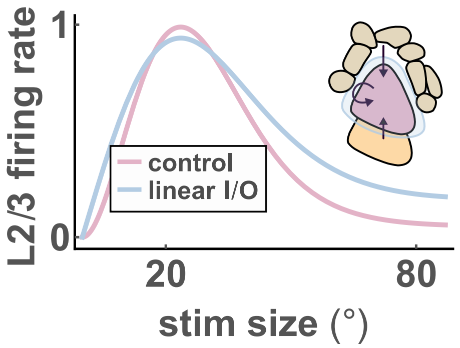


FIG. SF21. Effect of supralinear transfer function for classical stimuli. We show the size tuning curve of the recurrent layer when the inputs are left unaltered, but the rectified supralinear input/output function is substituted by a rectified linear one (light blue). Note that a supralinear transfer function enhances classical size tuning with respect to a linear one.

#### SS10. MODULATIONS INDUCED BY STIMULUS CONTRAST

In this section we ask how the minimal model with anatomically-realistic length scales and experimental input rate fields from L4 and HVAs, can recover contrast-dependent surround suppression.

##### A. Contrast dependent classical size tuning curves

The dataset at hand does not contain data on stimuli with different contrast. One simple way of simulating contrast-varying stimuli is to scale the inputs by a factor that is small for small contrasts. This situation is exposed in Fig.SF22. This model easily reproduces the contrast dependence of the amplitude of response and Surround Modulation Index (SMI, see main text) increases with increasing contrast for a nonlinear model, but not for a linear one (as expected, Rubin et al. (2015); Ahmadian and Miller (2021)). However, with this minimal assumption for contrast-dependence of the inputs, restricting the parameter space to biologically plausible values of the length scales, we cannot reproduce the observation that the preferred size of Pyr cells in L2/3 grows when the contrast decreases Angelucci et al. (2017); Mossing et al. (2021). Based on the arguments exposed above we could argue that contrast-dependence of the preferred size of L2/3 cells could be due to: i) an increase of the preferred size of L4 rate field for low contrasts (observed in data Mossing et al. (2021)), ii) an increase of the preferred size of HVAs rate field for low contrasts (yet not explored experimentally), iii) a decrease in the scaling of (L4 or) HVAs rate field width with stimulus size (yet not explored experimentally), iv) the fact that for low contrasts VIP are more active, thus SOM are less active (shown in experiments and models in Mossing et al. (2021)). We apply one at a time these modifiers to the inputs as a function of contrast and we show that in all cases we reproduce contrast dependent size tuning (see Fig.SF23). This suggests that all these factors may contribute to contrast dependent size tuning.

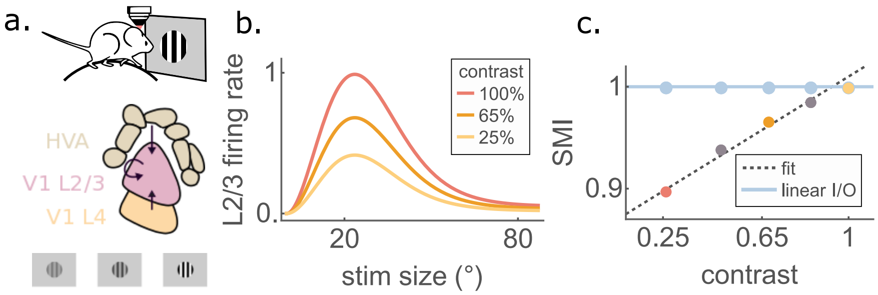


FIG. SF22. **a**. Sketch of the minimal model with the inputs considered. **b**. Comparison between the size tuning curve at maximum contrast and two smaller contrasts, simulated by reducing both feedforward and feedback input by a factor of $C_0 = 0.8$ (65%contrast) and $C_0 = 0.5$ (25%contrast). Dots are simulations, continuous lines are analytic results. **c**. Surround Modulation Index, calculated as the ratio of the center response at a large stimulus size and the center response at the preferred size, as in Mossing et al. (2021). The dots indicate the analytical results of the minimal model, red, orange and yellow dots are in color-code correspondence with panel **b**). The contrast $c$ is defined as $c = \log_2 2C_0$. The agreement with experimental data reported in Fig.1e in Mossing et al. (2021) is excellent. In light blue we plot the SMI for a system with linear transfer function.

#### B. Recovering preferred size dependence on contrast

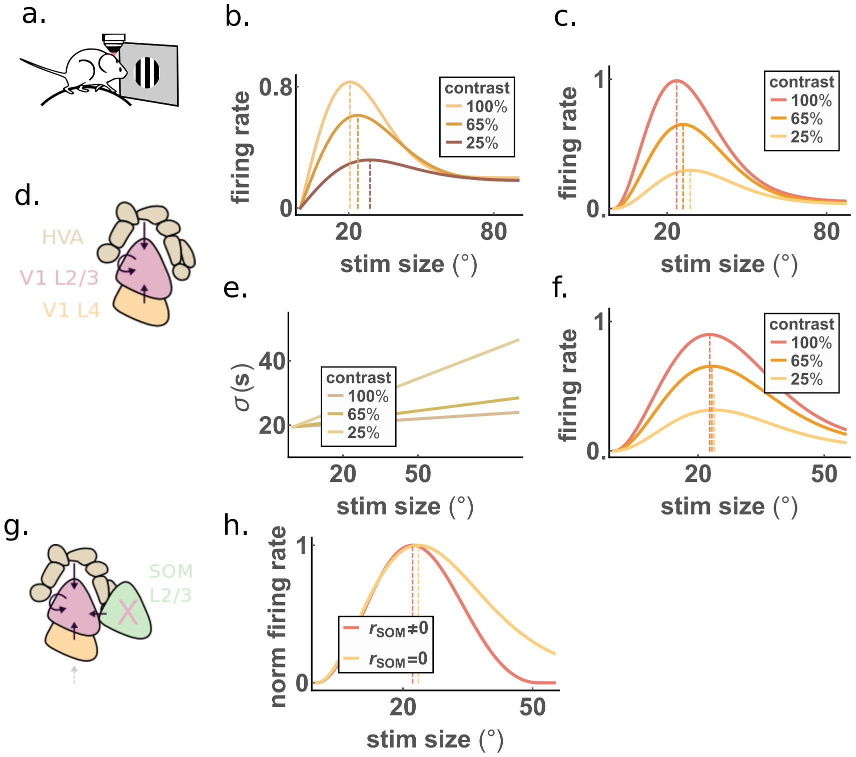


FIG. SF23. **a**. Sketch for classical stimulus condition. **b**. Plausible input from L4 at various contrast levels, showing a larger preferred size for small contrasts (as observed in recordings Mossing et al. (2021)). **c**. size tuning curves of L2/3 with the inputs from L4 as in **b**. A larger preferred stimulus for small contrast is partially inherited from L4. **d**. sketch of the minimal model relative to figures **b**-**f**. **e**. Illustration of another possible contribution to contrast dependent size tuning: for low contrasts the width of the input rate fields grows more rapidly with stimulus size. **f**. size tuning curves of L2/3 with the inputs from HVAs modified according to **e**. A larger preferred stimulus for small contrast could be partially due to a larger scaling of the width for low contrast. **g**. Sketch of the minimal model relative to panel **h**. **h**. Same as Fig. 3n of the main paper. Here we highlight that removing SOM input, the total input becomes more excitatory and increases the preferred size of L2/3. For small contrasts VIP cells are active (and therefore inhibit SOM) the most. Thus another factor contributing to the larger preferred size in L2/3 for small contrasts is the competition between SOM and VIP, consistent with Mossing et al. (2021).

#### SS11. MORE NUANCED PARAMETRIZATION OF THE INPUTS IN THE INVERSE STIMULUS CONDITION

In this Section we show that including more details in the parametrization of the inputs does not alter the behavior of the system significantly. In the main text we prefer a parameterization that fits the data less well but includes less parameters and has a simpler analytical expression, allowing to write the solution for the L2/3 rate field as a function of space and size in a more readable and interpretable way.

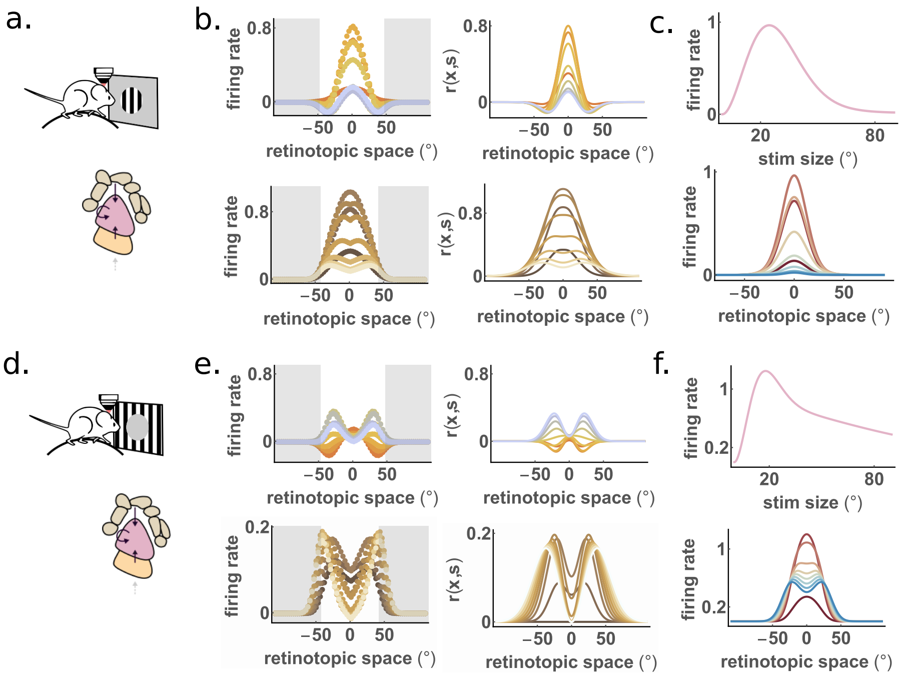


FIG. SF24. **a**. Sketch of classical stimulus and minimal model with inputs from HVAs and L4. **b**. (left) Experimental rate fields measured during classical stimuli presentation in L4 (upper panels) and HVAs (lower panels). (Right) more nuanced parametrization of the inputs. Compare with main text, Fig.2 and Fig.4. **c**. Analytic result for the size tuning curve (top) and the rate field (bottom) of Pyr+PV, using the more accurate parametrization of L4 and HVAs. **d**-**f**: same as a-c for inverse stimuli. Note that the parametrization of HVAs is different with respect to the one used in the main text, showing that the effect of inverse surround suppression is not dependent on the parametrization chosen, as long as the rate fields of HVAs share the main features with the recorded ones.

#### SS12. HVAS CELLS CLOSE TO THE EDGE OF THE INVERSE STIMULUS RESPOND MAXIMALLY

Comparing the scaling of the width of the outer Gaussian with stimulus size is useful at an analytical level. However, for the inverse stimulus, another spatial scale can be more relevant physiologically: the retinotopic location (relative to the center) where the rate field is maximal. If this quantity scaled with the stimulus radius, we could conclude that HVAs are mostly active close to the edge of the inverse stimulus. Figure SF25 shows that the scaling of this quantity is larger than the scaling of $\sigma_{HVA}$. Moreover if we consider all the sizes the data indicates that this quantity scales more weakly than the stimulus radius, but if we consider only intermediate sizes the data are consistent with the hypothesis that the HVAs cells that respond maximally are the ones whose RF is close to the edge of the 'hole'. It is tempting to speculate that this supports the involvement of HVAs in contour detection Angelucci et al. (2017). When the HVAs rate field is parameterized as a circle around the origin (see in Fig.6e in the main text), the y axis of Fig.SF25 represents the radius of that circle, $\sigma_{HVA,r}$.

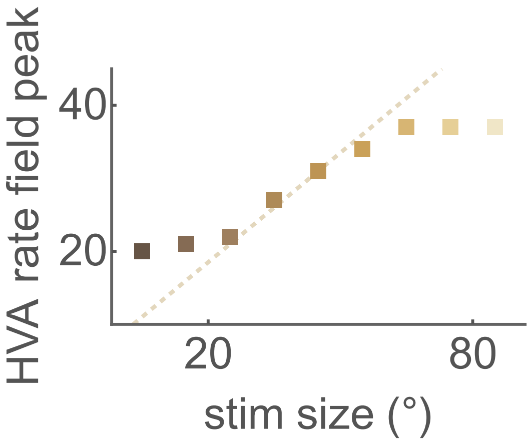


FIG. SF25. Symbols are the peak values of the experimental HVAs rate fields as depicted in Fig.SF10. The dashed line is the scaling of the radius of the stimulus $s/2$ ($s$ is the diameter of the stimulus). The peak amplitudes scale approximately as the radius of the stimulus for intermediate stimulus sizes.

### SS13. THE MINIMAL MODEL FOR INVERSE RESPONSE AND INVERSE SIZE TUNING

#### A. Optogenetic silencing of HVAs

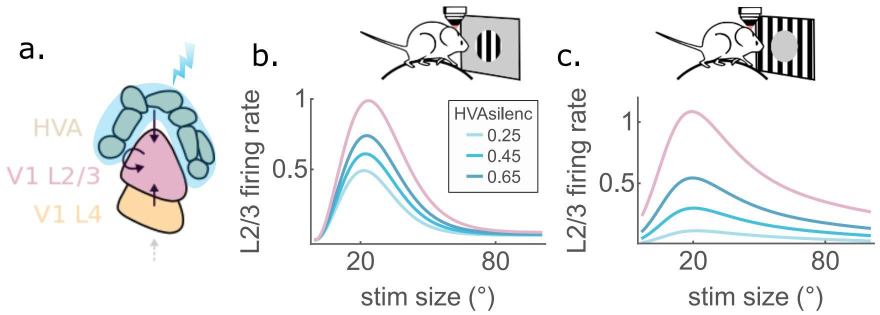


FIG. SF26. **a**. Sketch of the optogenetic silencing of HVAs. In the main text we mimic the optogenetic silencing of HVAs by reducing by a factor 0.65 the input from HVAs. Here we show how the response of L2/3 changes when we consider changes in the effectiveness of HVAs silencing, i.e. reducing by a factor (reported in the legend) the input from HVAs. **b**. and **c**. comparison of the response in the control condition (light red) and with optogenetic silencing of HVAs (light blue shades), respectively for classical (left) and inverse stimuli (right). Regardless of the effectiveness of the optogenetic silencing of HVAs, the effect on inverse response is more dramatic.

#### B. Contrast dependent inverse size tuning curves

Recordings on contrast dependent activity in L4 and HVAs during inverse stimulus presentation are not available. As a minimal assumption we can simulate reduced contrast by reducing both feedforward and feedback input by a factor of $C_0 = 0.8$ (65%contrast) and $C_0 = 0.5$ (25%contrast), as for Fig.SF22. The model reproduces the contrast dependence of the amplitude of response shown in Extended Data Fig.3h in Keller et al. (2020a). The trend of the SMI reported in Fig.SF27 is very similar to the classical case. However we do not consider this last result as a reliable prediction of the minimal model, since the behavior of the SMI in the inverse stimulus condition is not robust across parameter choices.

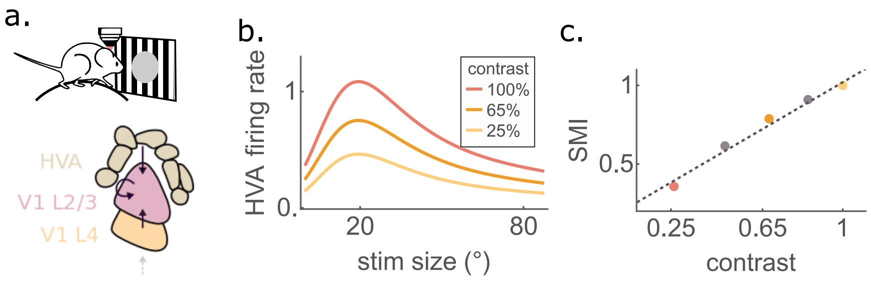


FIG. SF27. **a**. Sketch of the minimal model with the inputs considered. **b**. Comparison between the size tuning curve at maximum contrast (red) and two smaller contrasts, simulated by reducing both feedforward and feedback input as above ($C_0 = 0.8$ orange and $C_0 = 0.5$ yellow). Note that the behavior reproduces the experimental observation that inverse response decreases when contrast decreases Keller et al. (2020a). **c**. Surround Modulation Index (same plot as Fig.SF22) shows the same trend with respect to the classical response: inverse SMI increases with contrast (Extended Data Fig.3h in Keller et al. (2020a)).

#### SS14. THE ORIGIN OF INVERSE RESPONSE AND INVERSE SIZE TUNING

First of all, let us note that the agreement between analytic results and simulations for the inverse stimulus condition is less accurate than in the classical case because of the imprecision introduced by the parametrization of the inputs (see Fig.4b,c,e in the main text)and also because the ansatz itself, i.e. a Gaussian modulated by a slowly varying function. Nevertheless, in all the analyzed simulations, the agreement is fairly good, therefore for efficiency reason, we study the analytical solution. To tackle the origin of inverse response we change the value of the projection span from HVAs to L2/3 $\sigma_{L23-HVA}$ and we keep the rate field of HVAs fixed, for a fixed stimulus size. In the main text we explained that when this projection span is too small then inverse response is not allowed, meaning that the response of offset cells is larger than the response of aligned cells. This can be easily understood by looking at the input current from HVAs, which is a convolution between the connectivity and the rate field of HVAs. If the width of the connectivity is large enough, then the convolution transforms the ring profile of the rate field of HVAs into a function with a peak in the center thus generating inverse response.

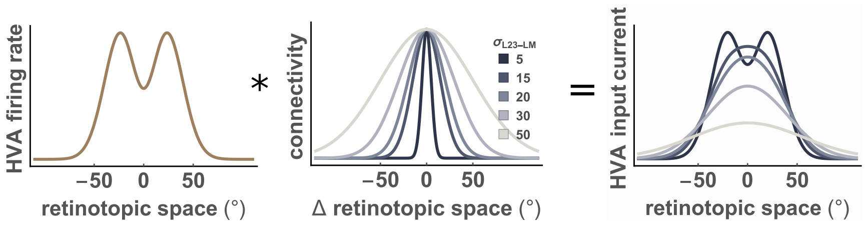


FIG. SF28. Illustration of the convolution between the HVAs rate field for stimulus size 15° and HVAs to L2/3 connectivity (normalized), with width $\sigma_{L23-HVA}$ as in the legend.

To tackle the origin of inverse size tuning in an analytical way we counterfactually change the profile width of HVAs, so that the width of the input ring (rate field of HVAs) does not grow with stimulus size. In particular here we use the parameterization of HVAs rate field with a difference of Gaussian and fix the growth of the width of both Gaussians to 0. In this case inverse surround suppression disappears. This is consistent with the results shown in Fig.6f-m of the main text, but in this case the modification leads to a complete fail of inverse surround suppression. This confirms that the origin of inverse surround suppression is the fact that for large stimuli the ring becomes too large to generate a convex function in the centre after the convolution with the connectivity. Instead if we fix the width of the ring profile, the convexity of the input current does not change.

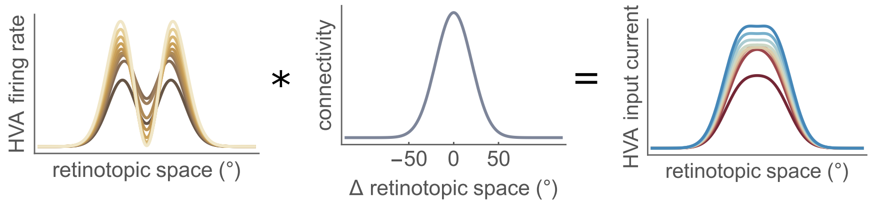


FIG. SF29. Illustration of the convolution between HVAs to L2/3 connectivity $\sigma_{L23-HVA}$ and the HVAs rate field under the counterfactual modification of null width scaling (note that mathematically, for the parameterization that we chose, if we only eliminate the growth of the width of the Gaussians with stimulus size, the values of the maxima change accordingly). Note that inverse surround suppression is absent in this case (blue curves – corresponding to large stimuli) are larger than red ones. Also note that the convexity of the rate field in the center does not change.

##### A. Analytical results for the minimal model in the inverse stimulus condition

Here we show that it is not possible to obtain inverse surround suppression if HVAs ring size does not increase with stimulus size. More precisely we show that the feedback current incoming to the cells aligned with the center of the stimulus grows indefinitely with stimulus size if HVAs ring size is fixed ($v'_{pH}(s) = 0, v'_{qH}(s) = 0$). With the parametrization chosen (see Eq.(17) and (22)in the STAR Methods), and given Eq.(27) in the STAR Methods we have:

$$
\begin{aligned}
I_H(\mathbf{x}=0,s) &= (\alpha_1^H+\alpha_2^H s)\frac{\bar{v}^i_{pH}}{\bar{v}^i_{pH}+v_{EH}} - \left[(\alpha_1^H+\alpha_2^H s)-\rho_1^H\left(\text{erf}\,\frac{s}{S^H_{\rho 1}}-\text{erf}\,\frac{s}{S^H_{\rho 2}}\right)\right]\frac{\bar{v}^i_{qH}}{\bar{v}^i_{qH}+v_{EH}}. \\
\frac{dI_H(\mathbf{x}=0,s)}{ds} &= I'_H(\mathbf{x}=0,s) = \alpha_2^H\frac{\bar{v}^i_{pH}}{\bar{v}^i_{pH}+v_{EH}} - \left[\alpha_2^H-\frac{2\rho_1^H}{\sqrt{\pi}}\left(e^{\frac{s^2}{\left(S^H_{\rho 1}\right)^2}}-e^{\frac{s^2}{\left(S^H_{\rho 2}\right)^2}}\right)\right]\frac{\bar{v}^i_{qH}}{\bar{v}^i_{qH}+v_{EH}}. \\
lim_{s\to\infty} I'_H(\mathbf{x}=0,s) &= \alpha_2^H\left(\frac{\bar{v}^i_{pH}}{\bar{v}^i_{pH}+v_{EH}}-\frac{\bar{v}^i_{pH}-1}{\bar{v}^i_{pH}-1+v_{EH}}\right) = \alpha_2^H v_{EH} \geq 0
\end{aligned}
\tag{12}
$$

Thus, under the parameterization chosen, the feedback current coming into the center in L2/3 (i.e. the location that codes for the stimulus center) keeps growing for very large sizes, i.e. there is no inverse surround suppression.

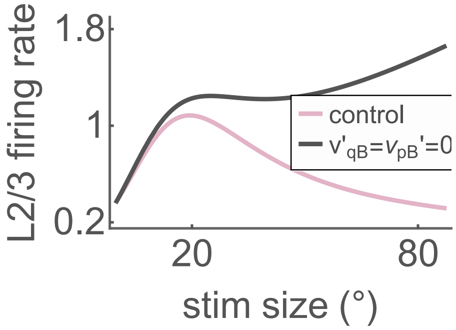


FIG. SF30. The growth of HVAs ring size is necessary condition for inverse surround suppression. Inverse size tuning curve for the control case (red) and for the counterfactual modification of fixed HVAs ring size ($v'_{pH}(s)=0, v'_{qH}(s)=0$), showing that the center response keeps increasing for large inverse stimuli.

#### SS15. A COMPUTATIONAL MODEL TRAINED TO FIT CELL-TYPE-SPECIFIC RECORDINGS FOR CLASSICAL STIMULI PREDICTS INVERSE RESPONSES

The operating regime of cortical computations emerges from the interplay of Pyramidal cells with multiple interneuron types. In mouse V1, approximately 80% of these interneurons are PV, SOM, or VIP cells (Tremblay et al., 2016; Pfeffer et al., 2013). In this Section we show that our results hold in a network model that includes all four of these cell types. We call this the 'full model' (see Fig. SF31a).

Since the synaptic coupling weights are not easily measured in the lab (e.g., see varying results in Billeh et al., 2020; Pfeffer et al., 2013; Karnani et al., 2016a), we infer them (as in Dipoppa et al., 2018; Keller et al., 2020b; Palmigiano et al., 2020; Mossing et al., 2021), together with threshold offsets specific to each cell type, by constraining the model to reproduce the recurrent rate fields recorded in the classical stimulus condition (see STAR Methods). In order to avoid a proliferation of parameters: i) we set to zero the connections that have been shown experimentally to be very small Billeh et al. (2020); Campagnola et al. (2021); Pfeffer et al. (2013); Karnani et al. (2016a,b); Fu et al. (2014); Garcia-Junco-Clemente et al. (2017); Pi et al. (2013); Jiang et al. (2015) as in Dipoppa et al. (2018); Keller et al. (2020b) and ii) we fix the connection widths based on anatomical measurements Rossi et al. (2020); Billeh et al. (2020); Karnani et al. (2016a) (see Fig. SF31b and Supplementary SS1).

In SS15 A we develop a semianalytical non-negative-least-squares approach to fit the free parameters of the model. Since all the inputs that SOM cells receive in the classical stimulus condition are surround suppressed, yet SOM cells are not, it is particularly hard to recover SOM size tuning properties. Possible solutions to this problem are to consider divisive inhibition from SOM to VIP Dipoppa et al. (2018) or additional inputs that grow with stimulus size Mossing et al. (2021). Here for simplicity we adopt the latter solution. A possible explanation is that SOM cells receive input from the surround that is stronger and/or has different orientation preference than the input from the center Marques et al. (2018); Adesnik et al. (2012); Samonds et al. (2017); Iacaruso et al. (2017). In Supplementary SS15 B we discuss the size of the extra input relative to the net other input.

We fit the model to reproduce the classical response profiles. We find a collection of parameter sets that fit the data well (for a model comparison see Supplementary SS15 C). In Fig. SF31b we plot one of the inferred parameter sets, and the resulting match of experimental and model classical size tuning curves (Fig. SF31c). With the inferred parameters, and given the external inputs from HVAs and L4 and the increase in strength of HVAs input for inverse vs. classical stimuli, the model quantitatively reproduces the inverse response and size tuning of all cell types in L2/3 (Fig. SF31d). This confirms that the inputs (in particular the input from HVAs) play a fundamental role in shaping the inverse patterns of activity, as suggested in Keller et al. (2020a) and clarified by the minimal model. Moreover, we show that the results presented here are robust against reasonable changes in the connection width, accounting for errors in the experimental estimates considered (see Supplementary SS16). In SS17 A and SS17 B we also show that the full model is stabilized by PV cells and that it recovers the modulations induced by changing the stimulus contrast, by silencing HVAs, and by suppressing SOM cells.

The full model can be leveraged to test the robustness of the mechanisms uncovered through the minimal model. In particular Fig. SF39 confirms that inverse response and size tuning in the full model require: i) broad enough feedback connections, ii) growth of the width of the HVAs rate field. Therefore the principles discussed and analysed in the main text for the minimal model extend to this more biologically realistic framework.

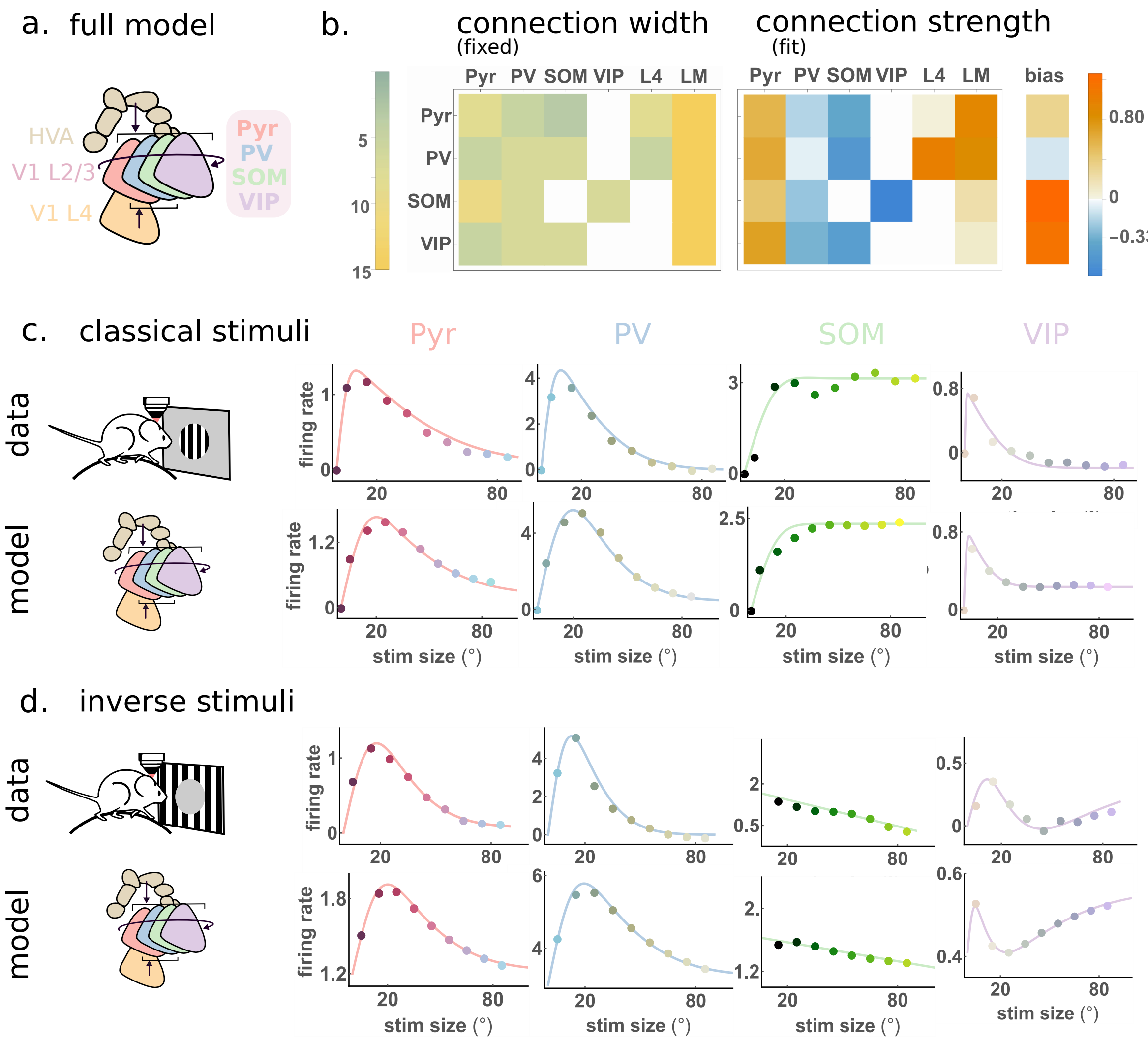


FIG. SF31. **A computational model trained to fit cell-type-specific recordings for classical stimuli predicts inverse responses a**. Sketch of the full model, with 4 recurrent cell types in L2/3 of V1 (Pyr, PV, SOM, VIP), plus feedforward input from L4 (targeting only Pyr and PV) and feedback input from HVAs (targeting all cell types). **b**. Left: a matrix of connection widths (standard deviations of Gaussian spatial functions, in degrees) estimated from the literature (see Supplementary SS1 and references therein). Right: the set of connection weights and biases, obtained by the non-negative-least-squares semi-analytic procedure (see Supplementary SS15 A). **c**. Classical size tuning curves. Comparison between experimental tuning curves (full circles top row) and steady states of the simulations of the full model (full circles bottom row) with free parameters as in **b**. The full lines are fits to the data/simulations. **d**. Same as **c** for inverse size tuning stimuli. The model with parameters inferred to fit the responses to classical stimuli generalizes to reproduce the inverse size tuning curves for all 4 cell types in L2/3.

##### A. Non-Negative-Least-Squares approach to fit the full model

In the full model the recurrent layer is composed of Pyramidal neurons (Pyr or E) as well as the 3 main interneuron classes, Parvalbumin (PV or P), Somatostatin (SOM or S) and Vaso-intestinal peptide (VIP or V), which together account for about $80-90\%$ of the neurons in mouse V1 Tremblay et al. (2016); Pfeffer et al. (2013). In addition, as in the minimal model, we consider external feedforward input from L4 (L) and external feedback input from HVAs (H). In order to keep the number of parameters as small as possible, we consider rectified quadratic transfer functions for all celltypes and we fix the projection span of each connection based on anatomical observations (see Supplementary SS1). Since the firing rates are not 0, we assume that the total input current to each population is positive. Then we can write the following equation for the rate field of population $A$ ($A \in \{E, P, S, V\}$; note that L4 and HVAs are taken as external inputs with rate fields taken from experimental measurement):

$$\sqrt{r_A(\mathbf{x})} = \sum_B \int W_{AB}(\mathbf{x}-\mathbf{y}) r_B(\mathbf{y}) d\mathbf{y} + T_A \tag{13}$$

where $B = \{E, P, S, V, L, H\}$ and $\mathbf{x}$ denotes 2 dimensional cortical or –equivalently for the model– retinotopic position. The term $T_A$ is a bias that we consider constant (across space and stimulus size), which can be understood as a cell-type-specific firing threshold.

As in the minimal model, we consider Gaussian assumptions and ansatz:

$$\begin{aligned} W_{AB}(\mathbf{x}-\mathbf{y}) &= \frac{\bar{W}_{AB}}{2\pi v_{AB}} e^{-\frac{(\mathbf{x}-\mathbf{y})^2}{2v_{AB}}} \\ r_A(\mathbf{x}) &= \bar{r}_A e^{-\frac{\mathbf{x}^2}{2v_A}} + b_A, \end{aligned} \tag{14}$$

where $b$ is the baseline, i.e. the firing rate in absence of inputs, uniform across space. Here we aim to solve for the $\bar{W}_{AB}$ given the $r_A(\mathbf{x})$ (see Supplementary SS2 B) and given the values of $v_{AB}$. In order to regress the connection weights, we set up a non negative least squares (NNLS) approach similarly to Dipoppa et al. (2018); Keller et al. (2020b). Let us consider first the NNLS procedure when we constrain only on one stimulus size. Then the loss function reads Note1:

$$\begin{aligned} S_A &= \int_{-\infty}^{\infty} d\mathbf{x} \left[ \sqrt{r_A(\mathbf{x})} - \sum_B \int W_{AB}(\mathbf{x}-\mathbf{y}) r_B(\mathbf{y}) d\mathbf{y} - T_A \right]^2 \\ &= \int_{-\infty}^{\infty} d\mathbf{x} \left[ \sqrt{\bar{r}_A e^{-\frac{\mathbf{x}^2}{2v_A}} + b_A} - \sum_B \bar{W}_{AB} \left( r_B \frac{v_B}{v_{AB+}} e^{-\frac{x^2}{2v_{AB+}}} + b_B \right) - T_A \right]^2 \end{aligned} \tag{15}$$

where we defined $v_{AB+} = v_{AB} + v_B$. It is easy to verify numerically that

$$\sqrt{r_A e^{-\frac{\mathbf{x}^2}{2v_A}} + b_A} \simeq q_A e^{-\frac{\mathbf{x}^2}{4v_A}} + \sqrt{b_A}, \tag{16}$$

is a very good approximation, where we defined $q_A = \sqrt{r_A + b_A} - \sqrt{b_A}$. The NNLS equations consist in minimizing the loss function (which codifies the error between the data and the model) with respect to the parameters. Thus when we minimize with respect to $\bar{W}_{AB}$ we obtain:

$$\begin{aligned} -2 \int_{-\infty}^{\infty} d\mathbf{x} \Bigg[ q_A e^{-\frac{\mathbf{x}^2}{4v_A}} - \sum_C \bar{W}_{AC} \left( \bar{r}_C \frac{v_C}{v_{AC+}} e^{-\frac{\mathbf{x}^2}{2v_{AC+}}} + b_C \right) + \\ + \sqrt{b_A} - T_A \Bigg] \left( \bar{r}_B \frac{v_B}{v_{AB+}} e^{-\frac{\mathbf{x}^2}{2v_{AB+}}} + b_B \right) = 0 \\ \frac{\bar{r}_B v_B 2 v_A q_A}{v_{AB+} + 2v_A} - \bar{r}_B v_B \sum_C \bar{W}_{AC} \frac{\bar{r}_C v_C}{v_{AB+} + v_{AC+}} - \bar{r}_B v_B \sum_C \bar{W}_{AC} b_C + \bar{r}_B v_B (\sqrt{b_A} - T_A) + \\ + b_B q_A 2 v_A - b_B \sum_C \bar{W}_{AC} \bar{r}_C v_C - \int_{-\infty}^{\infty} d\mathbf{x} \frac{b_B}{2\pi} \left( \sum_C W_{AC} b_C - (\sqrt{b_A} - T_A) \right) = 0 \end{aligned} \tag{17}$$

where we defined $v_{AB+} = v_{AB} + v_B/2$ Note that the second derivatives are always negative, thus the problem is always convex. Also note that these are 6 equations for each of the $\bar{W}_{AB}$ with $B = \{E, P, S, V, L, H\}$, i.e. one equation for each of the connection weights that enter into the self-consistent equation for the firing rate of population $A$. Since the space is continuous and infinite, to ensure convergence of the loss function, we have to impose:

$$\sqrt{b_A} - T_A = \sum_C \bar{W}_{AC} b_C \tag{18}$$

Conceptually this means that if we knew the $\bar{W}_{AB}$, then the knowledge of the baseline firings (and the transfer functions) would determine the firing thresholds. Moreover when we take the first derivative of the loss function with respect to $T_A$ and we take into account Eq. 18 we obtain:

$$q_A 2v_A - \sum_C \bar{W}_{AC} \bar{r}_C v_C = 0 \tag{19}$$

Finally, using both Eq. 18 and Eq. 19 in Eq. 17, we obtain

$$\sum_C (\bar{W}_{AC} \bar{r}_C v_C) \left( \frac{1}{v_{AB+} + 2v_A} - \frac{1}{v_{AC+} + v_{AB+}} \right) = 0. \tag{20}$$

We can solve this linear equation to find $\bar{W}_{AC}$ and then find $T_A$ from Eq. 18. If we solve the linear system in Eq 20 without any further caution, we find an error function that is strictly 0, but values of the parameters that are not compatible with the excitatory or inhibitory nature of the different populations. Therefore we devise an algorithm to solve this system of linear equations which constrains the positivity of each element $W_{AB}$. We start from a random initial set of $W_{AB}$ that satisfies our positivity constraints and solve one equation at a time. We update the initial parameter set if two conditions are verified: i) the solution still satisfies the positivity constraints and ii) the loss function is actually smaller than its value before this update. This simple algorithm allows us to find a family of solutions, compatible with the results from the simulation of Eq. 13 where the $\bar{W}_{AB}$ and the $T_A$ are fixed and the $\bar{r}_A$ and $b_A$ are the steady states reached by the system.

A limitation of this approach is that the additional inputs that we are explicitly ignoring (e.g. other layers of V1, synaptic plasticity, other input pathways, etc...) will be compensated for by our estimates of the recurrent couplings, thus conditioning the meaning of the effective synaptic coupling strengths.

Next, we generalize the procedure when constraining on all stimulus sizes in the classical stimulus condition. The loss function in this case reads:

$$S_A = \int ds \int d\mathbf{x} \left[ \sqrt{r_A(\mathbf{x}, s)} - \sum_{B=\{E,P,S,V,L,H\}} \int W_{AB}(\mathbf{x} - \mathbf{y}) r_B(\mathbf{y}, s) d\mathbf{y} - T_A \right]^2 \tag{21}$$

where $s$ is stimulus size and $r_A(\mathbf{x}, s) = \bar{r}_A(s) e^{-\frac{\mathbf{x}^2}{2v_A(s)}} + b_A$. We can follow all the same steps as above, until the very last one, where we cannot simplify due to the $\int ds$. We obtain:

$$\begin{aligned} \sqrt{b_A} - T_A - \sum_C \bar{W}_{AC} b_C = 0 \\ 2 \int ds \frac{v_A(s) v_B(s)}{2v_A(s) + v_{AB+}(s)} q_A(s) \bar{r}_B(s) - \sum_C \bar{W}_{AC} \int ds \frac{v_B(s) v_C(s)}{v_{AB+}(s) + v_{AC+}(s)} \bar{r}_C(s) \bar{r}_B(s) = 0 \end{aligned} \tag{22}$$

As before, we solve this numerically for $\bar{W}_{AC}$ with sign constraints.

Note that constraining on inverse size tuning curves would not introduce any conceptual complication (just more terms of the same type to calculate) but here we decide to not infer the model parameters to fit direct and inverse response, but instead we fit only the direct response and then benchmark against the ability to generate inverse size-dependent responses.

SOM cells aligned with the stimulus center respond strongly to large classical stimuli, suggesting that they receive a large excitatory current for such stimuli. Nevertheless, both L2/3 Pyramidal neurons and HVAs neurons are very surround suppressed, thus providing a much smaller excitatory current for large stimuli than for small stimuli. This suggests that SOM cells receive extra excitatory input, that increases with stimulus size. Since, to the best of our knowledge, empirical observations on the origin of such external current are still missing, in order to recover the size tuning curve and spatial profiles of SOM neurons we make a minimal assumption that they receive an extra excitatory current whose amplitude grows linearly with stimulus size. We then infer the effective strength of this interaction by adding one external field $X$ in the NNLS equations:

$$S_A = \int ds \int d\mathbf{x} \left[ \sqrt{r_A(\mathbf{x}, s)} - \sum_{B=\{E,P,S,V,L,H,X\}} \int W_{AB}(\mathbf{x}-\mathbf{y}) r_B(\mathbf{y}, s) d\mathbf{y} - T_A \right]^2 \tag{23}$$

with

$$\begin{aligned} r_X(\mathbf{y}, s) &= \sqrt{s} e^{-\frac{\mathbf{x}^2}{2v_X}} \\ W_{AX}(\mathbf{x}-\mathbf{y}) &= \delta_{AS} \frac{\bar{W}_{AX}}{2\pi v_{AX}} e^{-\frac{(\mathbf{x}-\mathbf{y})^2}{2v_{AX}}} \end{aligned} \tag{24}$$

The choice of $v_{SX}$ and $v_X$ is arbitrary and is made here following arguments of simplicity and analogy with other excitatory input currents ($v_{SX} = 8^2$, $v_X = 20^2$). We note that a different choice of these values would only weakly affect the geometry of the SOM rate field.

##### B. Inputs to SOM cells in the full model

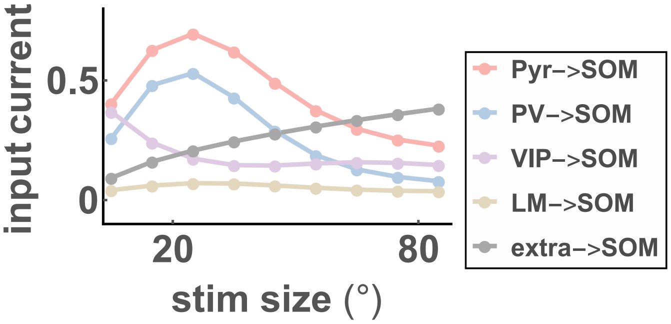


FIG. SF32. Input currents to SOM cell centered in the center of the stimulus $\mathbf{x} = 0$. The extra input here is an arbitrary function that grows with stimulus size and its strength is regressed with NNLS, as explained in Supplementary SS15 A. In absence of the "extra" input SOM would receive a smaller net input for large stimulus sizes than for small ones, thus leading to surround suppressed SOM cells. The extra input to SOM cells is introduced here in an artificial fashion, but we speculate that it could be due to feature- and position-specific connections from lateral Pyr cells or HVAs cells Marques et al. (2018); Iacaruso et al. (2017).

##### C. Model selection for the full model

We find 100 parameters sets through the NNLS procedure described above and we quantify the distance between the data and the simulated full system, by plotting:

$$\begin{aligned} \text{absolute error} &= \sum_{B=\{E,P,S,V\}} \sum_{s=5}^{85} \left(r_B(0,s) - \bar{\bar{r}}_B(0,s)\right)^2 \\ \text{normalized error} &= \sum_{B=\{E,P,S,V,L,H\}} \sum_{s=5}^{85} \left( \frac{r_B(0,s)}{\max_s r_B(0,s)} - \frac{\bar{\bar{r}}_B(0,s)}{\max_s \bar{\bar{r}}_B(0,s)} \right)^2 \end{aligned} \tag{25}$$

Here, $r_B(0, s)$ is the firing rate of cell type $B$ at position 0 to a stimulus of size $s$ (as in Eq.16 in the STAR Methods). As expected the two measures are correlated (Fig.SF33). Moreover, a fraction of the models found through the NNLS procedure diverge, which is not surprising given that the recurrent equations are not solved exactly. In what follows, as well as in the main text we present simulations with the parameter set that minimizes both the metrics above.

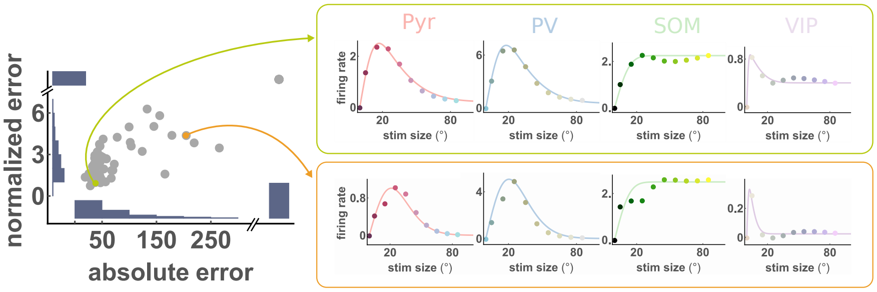


FIG. SF33. Left panel: scatter plots represents the absolute and normalized errors for the 100 models found, as defined by Eq.25. The insets are histograms showing the distribution of absolute and normalized errors, as well as the relative abundance of divergent systems. Right panels: size tuning curves for all 4 recurrent celltypes for two different parameter sets.

#### SS16. ROBUSTNESS OF THE FULL MODEL AGAINST ERRORS ON THE MEASURE OF THE PROJECTION WIDTHS

The confidence on the values of $\sigma_{AB} = \sqrt{v_{AB}}$ drawn from anatomical measures in the literature is not excellent, therefore we want to verify that the results presented here are not affected greatly by changes of these quantities (see Fig.SF34).

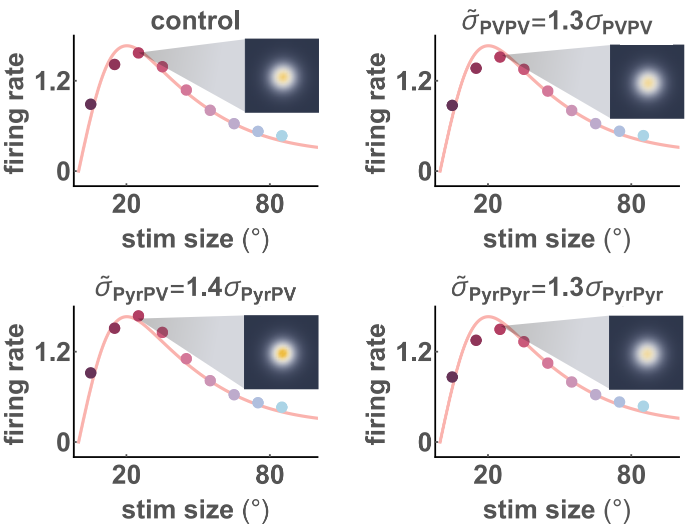


FIG. SF34. Comparison of the size tuning curve of Pyramidal neurons obtained simulating the full system with the inferred parameters using the original values of $\sigma_{AB}$ (top right) discussed in Section SS1 and using the same $\bar{W}_{AB}$ and $\sigma_{AB}$ as in the panel labels, corresponding to an increase of $30 - 40\%$ with respect to the control case. The changes are minimal both in the size tuing curves and in the rate fields widths (insets - the color corresponds to the amplitude of the rate field and xy coordinates of the plot are the two retinotopic directions). Note that the insets are all plotted with the same color code.

#### SS17. THE FULL MODEL EXPLOITS THE SAME MECHANISMS AS THE MINIMAL MODEL

##### A. PV cells stabilize the system

Previous studies Sanzeni et al. (2020); Palmigiano et al. (2020) show that PV neurons in mouse visual cortex respond paradoxically. We check that the system inferred from the experimental values of the classical rate fields is consistent with this result by freezing the response of PV cells to baseline level and verifying that the system diverges.

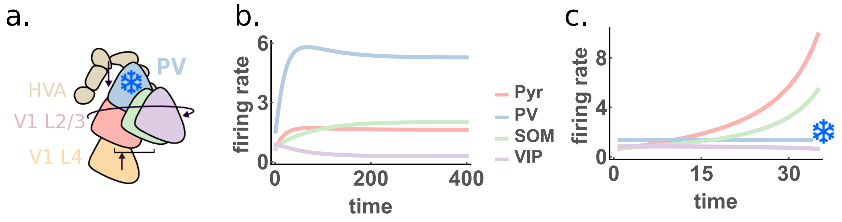


FIG. SF35. **a**. Sketch of the full system with freezing of PV cells. **b**. Time evolution of the firing rates of the 4 celltypes in L2/3 centered on the stimulus center for a stimulus of size $s = 25°$. A stable fixed point is reached. **c**. Time evolution of the system when PV cells are not allowed to update dynamically. The rate fields diverge, meaning that PV cells are responsible for stabilizing the system.

##### B. Modulations induced by changing stimulus contrast or optogenetic manipulations in the full model

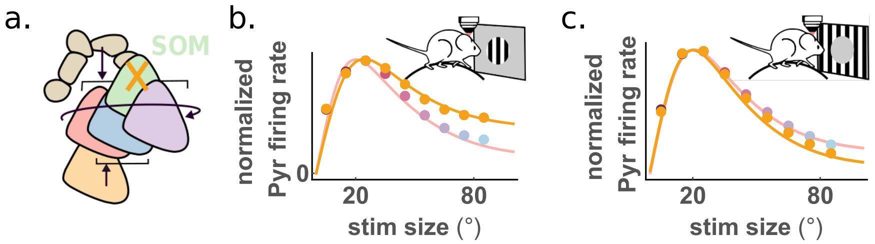


FIG. SF36. **a**. Sketch of SOM silencing in the full model. **b**. and **c**. Size tuning curves for L2/3 Pyr neurons centered on the stimulus center respectively for classical and inverse stimuli. Full circles and red curves indicate results of the simulations with SOM cells and yellow full circle and curves represent the fixed points obtained when SOM cells response is reduced, simulating hyperpolarization experiments. These results are consistent with the results shown in the main text for the minimal model, as well as with empirical observations for the classical stimuli. Hyperpolarization of SOM cells enhances inverse surround suppression.

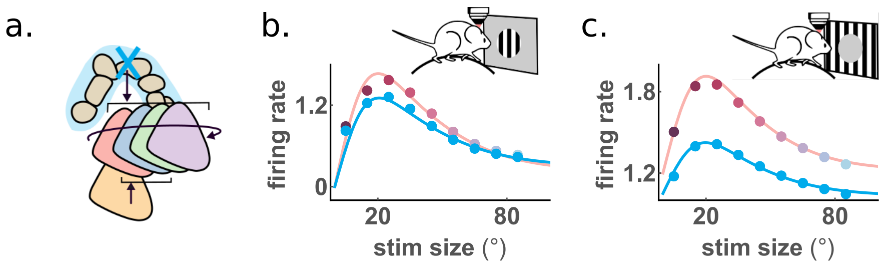


FIG. SF37. **a**. Sketch of optogenetic silencing of HVAs. Comparison between the L2/3 centered Pyr cells in the full model with inputs measured experimentally (black and red full circles) and with HVAs input reduced by a factor 0.65 (cyan circles). The results are in excellent agreement with observations during anaesthesia or optogenetic silencing of HVAs (see Fig.5 in Keller et al. (2020a)). Consistently with the experimental data, a reduction of HVAs input dramatically reduces the inverse response (panel **c**), but affects the classical response much more slightly (panel **b**). Here and in the following plots, full circles are fixed points of the simulations of the full model with parameter set reported in Fig.SF31 of the main text and full lines are their fits with differences of error functions.

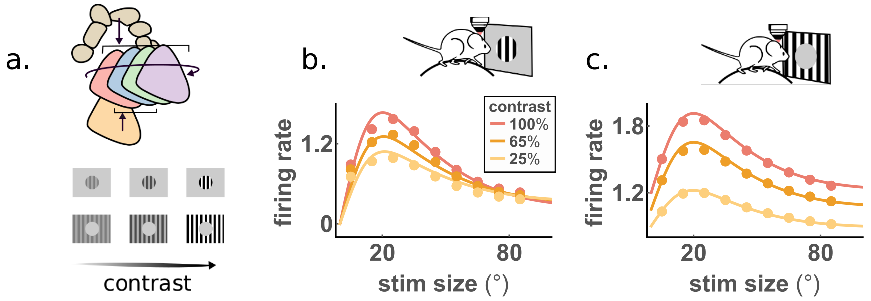


FIG. SF38. Contrast modulations on classical and inverse response of centered L2/3 Pyr cells, simulated by reducing simultaneously L4 and HVAs rate fields. Both classical (panel **b**) and inverse (panel **c**) responses are reduced when contrast is decreased. These results are consistent with the results of the minimal model and with experimental observations, although the full model does not seem to capture the increase in the preferred size when contrast decreases. As explained in the Supplementary SS10, this could be due to modulations in the activity of VIP cells or in the preferred size of the inputs with contrast that are not included in this model.

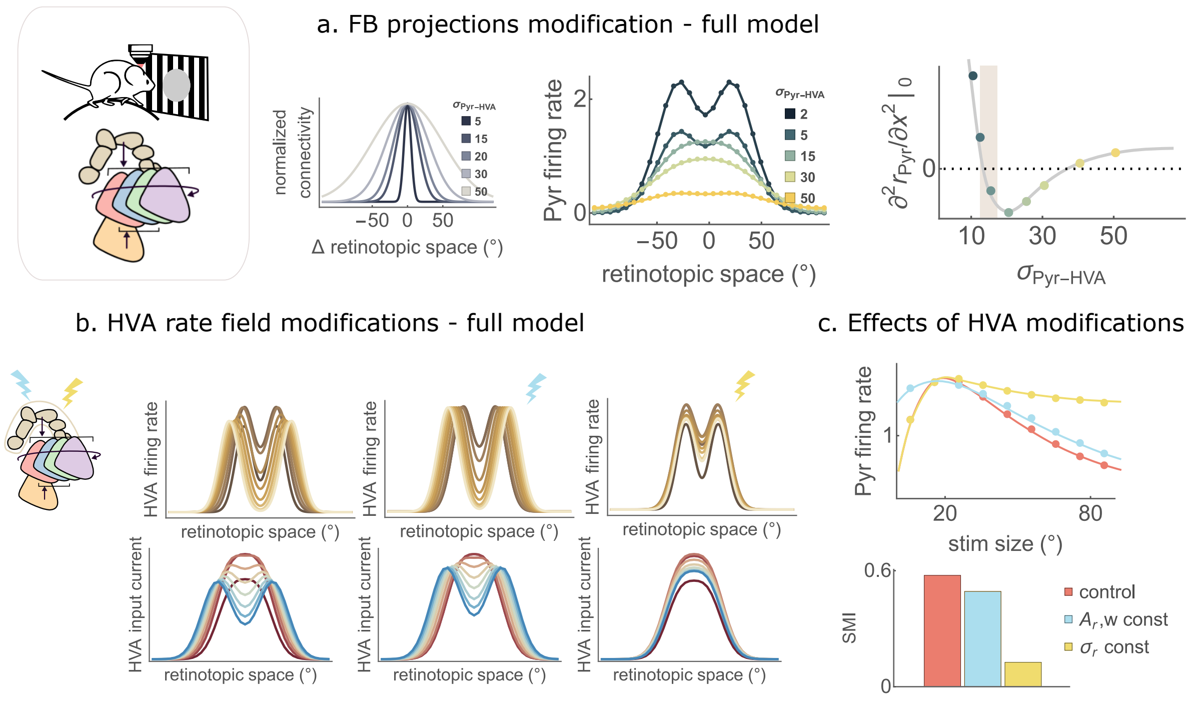


FIG. SF39. Inverse response and size tuning properties of Pyr cells in 4 cell-types model are shaped by HVAs input and require wide enough feedback projections and feedback activity profiles that scale with stimulus size. Same as Figure 6 in the main paper, but for the full model. **a**. Left: Counterfactual modification of the width of projections from HVAs to Pyr, $\sigma_{Pyr-HVA}$; here for visualization purposes the amplitudes are normalized to 1. Center: Rate field of the recurrent layer for different values of $\sigma_{Pyr-HVA}$. Inverse response depends on feedback projection's width. Right: numerical computation of the second derivative evaluated at the origin of the rate field, plotted as a function of $\sigma_{Pyr-HVA}$. The color code indicates the value of $\sigma_{Pyr-HVA}$ as in the left panel, while the shaded area corresponds to the experimental estimate in Marques et al. (2018). **b**. HVAs rate field (Top) and input current (Bottom) for the control condition (Left panels), and for two counterfactual modifications: either the size of the ring (Center panels, light blue symbol) or the amplitude and the width (Right panels, dark yellow symbol) change with stimulus size. The quantities that are kept constant are fixed at their value for $s = 15°$. **c**. (Top) Size tuning curve and (Bottom) Surround Modulation Index (see main text) for the control condition (red) and the modified ones (blue and yellow, as in **b**).

#### SS18. EXTENDING THE MINIMAL MODEL TO INCLUDE ORIENTATION TUNING

In this Section we extend the minimal model described in the STAR methods to account for orientation tuning. Given the salt-and-pepper structure of preferred orientation in mouse V1 Kaschube (2014), we assume that orientation-preference is independent of the RF location Billeh et al. (2020): in every retinotopic location all possible preferred orientations are represented. Thus we consider spatial locations and orientation as orthogonal degrees of freedom. Moreover in this Section we reduce the retinotopic space to 1D, for simplicity and without entailing any conceptual change (whereas the results presented in the main text for surround facilitation are obtained for 2 spatial dimensions plus one orientation dimension, for consistency with the other results discussed). The rate field of the recurrent layer is then $r(x,\theta)$. Let us assume that it receives a Gaussian input $I(x,\theta-\psi) = I_0 G(x,v) G_p(\theta-\psi,\lambda)$ at orientation $\psi \in (0,\pi]$, with $G_p(\theta,\lambda) = \frac{1}{\sqrt{2\pi\lambda}} \sum_{k=-\infty}^{\infty} e^{-\frac{(\theta-k\pi)^2}{2\lambda}}$ being a periodic Gaussian Persi et al. (2011) and $G(x,v) = \frac{1}{\sqrt{2\pi v}} e^{-\frac{x^2}{2v}}$. Let the recurrent connections be $W(x,y,\theta,\phi) = W_0 G(x-y, v_r) Gp(\theta-\phi,\lambda_r)$, and let the response function be rectified quadratic. We set $\sqrt{\lambda_r} = \pi/6$ based on estimates in Rossi et al. (2020). The SSN equations for the system can be written analogously to Eq(1) in the STAR Methods:

$$r(x,\theta) = \left[ I(x,\theta) + \int dy d\phi W(x,y,\theta,\phi) r(y,\phi) \right]^2 \tag{26}$$

and the exact same steps as the STAR Methods can be carried out. Obviously we will take an orientation-tuned ansatz:

$$u(x) = \alpha(x,\theta) G(x,v_u) G_p(\theta-\psi,\lambda_u) \tag{27}$$

where $\alpha(x,\theta)$ is a slowly varying function of both its arguments. All the calculations then are completely analogous to the Section on the minimal model of the STAR Methods Note2. We define $\lambda_{ru} = \lambda_r + \frac{\lambda_u}{2}$ and consider $2\lambda_u^{-1} \simeq \lambda^{-1} + \lambda_{ru}^{-1}$ then we find:

$$u(x) \simeq \frac{\pi}{W_0} \sqrt{\frac{v v_{ru} \lambda \lambda_{ru}}{\lambda_u v_u}} \left( 1 \pm \sqrt{1 - 2 I_0 W_0 \sqrt{\frac{v_u \lambda_u}{\pi v v_{ru} \lambda \lambda_{ru}}}} \right) G(x,v) G_p(\theta-\psi,\lambda) \tag{28}$$

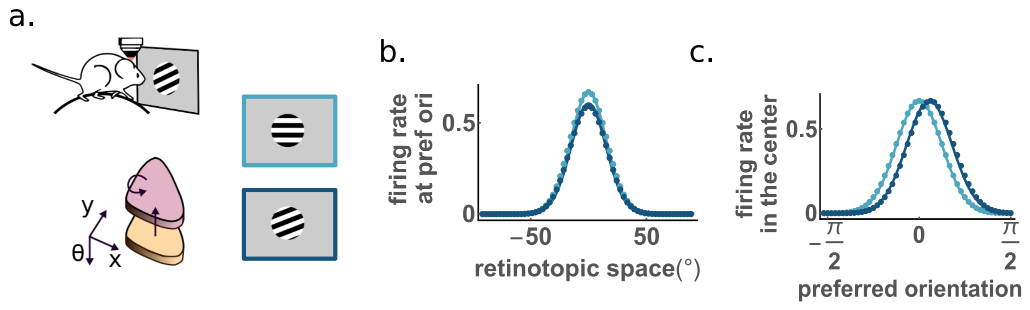


FIG. SF40. Results of the minimal model including orientation. Example of rate fields projections on the $x$ and $\theta$ axes (respectively panels **b** and **c**) for two classical stimuli of the same size but different orientation (as shown in panel **a**) More specifically panel **c** represents the response in the centre as a function of the cell's preferred orientation and panel **b** represents the response as a function of space for neurons whose preferred orientation is 0 (thus they respond slightly less to a stimulus with slightly shifted preferred orientation). Full lines are the analytic result of the minimal model and dots are the fixed points of the simulation of the 1 cell-type dynamical system.

#### SS19. LATERAL INPUT FROM SOM CELLS INCREASES SURROUND FACILITATION AND THE CONNECTION WITH SOM INVERSE RESPONSE

In Fig.SF41 we show the rate fields of SOM cells for a classical stimulus, an inverse stimulus at an orthogonal orientation and a cross stimulus, under the assumption that two rate fields of simultaneously presented stimuli at orthogonal orientations are additive (given that the populations of cells code for the two stimuli are different, see main text). We keep the stimulus size fixed to $s = 15°$. The Figure shows that SOM input current is larger in absolute value in the cross than in the classical condition. This could suggest that SOM cells suppress Pyr and PV more in the cross condition than in the classical one, therefore reducing surround facilitation. However this is not true, as shown in Fig.7 of the main text. On the contrary our model predicts that SOM cells enhance surround facilitation.

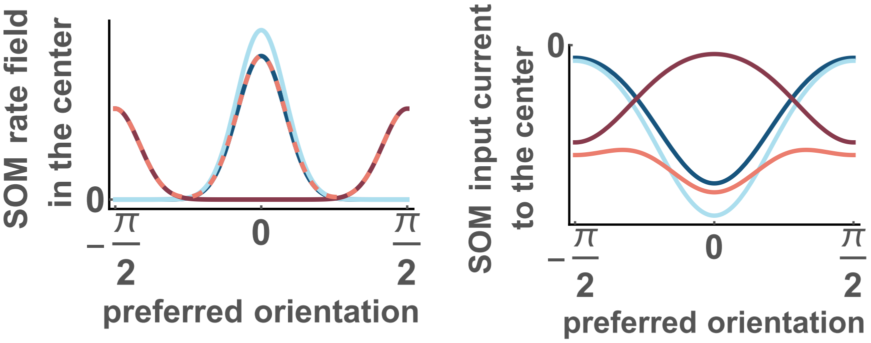


FIG. SF41. Rate field (left) and input current (right) of SOM cells in the center as a function of their preferred orientation.

This is due to the definition of surround facilitation index $SFI = \frac{r_x - r_c}{r_x + r_c}$ (based on previous choices in the literature -see e.g. $CMI$ in Keller et al. (2020b)), as opposed to $\frac{r_x}{r_c}$. Here $r_x$ (resp. $r_c$) is the firing rate of L2/3 cells centered in the stimulus and tuned to its orientation in the cross (resp classical) stimulus condition.

To better understand the role of SOM in surround facilitation we dissect the contributions to $S\tilde{F}I = \frac{u_x - u_c}{u_x + u_c}$, where $u_c$ and $u_x$ are the corresponding currents $[u_x]_+^2 = r_x$. $S\tilde{F}I$ has the same qualitative behavior as $SFI$ as shown in Fig.SF42 (compare with figure 7h in the main text)

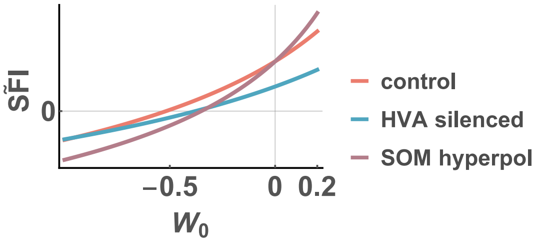


FIG. SF42. The currents surround facilitation index $S\tilde{F}I = \frac{u_x - u_c}{u_x + u_c}$ has the same qualitative behavior of $SFI = \frac{r_x - r_c}{r_x + r_c}$ (see Fig.7h in the main text).

We now define some quantities for the cross ($a = x$) or classical ($a = c$) stimulus condition:

- $u_a > 0$ is the total current incoming to Pyr+PV;
- $u_{Sa} < 0$ is the input from SOM (to Pyr+PV);
- $u_i = u_x - u_c > 0$ is the total current incoming to Pyr+PV by the effect of the surround in the cross stimulus condition;
- $u_{Si} = u_{Sx} - u_{Sc} < 0$ is the input current from SOM in the effect of the surround in the cross stimulus condition;
- $S\tilde{F}I_+$ the value of the currents surround facilitation index when SOM cells are active;
- $S\tilde{F}I_-$ the value of the currents surround facilitation index when SOM cells are silenced.

Then we have:

$$
\begin{aligned}
&\widetilde{SFI}_+ > \widetilde{SFI}_- \\
\Leftrightarrow\ & \frac{u_x - u_c}{u_x + u_c} > \frac{u_x - u_{Sx} - u_c + u_{Sc}}{u_x - u_{Sx} + u_c - u_{Sc}} \\
\Leftrightarrow\ & \frac{u_i}{u_x + u_c} > \frac{u_i - u_{Si}}{u_x - u_{Sx} + u_c - u_{Sc}} \\
\Leftrightarrow\ & u_i(u_{Sx} + u_{Sc}) < u_{Si}(u_x + u_c) \\
\Leftrightarrow\ & u_i(u_{Si} + 2u_{Sc}) < u_{Si}(u_i + 2u_c) \\
\Leftrightarrow\ & u_i u_{Sc} < u_{Si} u_c
\end{aligned}
\tag{29}
$$

Finally, since $u_{Sc} < 0$ and $u_{Si} < 0$, we have

$$
\frac{u_i}{u_c} > \frac{|u_{Si}|}{|u_{Sc}|}, \tag{30}
$$

which means that the inverse response of Pyr+PV population, normalized to its classical response, is larger than the inverse response of SOM population normalized to its classical response. In summary the surround facilitation index decreases when SOM cells are silenced because SOM cells respond less than Pyr+PV cells to the inverse stimulus (when responses are normalized to classical responses). In fact Pyr+PV cells respond more to inverse than to classical stimuli of their respective preferred size, whereas SOM cells respond more to classical than to inverse stimuli (see Fig.3n and 5d in the main text).

M. Dipoppa, A. Ranson, M. Krumin, M. Pachitariu, M. Carandini, and K. D. Harris, Neuron **98**, 602 (2018).

L. F. Rossi, K. D. Harris, and M. Carandini, Nature **588**, 648 (2020).

T. Marques, J. Nguyen, G. Fioreze, and L. Petreanu, Nature neuroscience **21**, 757 (2018).

E. Persi, D. Hansel, L. Nowak, P. Barone, and C. van Vreeswijk, PLoS computational biology **7**, e1001078 (2011).

H. Adesnik, W. Bruns, H. Taniguchi, Z. J. Huang, and M. Scanziani, Nature **490**, 226 (2012).

Y. N. Billeh, B. Cai, S. L. Gratiy, K. Dai, R. Iyer, N. W. Gouwens, R. Abbasi-Asl, X. Jia, J. H. Siegle, S. R. Olsen, et al., Neuron **106**, 388 (2020).

L. Campagnola, S. C. Seeman, T. Chartrand, L. Kim, A. Hoggarth, C. Gamlin, S. Ito, J. Trinh, P. Davoudian, C. Radaelli, et al., bioRxiv (2021).

A. J. Keller, M. M. Roth, and M. Scanziani, Nature **582**, 545 (2020a).

A. J. Keller, M. Dipoppa, M. M. Roth, M. S. Caudill, A. Ingrosso, K. D. Miller, and M. Scanziani, Neuron **108**, 1181 (2020b).

"Umap," `https://umap-learn.readthedocs.io/en/latest/faq.html`, accessed: 2022-05-09.

D. B. Rubin, S. D. Van Hooser, and K. D. Miller, Neuron **85**, 402 (2015).

D. Obeid and K. D. Miller, bioRxiv (2021), 10.1101/2020.12.30.424892, https://www.biorxiv.org/content/early/2021/01/02/2020.12.30.4248

A. Angelucci, M. Bijanzadeh, L. Nurminen, F. Federer, S. Merlin, and P. C. Bressloff, Annual review of neuroscience **40**, 425 (2017).

Y. Li and L.-S. Young, PLoS computational biology **17**, e1008916 (2021).

J. H. Siegle, X. Jia, S. Durand, S. Gale, C. Bennett, N. Graddis, G. Heller, T. K. Ramirez, H. Choi, J. A. Luviano, P. A. Groblewski, R. Ahmed, A. Arkhipov, A. Bernard, Y. N. Billeh, D. Brown, M. A. Buice, N. Cain, S. Caldejon, L. Casal, A. Cho, M. Chvilicek, T. C. Cox, K. Dai, D. J. Denman, S. E. J. de Vries, R. Dietzman, L. Esposito, C. Farrell, D. Feng, J. Galbraith, M. Garrett, E. C. Gelfand, N. Hancock, J. A. Harris, R. Howard, B. Hu, R. Hytnen, R. Iyer, E. Jessett, K. Johnson, I. Kato, J. Kiggins, S. Lambert, J. Lecoq, P. Ledochowitsch, J. H. Lee, A. Leon, Y. Li, E. Liang, F. Long, K. Mace, J. Melchior, D. Millman, T. Mollenkopf, C. Nayan, L. Ng, K. Ngo, T. Nguyen, P. R. Nicovich, K. North, G. K. Ocker, D. Ollerenshaw, M. Oliver, M. Pachitariu, J. Perkins, M. Reding, D. Reid, M. Robertson, K. Ronellenfitch, S. Seid, C. Slaughterbeck, M. Stoecklin, D. Sullivan, B. Sutton, J. Swapp, C. Thompson, K. Turner, W. Wakeman, J. D. Whitesell, D. Williams, A. Williford, R. Young, H. Zeng, S. Naylor, J. W. Phillips, R. C. Reid, S. Mihalas, S. R. Olsen, and C. Koch, Nature **592**, 86 (2021).

R. Tremblay, S. Lee, and B. Rudy, Neuron **91**, 260 (2016).

S. Denève and C. K. Machens, Nature Neuroscience **19**, 375 (2016).

A. Sanzeni, B. Akitake, H. C. Goldbach, C. E. Leedy, N. Brunel, and M. H. Histed, Elife **9**, e54875 (2020).

A. Palmigiano, F. Fumarola, D. P. Mossing, N. Kraynyukova, H. Adesnik, and K. D. Miller, bioRxiv (2020), 10.1101/2020.11.11.378729, https://www.biorxiv.org/content/early/2020/11/11/2020.11.11.378729.full.pdf.

J. S. Isaacson and M. Scanziani, Neuron **72**, 231 (2011).

M. Okun and I. Lampl, Nature neuroscience **11**, 535 (2008).

H. Bos, A.-M. Oswald, and B. Doiron, bioRxiv , 2020 (2020).

S. N. Chettih and C. D. Harvey, Nature **567**, 334 (2019).

H. Nienborg, A. Hasenstaub, I. Nauhaus, H. Taniguchi, Z. J. Huang, and E. M. Callaway, Journal of Neuroscience **33**, 11145 (2013).

Y. Ahmadian and K. D. Miller, Neuron (2021).

D. P. Mossing, J. Veit, A. Palmigiano, K. D. Miller, and H. Adesnik, bioRxiv (2021), 10.1101/2021.03.31.437953, https://www.biorxiv.org/content/early/2021/03/31/2021.03.31.437953.full.pdf.

C. K. Pfeffer, M. Xue, M. He, Z. J. Huang, and M. Scanziani, Nature neuroscience **16**, 1068 (2013).

M. M. Karnani, J. Jackson, I. Ayzenshtat, J. Tucciarone, K. Manoocheri, W. G. Snider, and R. Yuste, Neuron **90**, 86 (2016a).

M. M. Karnani, J. Jackson, I. Ayzenshtat, A. H. Sichani, K. Manoocheri, S. Kim, and R. Yuste, Journal of neuroscience **36**, 3471 (2016b).

Y. Fu, J. M. Tucciarone, J. S. Espinosa, N. Sheng, D. P. Darcy, R. A. Nicoll, Z. J. Huang, and M. P. Stryker, Cell **156**, 1139 (2014).

P. Garcia-Junco-Clemente, T. Ikrar, E. Tring, X. Xu, D. L. Ringach, and J. T. Trachtenberg, Nature neuroscience **20**, 389 (2017).

H.-J. Pi, B. Hangya, D. Kvitsiani, J. I. Sanders, Z. J. Huang, and A. Kepecs, Nature **503**, 521 (2013).

X. Jiang, S. Shen, C. R. Cadwell, P. Berens, F. Sinz, A. S. Ecker, S. Patel, and A. S. Tolias, Science **350** (2015).

J. M. Samonds, B. D. Feese, T. S. Lee, and S. J. Kuhlman, Journal of neurophysiology **118**, 3282 (2017).

M. F. Iacaruso, I. T. Gasler, and S. B. Hofer, Nature **547**, 449 (2017).


We make repeated use of the useful integrals:

$$\begin{aligned} \int e^{-\frac{(\mathbf{x}-\mathbf{y})^2}{2v_1}} e^{-\frac{\mathbf{y}^2}{2v_2}} d\mathbf{y} &= \frac{2\pi v_1 v_2}{v_1+v_2} e^{-\frac{\mathbf{x}^2}{2(v_1+v_2)}} \\ \int e^{-\frac{\mathbf{x}^2}{2v_1}} e^{-\frac{\mathbf{x}^2}{2v_2}} d\mathbf{x} &= \frac{2\pi v_1 v_2}{v_1+v_2} \end{aligned} \tag{31}$$

.


M. Kaschube, Current opinion in neurobiology **24**, 95 (2014).

The convolution of two periodic Gaussians is a periodic Gaussian whose variance is the sum of their variances. Also, note that in the case of periodic Gaussians, the equivalent of Eq.(6) in the STAR Methods holds only approximately, but it is a very good approximation, when $\lambda$ is sufficiently small compared to $\pi$.
